# *Streptomyces* Autoregulator Biosensors from Natural Product Cluster-Situated Regulators

**DOI:** 10.1101/2025.09.05.673737

**Authors:** Lauren E. Wilbanks, Carson B. Roberts, Manuela Frias-Gomez, Haylie E. Hennigan, Kylie G. Castator, Zach L. Budimir, Caroline Zu, Elizabeth I. Parkinson

## Abstract

The soil dwelling bacteria *Streptomyces* is an abundant producer of numerous anticancer, antifungal, and antibiotic compounds (i.e. Natural Products, NPs). The sophisticated cellular machinery required to produce NPs is frequently regulated by quorum-sensing systems, consisting of cluster situated regulators (CSRs), such as TetR-like repressors, and small-molecule autoregulator (AR) ligands. Only a small fraction of bioinformatically predicted quorum-sensing AR circuits have been experimentally determined, and fewer still have been engineered as inducible expression systems for synthetic biology. This research details the development of eight CSR-based AR biosensors and the synthetic routes to their AR ligands. Overall, the AR biosensors exhibit a range of maximum activation, AR affinity, and AR selectivity. We examined crosstalk between noncognate CSRs and ARs, as well as the ability of CSRs to regulate alternative operators. Additionally, we establish these biosensors can be cocultured with *Streptomyces* for rapid analysis of AR production. Finally, we demonstrate the CSR-based biosensor vectors can be combined to create orthogonal signaling systems in bacterial coculturing or multi-input genetic circuits. Longterm, these *Streptomyces* AR biosensors will contribute to the elucidation of small molecule quorum sensing circuits employed by *Streptomyces* as well as increasing the complexity of genetic circuits used in industrial or agricultural settings.

## Introduction

Signaling systems that utilize quorum sensing are widespread throughout bacteria, regulating diverse processes such as virulence, sporulation, and fluorescence ^1–5^. These signaling processes operate by using genetic elements to control gene flux in response to a stimuli (i.e. transcription factors, operators, promoters). Synthetic biologists have identified and quantified these parts for implementation into engineered constructs capable of sensing a defined input (chemicals, light, pH etc.) to trigger a change in gene expression ^6–10^. Much focus has been paid to acyl-homoserine lactones (AHLs) biosensors for designing inducible genetic circuitry for use in synthetic biology^11–17^. While AHL-based circuits demonstrate extensive utility, they can be limited by crosstalk between molecules and transcription factors^11,14^. Transcriptional repressors like TetR, LacI, λ CI, and AraC have also been extensively used for designing multi-logic gates, toggle switches, oscillators, or cellular memory, but some combinatorial logic gates can suffer from ligand-induced circuit failure^18–27^. To induce multiple specific or synchronized behaviors across many bacteria, and potentially many genera, would require a diverse set of signaling molecules and regulators, broadly employable by many organisms.

*Streptomyces*, a bacterial genus in the Actinomycetota phylum, is enriched in therapeutically relevant Natural Products (NPs). The enzymatic machinery required to biosynthesize NPs are frequently colocalized on bacterial genomes as biosynthetic gene clusters (BGCs). Production of NPs is tightly controlled by sophisticated regulatory networks that often include quorum sensing^28–32^. NP quorum sensing circuits regularly utilize TetR-like repressors encoded within the BGC (a cluster situated regulator, CSR) to constitutively suppress NP gene transcription in the absence of their cognate signaling autoregulators (ARs), resulting in silent BGCs ^1,33–36^. *Streptomyces* employ a wide variety of ARs responsible for regulating NP production, with the best studied classes including gamma-butyrolactones (GBLs) like the *Streptomyces coelicolor* butanolides (SCBs), butenolides (BNs) such as avenolide, and furans like the methylenomycin furans (MMFs) (**Fig. 1A**)^31,37^. Despite the fact that thousands of TetR-like repressor/AR quorum sensing systems are bioinformatically predicted in *Streptomyces,* as well as other Actinomycetota, only 10 CSR/AR pairs have been experimentally confirmed^37–40^. Previous research has relied on relatively low throughput and semiquantitative assays to identify CSR repressors and their ARs^41–47^. Using modern molecular techniques, researchers have assayed known *Streptomyces* CSRs for their capabilities as repressor proteins^18^. Negative inducible systems have also been developed, wherein assays utilizing the model gamma butyrolactone-binding CSR ‘ScbR’ quantified different responses to derivatives of its ARs, the SCBs^48–50^. However, these modern methods have not been applied to discovery of Actinomycetota ARs or adopted widely for use in genetic circuits.

**Figure 1:**
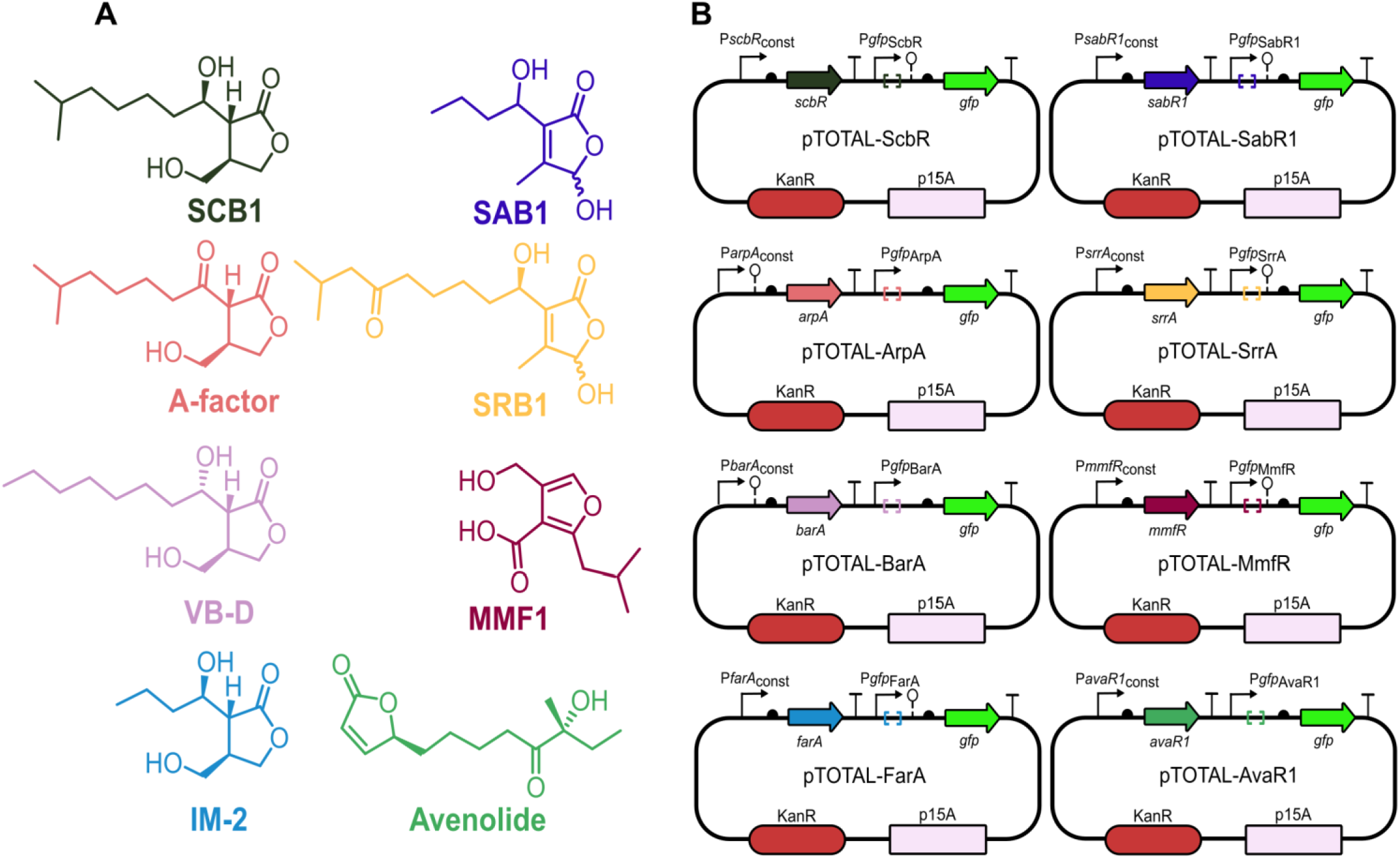
*Streptomyces* CSR-based AR biosensors and their cognate quorum sensing ARs. **A)** A subset of *Streptomyces* ARs; they are color coded according to their cognate CSR biosensors in part B. **B)** Plasmid maps of CSR-based AR biosensor vectors used for the AR induction assay. These vectors are referred to as ‘pTOTAL’ with the CSR gene name. More details of the plasmid design can be found in **Supplementary Figure 1**.

The chemical diversity of *Streptomyces* NP quorum sensing circuits, as well as their ligand/CSR specificity, makes these components excellent candidates for the development of a suite of inducible CSR-based genetic circuits ^48,50^. In this work, we introduce eight novel vectors to the synthetic biology toolkit and detail CSR/AR interactions. We also employ these vector-based biosensors in coculture with *Streptomyces* to ascertain their viability as qualitative and quantitative indicators of the presence of NP-regulating ARs. Finally, we quantify AR orthogonality in microbial coculture and design a dual input AND gate using two CSRs.

## Results

### Generation of AR biosensor vectors

Building on our previously published ScbR biosensor, we modified the origin of replication from high copy pMB1 to low copy p15A to reduce the burden of plasmid replication^50,51^. To allow for high levels of GFP production, the promoter controlling GFP expression was changed from a *Streptomyces* derived promoter to BBa_23100 (Registry of Standard Biological Parts)^52,53^. After these modifications, we generated eight vectors, each constitutively expressing a *Streptomyces* CSR repressor: ScbR, ArpA, BarA, FarA, SabR1, SrrA, MmfR and AvaR1 (**Fig. 1B** and **Supplementary Fig. 1**). Operator sequences for each repressor were identified from previous literature and were cloned downstream of the *gfp* -35 and -10 sequences, affording control of GFP expression by the CSR repressor^41,43–45,48,54–56^. The genetic insulator BydvJ was added to the *gfp* promoter region of each vector to normalize the mRNA 5’ UTR^57–59^. The presence of BydvJ increased both the baseline GFP expression and the fluorescence maxima, affording large potential induced fold changes for most of the vectors. For AvaR1, ArpA, and BarA vectors, the increase in baseline was not matched by an increase in maxima, therefore the overall fold change was reduced. For this reason, the BydvJ sequence was removed from the *gfp* promoter of these three vectors. A promoter scan was employed to determine the level of constitutive repressor expression that balanced the burden of repressor protein expression with the level required to achieve sufficient repression of GFP ^18,60,61^. Natural variability in fluorescence baseline and culture growth was observed for each pTOTAL-CSR vector, likely the result of a combination of factors, including CSR expression levels, strength of binding between repressor and operator, and inherent changes in RNA polymerase transcription due to the operator sequences (**Supplementary Table 1** and **2**)^62^.

### Chemical synthesis and testing of ARs with cognate repressors

To generate inducible genetic circuits, synthetic standards of seven *Streptomyces* quorum sensing ARs were generated (**Fig. 1A**). Previously, we have developed enantioselective routes to SCB1, A-factor, VB-D, MMF1, and SRB-1, which are native ARs for ScbR, ArpA, BarA, MmfR, SrrA, respectively^50,63–65^. Herein, we have also developed routes to the natural ligands IM-2 and SAB1. To obtain IM-2, the natural ligand for FarA, an aldol addition of protected (*R*)-paraconyl alcohol (**9**) and butanal was modified from that reported by Appayee and coworkers, followed by silyl deprotection with tetrabutylammonium fluoride (TBAF) (**Scheme 1A**)^66,67^.

**Scheme 1:**
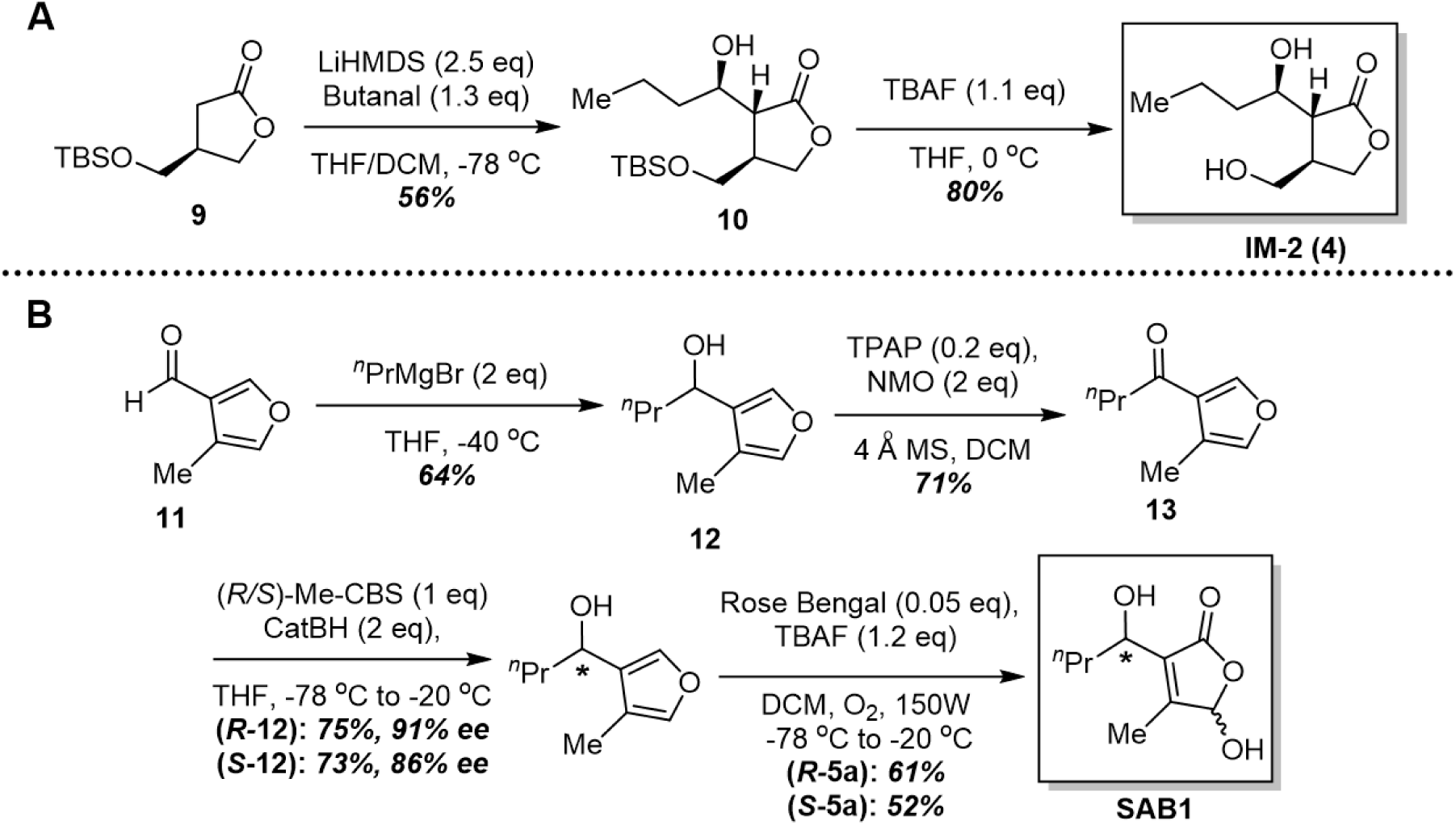
Synthetic routes for *Streptomyces* ARs. **A)** Synthesis of IM-2 (**4**)**. B)** Synthesis of (*R*)- and (*S*)- SAB1 (**5a**).

For SAB1, the natural ligand for SabR1, the original isolation did not deduce the stereochemistry at C1^55^. We hypothesized that we could synthesize both epimers at this position and test them using the GFP assay with SabR1 to determine the stereochemistry at this position. To access SAB1, the known furaldehyde **11** was alkylated with *n*-propylmagnesium bromide to afford the secondary alcohol **12**^55^. To obtain the *R* and *S* epimers, the secondary alcohol was subjected to Ley oxidation followed by Cory-Bakshi-Shibata reduction with both the *R* and *S* oxazaborolidine catalysts and catechol borane as the hydride source, in high enantiopurity (91% ee for (*R*)-**12** and 86% ee for *(S*)-**12**). The enantiomers were then subjected to photo-oxidation to afford (*R*)-SAB1 (21% overall yield) and (*S*)-SAB1 (17% overall yield). Further discussion of the synthesis can be found in **Supplementary Methods**. When subjected to the induction assay with the pTOTAL-SabR1 biosensor, the (*S)-* SAB1 was found to have a 10-fold lower K_D_ than (*R*)-SAB1, strongly suggesting that the natural configuration at C1’ is (*S*) (**Supplementary Fig. 2**). Initially, we hypothesized that the configuration at C1’ would be (*R*) as seen in the SRBs but instead the configuration appears to match that of the γ-hydroxybutenolides isolated from *Streptomyces antibioticus* Tü 99^68,69^. This is interesting as it could indicate the potential for cross talk between the two species. Given that the activity of the *(S)*-SAB and the racemic mixture were very similar, racemic SAB1 was used for the rest of the studies described herein.

### Activity of ARs for cognate and non-cognate CSRs

Each ‘pTOTAL’ CSR vector (referred to onwards by the CSR gene name) was initially evaluated with a gradient of its cognate AR: ScbR/SCB1, ArpA/A-factor, BarA/VB-D, FarA/IM-2, SabR1/SAB1, SrrA/SRB1, and MmfR/MMF1. Final dilution gradients were designed to centralize the K_D_ of each molecule within the dilution curve. (**Fig 2** and **Supplementary Table 1**). K_D_s varied from 30 nM for FarA/IM-2 to 111 µM for MmfR/MMF1. While these values differ from the previously reported values, previous researchers typically utilized AR synthase knockout strains of *Streptomyces* supplemented with synthesized or purified AR to determine the minimum effective concentration necessary to produce NPs, not the K_D_s (**Supplementary Table 3**). We posit a reasonable minimum effective concentration derived from the results of the induction assay would be the concentration at which each AR induced a fluorescence signal >2.5-fold over baseline. Under this assumption, the minimum effective concentration from the induction assay is in the hundred nanomolar range for most biosensors, with notable exceptions of FarA exhibiting higher sensitivity (3 nM) to IM-2 and ScbR/MmfR showing less sensitivity (2 μM) to their respective ARs, SCB1 and MMF1 (**Supplementary Table 1**). This median value is in line with what other techniques have shown, with the caveat that previous protocols to determine minimum effective concentration have used multiple *Streptomyces* genetic variants, AR derivatives, and protocols (**Supplementary Table 3)**.

**Figure 2:**
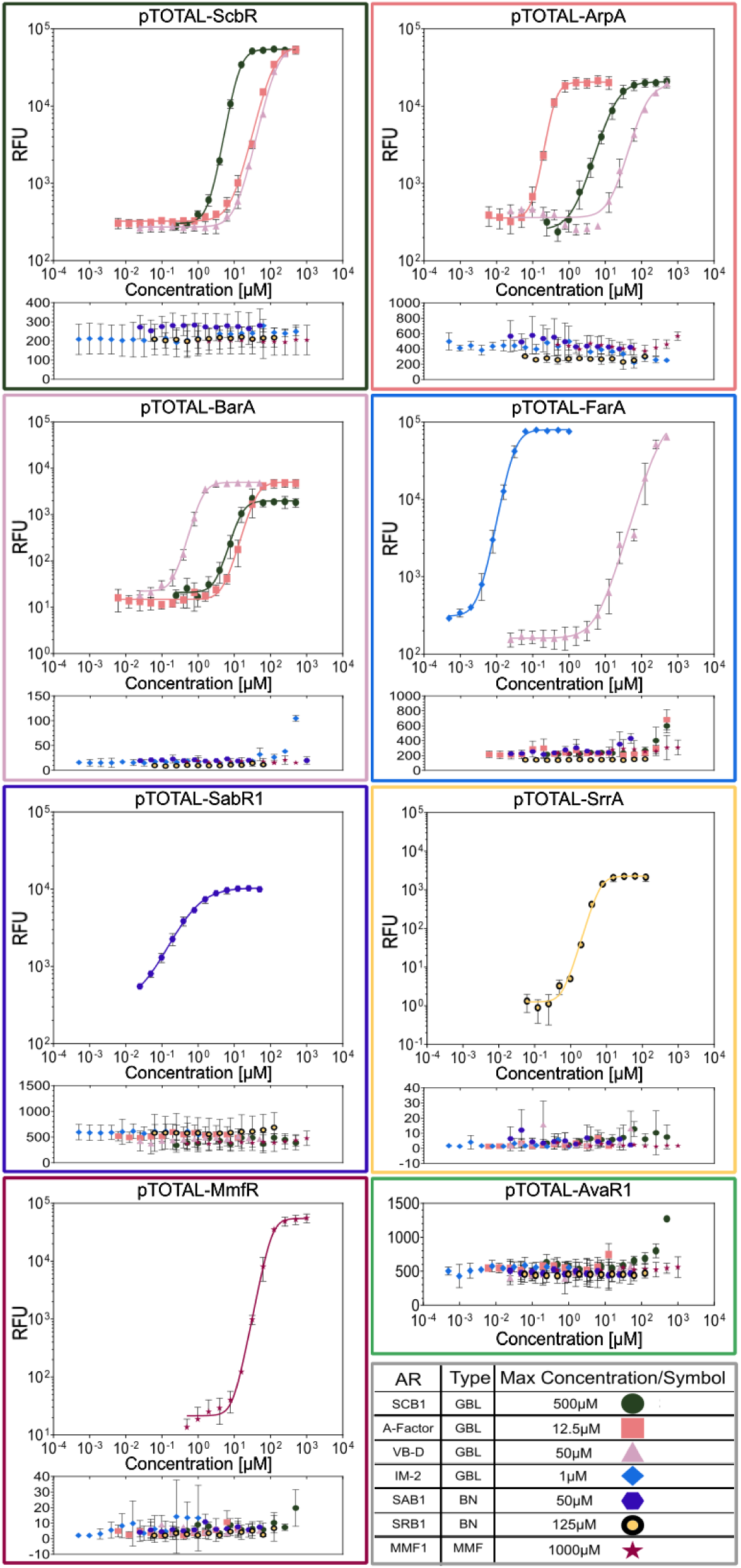
Results from fluorescence induction assays with eight CSR biosensor vectors. Graph title indicates the CSR biosensor vector that was tested, cognate molecule/CSR share a color. Molecule dilutions begin at max concentration indicated in the key, with each subsequent point a 1:2 dilution. Response is recorded as Relative Fluorescence Units (RFU). Each point is an average of three biological replicates analyzed on different days. Error bars +/- 1 SD. ‘Type’ denotes the class of *Streptomyces* AR.

After validating the induction concentrations of AR biosensors with their cognate ARs, all biosensor vectors were induced with every synthesized AR (**Fig. 2**). ScbR, ArpA, BarA, and FarA are among the best documented members of the gamma-butyrolactone receptor (GBLR) family of CSRs. The CSRs share high amino acid sequence similarity and the GBL ARs (SCBs, A-factor, VBs, and IM-2) all possess a similar lactone core (**Fig. 1A**, **Supplementary Fig. 4**, **Supplementary Fig. 5**). Previous research has noted AR crosstalk between GBLRs and GBLs^70^. As constructed, each biosensor vector exhibited different baseline fluorescence, K_D_, fluorescence maximum, and fold changes. FarA possessed both the highest fold-change of the four at 396.4- fold when induced with IM-2, as well as the most potent CSR/AR affinity with a K_D_ of 30 nM. In contrast, ArpA exhibited a high baseline and low maximum RFU, and thus had a comparatively weaker fold change of 53.7-fold with its cognate AR A-factor. ScbR exhibited a strong fold-change of 235.5 with SCB1 but demonstrated the lowest cognate molecule affinity of the GBLRs with a K_D_ of 13.3 μM (**Fig. 2**, **Supplementary Table 1**). BarA, potentially as result of not possessing a genetic insulator in the *gfp* promoter, showed the lowest induced fluorescence maximum of ∼5,000 RFU; however, the biosensor also exhibited a very low baseline fluorescence (∼21 RFU), and therefore had a maximum induced fold-change of 240.3 with VB-D (**Supplementary Table 1**). GBL crosstalk between ScbR, ArpA, and BarA indicate ARs with similar structures can overcome low CSR/AR affinity at high concentrations. However, these molecules do not achieve the same affinity of binding of a cognate AR, as exhibited by their increased K_D_ (**Fig. 2**, **Supplementary Table 1**). For example the K_D_ for ScbR induced with A-factor or VB-D is 117.3 μM and 146.3 μM, respectively, indicating ScbR is approximately 10x less sensitive to these molecules than its cognate AR SCB1 (**Supplementary Table 1**).

FarA demonstrates the most unique results of the GBLRs. Although its cognate AR IM-2 shares a similar core structure to other GBLs (*i.e.* SCB1, A-factor, and VB-D), it has a much shorter acyl chain (**Fig. 1A**). While IM-2 is a very potent AR of its cognate CSR FarA, the only other activity it exhibited was a low level of activation of BarA (∼5-fold induction). Even at the very high concentration of 500 µM, IM-2 does not derepress ScbR or ArpA; compared to its K_D_ of 30 nM, FarA shows an impressive >16,000 fold selectivity for its cognate AR IM-2 (**Fig. 2**). In testing FarA for crosstalk with other ARs, linear VB-D is the only other AR capable of its derepression, although it has a >11,000-fold decrease in potency when comparing the K_D_ of both molecules. This indicates the linear acyl chain may be more amenable to binding FarA compared to the branched chain of SCB1 and A-factor (**Fig. 1A**, **Fig. 2**). Molecular docking was employed to examine ligand/CSRs interactions as a potential tool to distinguish cognate pairs *in silico*. SCB1, A-factor, and IM-2 were each docked into AlphaFold3 models of ScbR, ArpA, and FarA^71^. Docking software scored IM-2 as the best fit for all three CSRs, contradicting our empirical data (**Supplementary Fig. 6**). The docking scores of IM-2 may be explained by the short acyl chain allowing for easier access to binding pockets of variable sizes. However, even if IM-2 can easily access the CSR binding pocket, this does not necessarily result in the ligand-amino acid interactions and subsequent change in CSR conformation required for biological activity. We assert molecular docking should not be used to unilaterally rank ligands and would caution against using docking scores as a predictor of biologically active CSR-ligand interactions.

All of the CSR-based biosensors tested, save SabR1, demonstrated ultrasensitive responses to their cognate ARs, with Hill coefficients of >2.5 (**Supplementary Table 1**)^72,73^. The SabR1 vector has a Hill coefficient of 1.1, right at the boundary of cooperativity, and has the smallest fold change over baseline of all tested vectors at 29.2 (**Supplementary Table 1**). The source of this CSR vector’s relative poor performance is unknown; results may indicate high toxicity of SabR1, improper protein folding and dimerizing, poor binding to the operator sequence, or off-target effects in the cell (**Supplementary Table 2**, **Supplementary Fig. 3**)^74^. SAB1 did not activate any CSRs other than SabR1 at the highest concentration tested (50 µM) (**Fig. 2**). Considering SAB1 is a BN-type AR, it is unsurprising to see lack of derepression for the GBLs and MMFs. SRB1, which has the same BN core but a longer and more complex alkyl chain, has no effect on the SabR1 CSR up to the highest concentration tested, 125 µM (**Fig. 2**). MMF1 with its native CSR MmfR had a large 4.91×10^3^ fold induction (**Supplementary Table 1**). This fold change is a result of a very low baseline fluorescence and a high induction maximum; however, MmfR also demonstrated the weakest binding affinity of any of the native CSR/AR pairs with a K_D_ of 111 µM (**Supplementary Table 1**). MMF1 did not activate any of the other vectors even at the high concentration of 1 mM (**Fig. 2**). This is unsurprising given the structural uniqueness of MMF1 compared to the other ligands tested (**Fig. 1A**). It is important to note the MmfR/MMF quorum sensing circuit and the ScbR/SCB quorum sensing circuit are both present in the genome of the wild-type *Streptomyces coelicolor* A3(2)^75^. Although only one derivative of each AR was tested in the synthesized molecule induction assay, no cross-activation was observed, even at their respective maximum concentrations. These results suggest that chemically diverse quorum sensing AR circuits may be one way that *Streptomyces* has evolved orthogonal NP regulatory systems.

Treatment of SrrA with the cognate SRB1 ligand shows a 1.04×10^3^ maximum fold change and a K_D_ of 6.8 µM (**Supplementary Table 1**). SRB1 did not induce any significant change (>2.5-fold change in RFU) with any of the other repressors at the highest concentration tested (125 µM) (**Fig. 2**). Additionally, none of the other molecules induced a response from the SrrA CSR. Interestingly, SAB1 had no effect on SrrA at the highest concentration tested (50 µM). This is likely because SRB ARs possesses a longer acyl chain compared to the SABs (C9-10 for SRBs vs C3-4 for SABs). (**Fig. 1A**) The results for SabR1 and SrrA are analogous to those observed with the GBLs and their receptors and reiterate the importance of the acyl chain length in the specificity of the ligands. We were unable to successfully synthesize avenolide to explore the AvaR1 biosensor vector with its cognate AR (**Fig. 1A**). AvaR1 showed no activation against any of the ARs tested, indicating strong specificity for its cognate butenolide (**Fig. 2**). However, we were able to confirm the activation of the AvaR1 vector using crude extracts from the avenolide producing strain *S. avermitilis* (**Fig. 3**).

**Figure 3:**
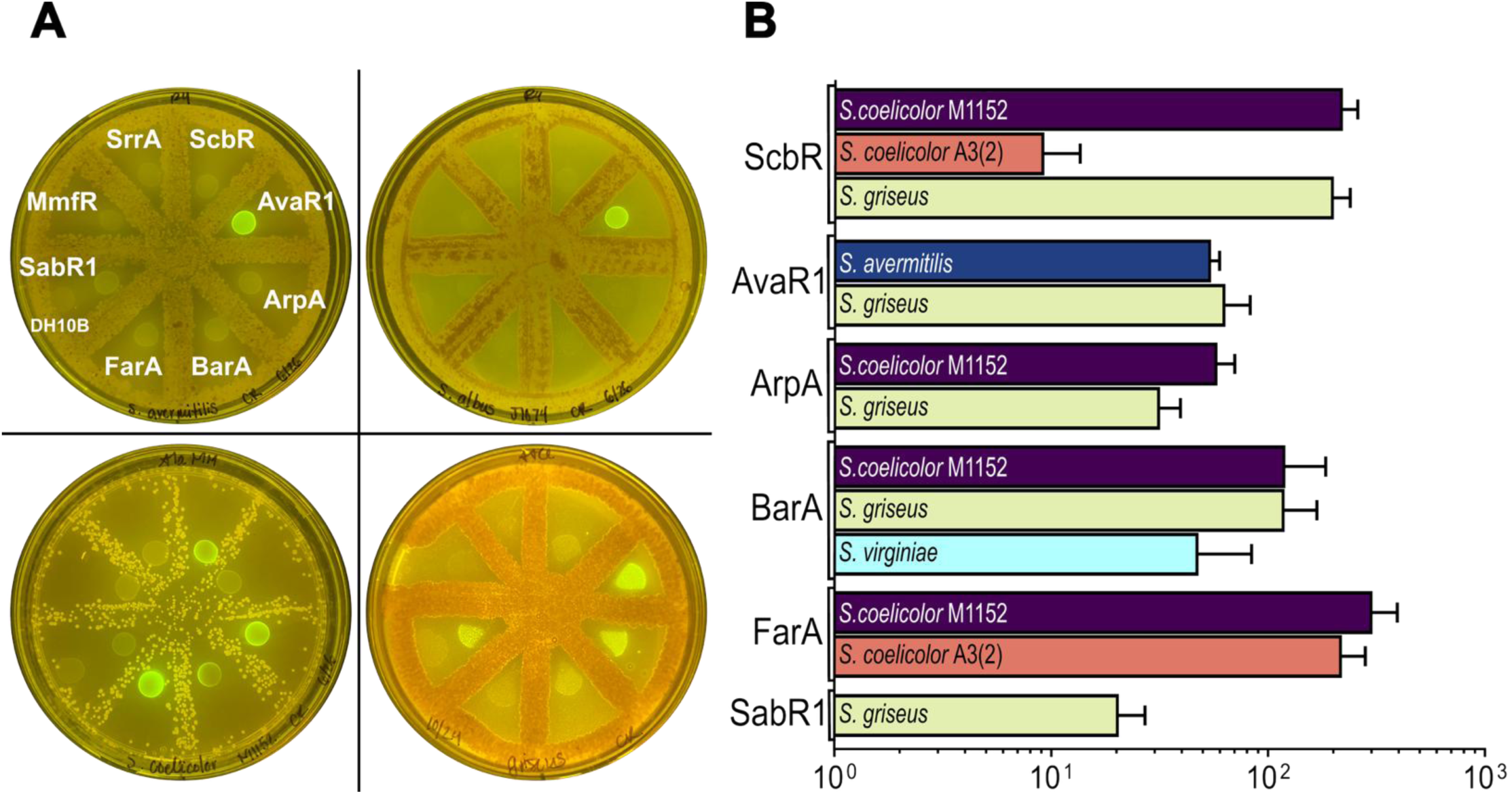
Coculture of biosensor strains with *Streptomyces sp.* and crude extract induction results. **A)** Pictures of *Streptomyces* and *E. coli* biosensor coculture, clockwise from upper left: *Streptomyces avermitilis* B- 31267, *Streptomyces albus* J1074, *Streptomyces griseus* F-5144, *Streptomyces coelicolor* M1152. Image is one representative of biological triplicate. Indicated in white is the map of the CSR biosensor plated in that slice on all coculture plates – DH10B control was plated with SabR1. **B)** Results of crude extract induction analysis. Response is reported as fold change, calculated as average of maximum induced fluorescence over baseline fluorescence. Left label is CSR biosensor, label on the bar is *Streptomyces* sp. crude extract. Results are an average of three biological replicates analyzed on different days. Error bars +/- 1 SD.

### Activation of CSR-based AR Biosensor via Microbial Coculturing and Crude Extract Induction

Historical identification of *Streptomyces* ARs and their cognate CSRs typically involved a single CSR/AR pair identified using a combination of knockout strain generation, heterologous protein expression, gel shift assays, and crude extract or non-stereospecific AR synthesis (**Supplementary Table 3**)^42,43,45,46,55,75–78^. These processes, while very successful, are arduous and lack the scalability necessary to investigate the multitude of CSR quorum sensing circuits^37,76^. After observing robust responses to synthesized ARs, we hypothesized that the CSR vectors could produce qualitative GFP signal when cocultured with *Streptomyces* on solid media. If confirmed, these vectors could be used as biosensors for rapid identification of ARs. Analytical methods of AR identification are challenging, as these small molecules generally have poor absorbance by UV-vis and are difficult to detect via mass spectrometry^79^. For qualitative identification of ARs, plates were streaked with liquid cultures of *Streptomyces* and incubated for two days at 30 °C, after which liquid culture from each CSR vector culture was plated. A vectorless DH10B culture was also tested as a negative control. Gratifyingly, all *E. coli* biosensor cultures grew when plated alongside *Streptomyces*. Additionally, crude extract induction assays were performed by inducing the pTOTAL-CSR cultures with extracts from *Streptomyces* strains grown on solid media. Six out of the eight biosensors generated qualitative fluorescent signal when cocultured on plates and quantitative signal in the crude induction assay. Though visible reduction in GFP signal or *E. coli* colony growth was observed when proximal to *Streptomyces* strains, this did not hinder the activation of biosensors (**Fig 3A**).

The AvaR1 biosensor vector shows robust response to *S. avermitilis,* AvaR1’s strain of origin and producer of the AR avenolide (**Fig. 3A**). This was further confirmed with the crude extract induction assay, where AvaR1 exhibited a 59.8 ± 6.7-fold increase in fluorescence (**Fig. 3B**). The AvaR1 biosensor also demonstrated strong qualitative fluorescence when plated with *S. albus*, a producer of avenolide derivatives (**Fig. 3A**). *S. albus* avenolide derivatives have been previously reported to activate avermectin production in *S. avermitilis*^80^. Our data strongly indicates the derivatives produced by *S. albus* activate avermectin by binding to AvaR1 in a similar manner to *S. avermitilis*’ native avenolide.

*S. virginiae* is the native producer of the ligands for BarA, the VBs. Slight induction was seen with the BarA biosensor when cocultured with *S. virginiae*; no other vectors showed activation (**Supplementary Fig. 7**). Since synthesized VB-D was able to derepress other vectors at high concentrations (*e.g.* ScbR, ArpA, and FarA), absence of fluorescent signal from these biosensors suggests that *S. virginiae* is producing very low levels of VBs under these culturing conditions. (**Fig. 2**, **Supplementary Table 1**). When induced with *S. virginiae* crude extract, the BarA biosensor demonstrated a 33.2-fold increase in fluorescence (**Fig. 3B**). *S. coelicolor* M1152, a mutated strain that exhibits enhanced SCB production, triggered GFP production of its cognate CSR biosensor ScbR, as well as other GBL-type AR CSRs ArpA, BarA, and FarA (**Fig. 3A**, **Fig. 3B**).^81^ The initial enzyme for GBL biosynthesis ScbA, an acyltransferase, is permissive to different β-keto esters, leading to the production of SCBs with varying chain lengths^48^ (**Supplementary Fig. 5**). As indicated in the synthesized molecule results, ArpA and BarA are responsive to SCB1and can likely be activated by other SCB-type GBLs produced by *S. coelicolor* M1152. SrrA and MmfR biosensors did not consistently fluoresce either during plate coculture or crude induction assay, likely due to low level production of the cognate ARs; additional results are discussed in **Supplementary Fig. 7**.

Perhaps the most intriguing results were in response to *S. griseus.* Plate coculture experiments resulted in fluorescence in the CSR vectors ScbR, AvaR1, ArpA, BarA, and SabR1. ArpA is the CSR derived from *S. griseus,* thus an active response for ArpA is expected (**Fig. 3A**). Responses by ScbR and BarA are likely due to structural similarities between ARs explored in the synthesized molecule induction assays. *S. griseus* was previously reported to make a molecule structurally similar to avenolide, suggesting potential reasons for AvaR1 activation (**Supplementary Fig. 8**)^76,82,83^. *S. griseus* has not thus far been identified as a producer of SAB AR derivatives and this work is the first to show crosstalk between SabR1 and ARs produced by this strain. All plate coculture results were confirmed by fluorescence assay results from crude extract of *S. griseus* (**Fig. 3B**).

### Crosstalk Between Regulators and Operators

After exploring AR crosstalk, we examined the ability of each CSR to repress noncognate operators. It is essential to verify the crosstalk between CSRs if they are to be used concurrently in a cell. All CSRs tested are homologs from *Streptomyces* and possess high amino acid sequence similarity in the N-terminus DNA binding motif, indicating a strong potential for the CSR to bind to similar operator motifs (**Supplementary Fig. 4**). The tested operator sequences all possess conserved ‘AT – GC’ motifs with a 10-base spacer (**Fig 4B** and **Supplementary Fig. 9**). To gauge operator crosstalk, a ‘pREP’ vector was cloned that constitutively expressed one of the eight CSRs, and a ‘pGFP’ vector was cloned, which had one of the eight operators within a *gfp* promoter. Each pREP plasmid was co-transformed with each pGFP, generating 64 unique repressor/operator combinations, and all 64 strains were assayed (**Fig. 4A** and **4C**). Using the cognate CSR/operator strain as a benchmark, each CSR was evaluated for its ability to bind to every operator.

**Fig. 4:**
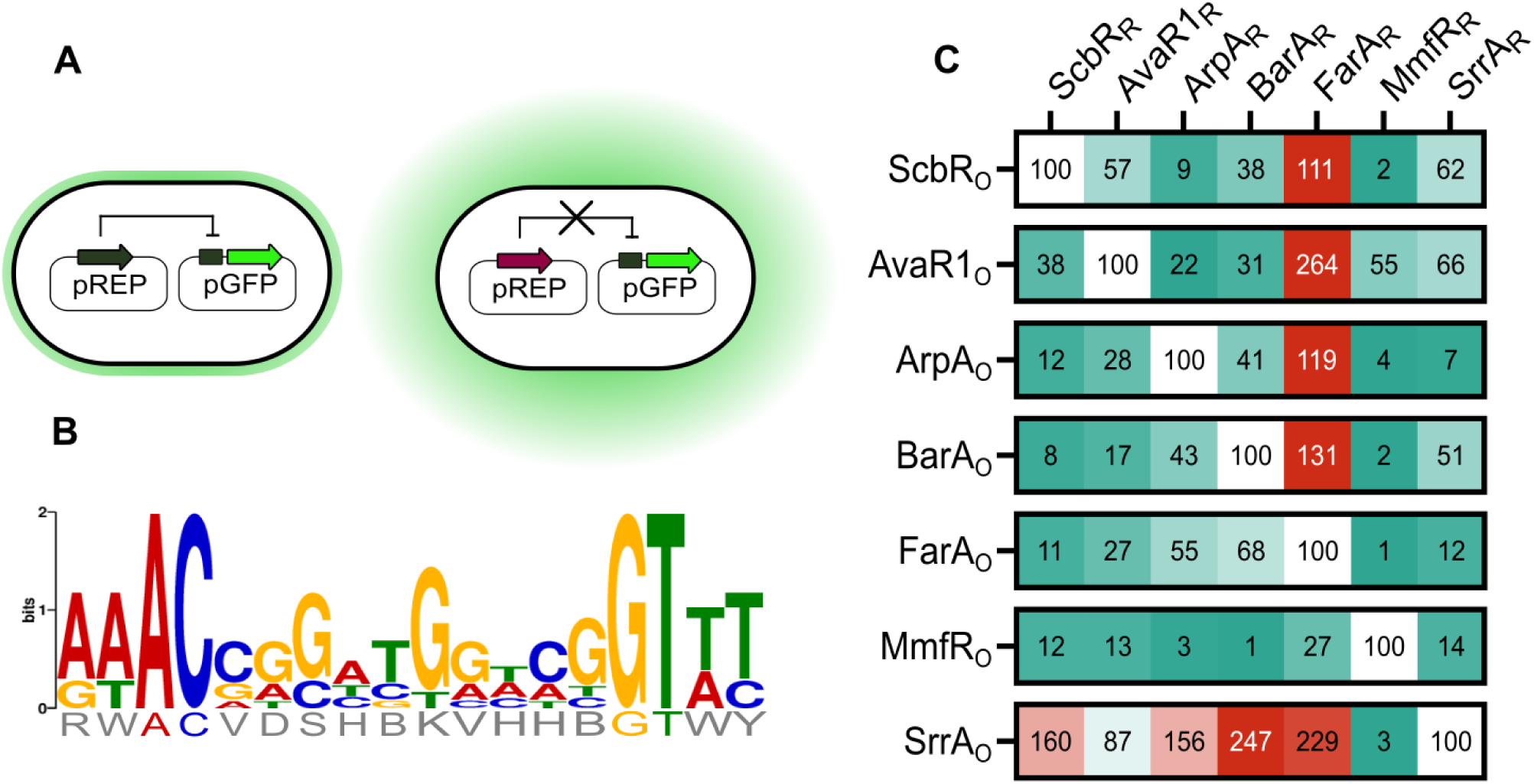
CSR repressor-operator crosstalk. **A)** One vector expressing a CSR and another vector with a CSR_O_- regulated *gfp* were cotransformed into *E. coli* for analysis. CSR that are capable of binding the operator will result in low fluorescence (left), unrepressed operators will allow for high GFP production (right). **B)** Position Weighted Matrix of CSR operator sequences. **C)** Results of operator crosstalk analysis, in percent repression compared to cognate CSR/operator pair. RFU of each cotransformed strain was first divided by the maximum fluorescence of the individually grown pGFP vector with the corresponding operator-*gfp*. The generated fold repression of each strain was further divided by the fold repression of the cognate CSR/operator pair, and results are presented as percent repression as compared to the cognate CSR/operator pair. Values over 100% indicate superior repression than cognate CSR/operator, under 100% show incomplete repression.

As predicted, crosstalk was observed between many of the CSRs. MmfR_R_ exhibited the highest selectivity, with significant activity only being observed for its native operator and some binding to the AvaR1 operator (**Fig 4A**). FarA_R_, on the other hand, demonstrates promiscuous repression of most operators, frequently exhibiting superior repression to the cognate CSR/operator (**Fig 4A**). SrrA_O_ appears to be an operator sequence that is bound by many of the tested repressors (**Fig 4A**). The complexities of SabR1_R_ complicate drawing firm conclusions of its interactions (**Supplementary Fig. 10**). The difference in operator binding specificity for the CSRs is most likely a result of small structural differences made by the residues in the DNA binding region and not the result of a single amino acid substitution (**Supplementary Fig. 4 and 9**). ScbR_R_ and MmfR_R_ both demonstrated low repressor affinity to one another’s tested operator (**Fig 4A**). These results, taken with the orthogonality detailed in the synthesized molecule induction assay, further support that these systems perform as independent regulatory circuits (comporting with published data)^56^. However, ScbR_R_ and MmfR_R_ also demonstrate some ability to bind to the DNA sequence for to AvaR1_O_, indicating a possibility for both repressors to regulate alternative operator sequences if present in the genome.

### CSR Biosensor E. coli Strain Coculture

As previously described, molecules with disparate core structures (GBL vs BN) or chain lengths (SAB1 vs SRB1) exhibit orthogonality in binding to CSRs. Therefore, AR biosensors that fit these parameters may be activated in an orthogonal manner when cocultured. We sought to verify these observations by coculturing two strains, each strain possessing a vector with a different CSR regulating a different fluorophore (**Fig. 5A**). Overnight cultures were subdiluted and mixed 1:1, incubated for outgrowth, then induced with either individual or combined ARs. Results from mCherry vectors demonstrate the well-documented complexity of changing genetic parts (**Supplementary Table 4** and **Supplementary Fig. 11**)^74,84–87^. Baseline fluorescence was different for GFP vectors versus mCherry vectors of the same CSR, and reduction in maximum fold-change was noted, for example FarA-GFP (396.4 ± 43.7-fold) versus FarA-mCherry (153.6 ± 3.2-fold). Maximum fluorescence was consistently lower for mCherry vectors, with GFP RFU maximum ranging from 10^3^ to 10^4^, but no mCherry RFU maximum was observed above 10^3^ for any of the vectors (**Fig. 5B** and **Supplementary Fig. 11**).

**Figure 5:**
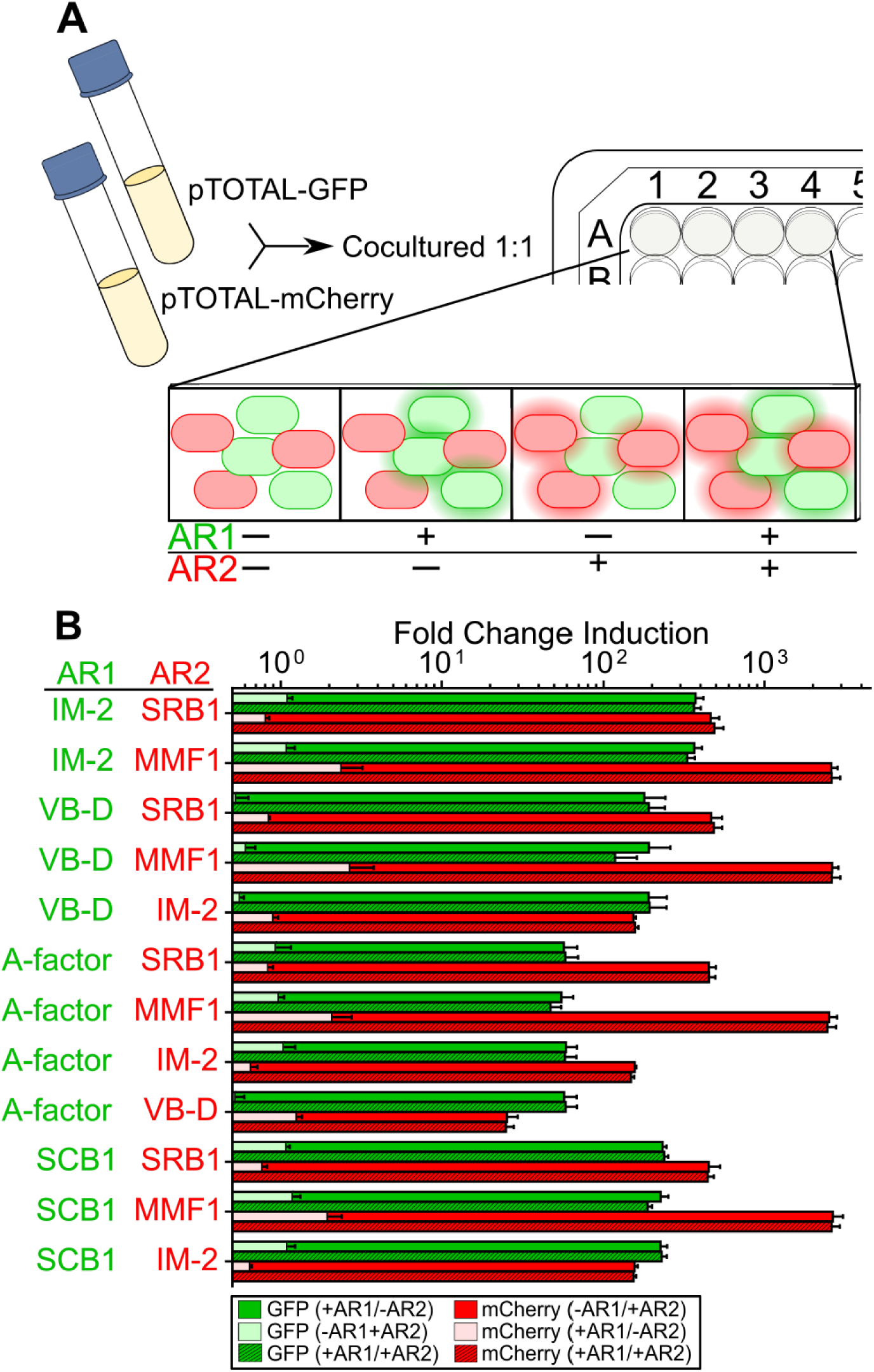
CSR biosensor orthogonality in microbial coculture. **A)** Outline of experimental technique Overnight of *E. coli* cultures with pTOTAL-GPF or pTOTAL-mCherry vectors are cocultured and induced with vehicle, single AR, or both ARs. CSR vectors were induced with the following concentrations: 31.5 μM SCB1 for pTOTAL-ScbR, 0.78 μM A-factor for pTOTAL-ArpA, 6.25 μM VB-D for pTOTAL-BarA. 0.625 μM IM-2 for pTOTAL-FarA, 500 μM MMF1 for pTOTAL-MmfR, 7.8 μM SRB1 for pTOTAL-SrrA. **B)** Results from CSR biosensor coculture experiments. Each vector name is colored by its fluorescent reporter (Green for GFP, red for mCherry). Results are an average fold change over baseline of three biological replicates analyzed on different days. Error bars +/- 1 SD.

Understanding these caveats, the biosensor vectors still demonstrated robust response to cognate ARs in an orthogonal fashion; no crosstalk between ARs was observed (**Fig. 5**). GFP vectors demonstrated fold changes proximal to those observed in the previous synthesized molecule inductions (**Supplementary Table 1** and **Fig. 5B**). The mCherry vector also reported excellent fold-changes, in particular, pTOTAL-SrrA-mCherry and pTOTAL-MmfR-mCherry exhibited a 463.0 ± 15.4-fold and 2.59×10^3^ ± 62.2-fold change, respectively. Fluorophore expression stayed consistent when cocultured with different vectors, for example, whether cocultured with FarA-mCherry, MmfR-mCherry, or SrrA-mCherry, ScbR-GFP exhibited a fold change average of 223.8 ± 16. While fine-tuning will be required when implementing these parts in higher-order genetic circuits, these experiments confirm *Streptomyces* CSR/AR biosensors as capable of differential interpretation of signal and response in a coculture setting.

### AND Gate Construction Using CSRs MmfR and SrrA

With the operator and AR orthogonality established, we endeavored to combine individual CSR vectors into a dual input AND gate. Multi-input genetic circuits are necessary for engineered cellular sense and response systems^17,22^. To determine which CSRs were to be used for the AND gate, we considered protein expression burden, baseline and maximum fluorescence, as well as AR and operator orthogonality. We decided to construct a CSR vector such that MmfR and SrrA form an AND gate that requires the addition of both ARs (SRB1 and MMF1) for full induction of GFP (**Fig. 6A-C**). When induced with 7.8 μM SRB1 or 500 μM MMF1, inductions of 5.5 ± 1.2-fold and 25.1 ± 4.6-fold were observed, respectively (**Fig. 6D**). Fluorescence induction of 165.6 ± 31.3- fold occurred when the strain was induced with both molecules simultaneously (**Fig. 6D**). Improvements on the design of the vector may reduce unwanted induction in the single molecule conditions, including placement of operator or modifying the repressor’s promoter regions^19^.

**Fig 6:**
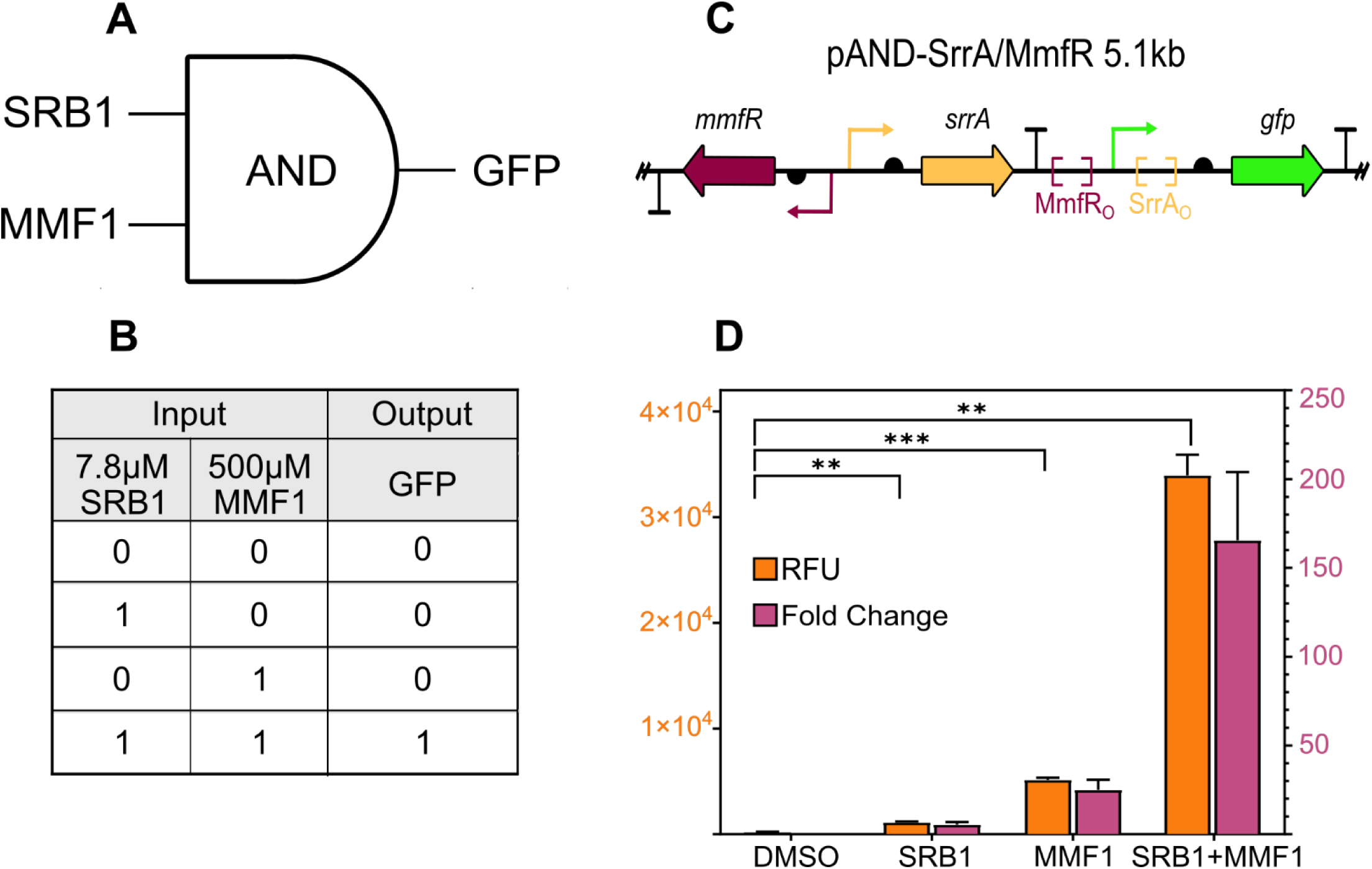
AND Gate genetic circuit using SrrA and MmfR CSRs. **A)** Logic gate and **B)** Truth table of input combinations of CSR two input AND gate. **C)** Diagram of pAND plasmid. MmfR and SrrA are expressed by independent promoters; operators were cloned flanking the -35/-10, with MmfR_O_ upstream and SrrA_O_ downstream. **D)** Results from individual and dual molecule induction. RFU presented on left Y axis, fold change on right Y axis. pAND induction with individual molecules results in some observed change in fluorescence, maximum induction occurs when both ARs are added. Significance was determined using a paired, two-tailed Student’s *t*-test, p <0.05.

## Discussion

Inducible, repressor-based genetic circuits have been a versatile cornerstone of modern synthetic biology^18,23,24,27,88–92^. Overall, our work demonstrates the utility of TetR-like CSRs and their ARs as diversifiable components of the synthetic biology toolkit. Building on previous work, we devised improved vectors to test a suite of CSRs against cognate and non-cognate molecules and detailed these CSR/AR interactions. This analysis was enabled by efficient and diversifiable synthetic routes for accessing ARs, which were developed in house. Additionally, for the first time, we have been able to explore the structure activity relationships for all classes of *Streptomyces* ARs with purified ligands, revealing their high selectivity for their native ligands. The lowest selectivity was observed for the GBL ARs such as SCB1, A-factor, and VB-D, which is logical given their high structural similarity. However, even for the GBLs, the selectivity of the CSRs is more than sufficient for use as AR biosensors and in genetic circuits. *In silico* circuit design, coupled with combinatorial design or directed evolution, could be used to improve the CSR circuits, crafting biosensors with less toxicity, more sensitivity, and maximally efficient ON/OFF states for advanced genetic architecture^20,88,93–96^.

The CSR-based AR biosensors demonstrated robust sensitivity when presented with ARs as synthesized molecules, in crude extract, or during microbial coculture. Our solid media *Streptomyces*/biosensor coculturing technique, coupled with the crude extract induction, provides tests to assess the presence of ARs produced either in liquid culture or on solid media. We assert these experimental techniques can be widely adapted to rapidly assess organisms for their ability to produce *Streptomyces* quorum sensing ARs. Future analysis will yield greater understanding of small-molecule signaling in Gram-positive organisms, which have been described as possessing primarily peptide-based quorum sensing^1,4,12,38,39,97^. These investigations may reveal novel inter and intra species quorum sensing communication and harnessing this interoperable signaling may yield synthetic biology applications.

We assayed each CSR for its ability to bind to the operators of the other CSRs and confirmed operator crosstalk is widespread in the CSRs. This crosstalk is likely observed because of the sequence similarity of the DNA binding domains of the CSRs. We identified FarA as a promiscuous binder of operators, while the SrrA operator sequence was repressed by many of the CSRs with greater or equal strength as the cognate CSR SrrA. We also confirmed that, for the operator and molecule combinations tested, the quorum sensing systems ScbR/SCB and MmfR/MMF show orthogonality. Understanding the full range of CSR binding sequences may lead to more orthogonal operator/CSR combinations and increase the number of CSR circuits that can be combined in a single cell. Having a complete picture of CSR binding will also identify the CSR quorum sensing regulon *in vivo*. The vast number of CSRs can aid research into ligand-receptor interactions, providing insight on DNA-binding or ligand-binding domain engineering, or as building blocks for biocontainment structures^19,21,88,98–100^.

A necessary component for higher order genetic circuits is independent processing of input signals ^20,101–105^. After determining the ligand specificity of each CSR, we tested numerous combinations of molecules to verify their orthogonality in coculture. No crosstalk was observed in any of the tested combinations, and the activation of the GFP-expressing biosensors were indistinguishable from their performance during single molecule induction. NP ARs have demonstrated a robust ligand orthogonality compared to AHL biosensors and provide an alternative to monosaccharide or other primary metabolite biosensors, which have been implicated in circuit failure^19,21,22^.

*Streptomyces* AR biosensors are an example of implementing synthetic biology techniques to address questions of both basic and applied science^106^. We demonstrated that these highly modular inducible genetic circuits can rapidly identify the presence of ARs responsible for regulating NPs. We also assert these biosensors can have future applications for biomanufacturing. *Streptomyces* sp. chassis struggle with genetic tractability, fastidious growth conditions, and a reduced bioengineering toolkit^107–111^. Nevertheless, *Streptomyces* strains are currently used as biomanufacturing organisms for production of their essential therapeutic chemicals^112,113^. Research has linked *Streptomyces* AR quorum sensing to primary metabolism regulation, suggesting these molecules are involved in harmonization of primary and secondary metabolism when producing NPs ^114–116^. Understanding this balancing act may provide insight on designing maximally efficient industrial strains^2,117–119^. Researchers interested in producing value-added compounds via microbial biomanufacturing are increasingly looking toward microbial consortia, or coculturing, as a way to divide the metabolic burden of producing a desired product ^120–125^. *Streptomyces’* ability to genetically maintain and express a wide variety of complex molecules makes them a strong candidate for implementation in advanced microbial consortia biomanufacturing ^122,125–129^. Their capacity is made even more intriguing with ongoing research into rationally designing NP biosynthesis enzymes to make NP derivatives or new-to-nature chemicals^130–132^. Synthetic biology-driven insights into *Streptomyces* Natural Product regulation promises impactful biotechnology advancements.

## Methods

A description of the methods used in this paper can be found in the Supplementary Information document.

## Abbreviations

NP: Natural Products
CSR: Cluster-situated Regulators
AR: Autoregulators
AHL: Acyl-Homoserine Lactone
BGC: Biosynthetic Gene Cluster
GBL: gamma-butyrolactone
BN: butenolide
SCB: *Streptomyces* coelicolor butanolides
SAB: *Streptomyces* autoregulator butenolides
A-factor: autoregulatory factor
IM: inducing molecule
VB: virginiae butanolide
SRB: *Streptomyces rochei* butenolide
MMF: methylenomycin furan
UTR: untranslated region
GFP: Green Fluorescent Protein
TBAF: Tetrabutylammonium Fluoride
RFU: Relative Fluorescence Units

## Author Contributions

L.E.W. and E.I.P. conceived of the project. L.E.W designed and built the DNA constructs and designed the experimentation. H.H. and K.C. synthesized the molecules for the synthetic autoregulator induction assays. L.E.W performed the synthetic molecule induction assay, *E. coli* biosensor coculture, and AND gate protocols. L.E.W. and C.R. designed the *Streptomyces*/*E. coli* coculture and crude induction experiments, L.E.W., C.R. and M.F.G. performed the experimentation. M.F.G. performed the operator crosstalk experimental replicates, as well as replicates for the synthetic molecule induction assay. Z.B. and C.Z. performed the Alpha Fold generation and autoregulator docking. L.E.W. performed data analysis, designed figures, and wrote the manuscript. All authors participated in discussion and revision of the manuscript.

## Acknowledgments

We thank A. Eggly for supplying the MMFs used in this study. This work was funded by an NSF CAREER Award to E.I.P. (CHE 223689). L.E.W. was supported by a Purdue Research Foundation Ross-Lynn Grant. Zachary Logan Budimir acknowledges the National Science Foundation for support under the Graduate Research Fellowship Program (GRFP) under grant number DGE-1842166. This work was supported in part by the Research Instrumentation Center in the Department of Chemistry at Purdue University. The authors acknowledge the support from the Purdue Center for Cancer Research, NIH grant P30 CA023168.

