## Supplementary Information and Methods for "*Streptomyces* Autoregulator Biosensors from Natural Product Cluster-Situated Regulators"

**References** .......... **Error! Bookmark not defined.**

### Supplementary Tables

| Name | $K_D$ (μM) | Hill Coefficient | Ymax | Ymin | Average Experimental Maximum | Average Baseline RFU | Average Maximum Fold change | Minium Effective Concentration (nM) |
|---|---|---|---|---|---|---|---|---|
| **ScbR/SCB1** | **13.3** | **2.73** | **54525** | **302.3** | **55886.8± 1696.9** | **238.2 ± 18.2** | **235.5 ± 12.0** | **1950** |
| ScbR/A-factor | 117.3 | 2.00 | 59629 | 324.5 | 53350.3 ± 1757.1 | 238.2 ± 18.2 | 225.5 ± 21.7 | |
| ScbR/VB-D | 146.3 | 2.13 | 61156 | 272.7 | 53889.7 ± 1572.1 | 238.2 ± 18.2 | 227.7 ± 20.3 | |
| **ArpA/A-factor** | **0.39** | **3.13** | **20529** | **356.4** | **21753.9 ± 2967.7** | **412.5 ± 35.7** | **53.7 ± 11.2** | **159** |
| ArpA/SCB1 | 18.15 | 1.72 | 20782 | 251.3 | 20985.3 ± 2442.7 | 412.5 ± 35.7 | 50.9 ± 3.9 | |
| ArpA/VB-D | 121 | 1.94 | 19301 | 362.1 | 19835.8 ± 2094.3 | 412.5 ± 35.7 | 48.3 ± 5.6 | |
| **BarA/VB-D** | **1.34** | **2.87** | **4957** | **22.16** | **4919.0 ± 595.7** | **20.7 ± 3.4** | **240.3 ± 21.6** | **195** |
| BarA/SCB1 | 15.52 | 2.82 | 1982 | 20.81 | 2031.0 ± 359.3 | 20.7 ± 3.4 | 98.4 ± 7.3 | |
| BarA/A-factor | 41.72 | 2.72 | 5017 | 14.81 | 4841.8 ± 790.3 | 20.7 ± 3.4 | 237.5 ± 41.2 | |
| **FarA/IM-2** | **0.03** | **2.49** | **80110** | **307.9** | **79844.6 ± 2552.1** | **203.3 ± 18.2** | **396.4 ± 43.7** | **4** |
| FarA/VB-D | *357* | *1.60* | *115326* | *159.8* | 64573.9 ± 2246.7 | 203.3 ± 18.2 | 319.8 ± 26.9 | |
| **SabR1/SAB1** | **0.72** | **1.11** | **10424** | **317.1** | **10394.2 ± 787.1** | **359.0 ± 25.1** | **29.2 ± 4.0** | **98** |
| **SrrA/SRB1** | **6.83** | **3.07** | **2248** | **1.258** | **2379.6 ± 262.4** | **2.4 ± 0.1** | **1042.4 ± 338.5** | **970** |
| **MmfR/MMF1** | **110.8** | **3.14** | **54708** | **21.38** | **55473.2 ± 8096.8** | **11.3 ± 1.9** | **4909.7 ± 112.1** | **1950** |

'Average Experimental Maximum' is the average of the highest quantified fluorescence of each biological replicate
'Average Baseline RFU' is an average of the fluorescence of the DMSO control cultures
'Average Maximum Fold Change' is the largest fold change of each biological replicate averaged

**Supplementary Table 1: CSR biosensor response function parameters and experimental results.** Results from the synthetic molecule induction assay. The name indicates the CSR and the molecule tested (CSR/AR). Cognate pairs of CSR/autoregulator are bolded. Grey values, $K_D$ (in μM), Hill coefficient, $Y_{max}$ and $Y_{min}$, were determined by fitting experimental values to a Hill function. Note for 'FarA/VB-D' the tested induction concentrations did not manifest a sigmoidal curve.

| CSR Vector | Average OD600 | Growth Reduction |
|---|---|---|
| pTOTAL-ScbR | 0.134 | 17% |
| pTOTAL-AvaR1 | 0.127 | 21% |
| pTOTAL-ArpA | 0.114 | 29% |
| pTOTAL-FarA | 0.143 | 11% |
| pTOTAL-BarA | 0.140 | 13% |
| pTOTAL-SabR1 | 0.116 | 28% |
| pTOTAL-MmfR | 0.096 | 41% |
| pTOTAL-SrrA | 0.156 | 3% |
| DH10B Control | 0.162 | |

**Supplementary Table 2: Growth comparison of CSR biosensor strains.**
$OD_{600}$ from a subset of pTOTAL-CSR vector cultures from one biological replicate were averaged. During the synthetic molecule induction assay trials, all vectors diluted from overnight cultures took ~3-4 hours to reach $OD_{600}$ 0.1-0.2. Notably, pTOTAL-AvaR1, pTOTAL-ArpA, pTOTAL-SabR1, and pTOTAL-MmfR all demonstrate a reduction in $OD_{600}$ >20% when compared to a vectorless DH10B control induced with vehicle (DMSO)

### **Supplementary Table 3:** Overview of Historical CSR/Cognate Autoregulator Investigations.

| CSR | Year/ Author | Strain/**AR** | Media | Protocol Summary | MEC/$K_D$ | Comments |
|---|---|---|---|---|---|---|
| ScbR | 2000/ Takano[1] | *S. coelicolor* M145/ **SCB1** | SMMS Agar | Various concentrations of SCB1 were added to 8 mL molten agar, poured into 9-cm dishes. Spores of the strain were spread on the solid media and visible inspection of the plates determined the presence of precocious pigment production after ~33 hours | 256 nM | Possesses SCB1 biosynthetic gene cluster and SCB1 biosynthesis capabilities |
| | 2001/ Takano[2] | **SCB1** | | Electrophoretic Mobility Shift Assay (EMSA) | 1 µg (1.6 µM) | MEC is described as the 'concentration at which the formation of the DNA–protein complex was markedly reduced'. Not a result of a dilution gradient. |
| | 2009/ Hsaio[3] | *S. coelicolor* LW16 (*ΔscbA/ΔscbR)*::pTE134/ **SCB1** | Difco Nutrient Agar | Strain spores were spread on a plate supplemented with kanamycin. Concentrations of SCB1 were aliquoted onto the spores, then incubated for 3 days. The presence of resistant colonies indicated the derepression of ScbR by a cognate ligand. | 0.025 µg | pTE134 integration re-introduces *scbR* and a kanamycin resistance gene with a modified promoter, resulting in ScbR-regulated production of kanamycin resistance. |
| ArpA | 1982*/ Hara, Beppu[4] | *S. griseus* FT-1, A-factor production deficient mutant/ **A-factor** | Nutrient Agar | The strain was grown for 2 days, homogenized then an aliquot was mixed with nutrient soft agar and overlaid on the nutrient agar base plate. A paper disc containing A-factor was placed onto the agar plate. After a 2-day outgrowth, a nutrient soft agar containing *B. subtilis* was overlaid. The plate was further incubated for ~1 day and the diameter of the zone of inhibition measured. | 2 ng in text | Strain was generated by UV irradiation<br>Fig. 3 shows MEC of 0.01 µg |
| | 1982*/ Hara, Beppu[5] | *S. griseus* FT1, IFO 13189, 2247, IFO13350/ **A-factor** | Liquid GMP medium | Seed cultures of the strains were subdiluted into fresh media containing various concentrations of streptomycin, with or without A-factor. Growth was observed after an additional incubation of 3 days. | 1 ug/mL | All strains A-factor deficient, growth indicates rescue of streptomycin production induced by A-factor<br>*These two works by Hara and Beppu are cited as the source of the observation that A-factor stimulates streptomycin production at 1 nM |
| | 1989/ Miyake[6] | *S. griseus* IFO 13350, A-factor production deficient mutant/ **A-factor** | | The strain was grown to log phase in liquid media and crude cell extract was generated by lysing the pelleted mycelia. Cell extract (containing ArpA protein) was combined with radiolabeled A-factor. Saturation of binding was quantified using Scatchard analysis comparing free A-factor to ArpA-bound A-factor. | ~0.7 nM | Strain was generated by heat treatment<br>[Named *S. bikiniensis* in source publication[4]] |
| | 1997/ Onaka, Horinouchi[7] | **A-factor** | | EMSA | 32-160 nM | Noted aggregation of ArpA during protein isolation. This observation has been widely reported when isolating *Streptomyces* CSRs and observed by our group. |
| BarA | 1988/ Nihara[8] | *S. virginiae*/ **VB-C** | Liquid F Medium. | Liquid cultures of the strain were subject to rounds of subdilution and outgrowth, then an aliquot was combined with VB diluted in fresh media and grown for an additional 4 hours. The culture was centrifuged, and supernatant was analyzed for antibiotic activity using disk diffusion on a confluent lawn of *B. subtilis.* | 0.8 ng/mL | Strain history is unclear. Possibly possesses WT VB biosynthetic gene cluster and VB biosynthesis capabilities (implied in Thao et al. 2017) *B. subtilis* media: 5 g polypeptone, 3 g meat extract, and 15g/L agar |
| | 2017/ Thao[9] | *S. virginiae* (Δ*barX*) / **VB-C** | Liquid F medium | References Nihira et al. 1988 protocol[8] | 79 nM | Strain is a VB synthase knockout |
| FarA | 1989/ Sato[10] | *S. lavendulae* FRI-5/ **IM-2** | Liquid F medium | Liquid cultures of the strain were subject to rounds of subdilution and outgrowth, then an aliquot of a 5-hour cell suspension was added to of fresh medium with dilutions of IM-2. The production of a blue pigment was monitored over time | 0.6 ng/mL | Strain history is unclear. Possibly possesses WT IM-2 biosynthetic gene cluster and IM-2 biosynthesis capabilities (implied in Thao et al. 2017) |
| | 2017/ Thao[9] | *S. lavendulae* FRI-5 (ΔfarX)/ **IM-2** | Liquid F medium | References Sato et al.1989 protocol[10] | 5 nM | Strain is an IM-2 synthase knockout |
| SabR1 | 2018/ Wang[11] | *S. ansochromogenes* (*ΔsabA*)/ **SAB1** | Liquid SP medium | SAB1 was added into cultures of the strain at the beginning of fermentation. After incubation for 5 days, nikkomycin production was analyzed by HPLC | 5 nM | Strain is a SAB synthase knockout. Production was fully restored to WT strain levels at 100 nM |
| SrrA | 2012/ Arakawa[12] | *S. rochei* 7434AN4 KA20 (ΔsrrX)/ **SRB1** | Liquid YM medium | A seed culture of the strain was supplemented with an aliquot of purified SRB and grown for 24 h. Lankacidin and lankamycin production was analyzed by HPLC | 40 nM | Strain is a SRB synthase knockout |
| MmfR | 2008/ Corre[13] | *S. coelicolor* W81*(ΔmmfLHP)*/ **MMF1** | AlaMM | Two cubes were cut out from an agar lawn of Methylenomycin-sensitive *S. coelicolor* M145. Two agar plugs were added, one contained purified MMF1, the other was from a confluent lawn of *S. coelicolor* W81 cells. A zone of inhibition of *S. coelicolor* M145 around the W81 plug indicated MMF1-induced production of Methylenomycin. Production also confirmed by HPLC | 1 ug | *S. coelicolor* A3(2) possesses the linear plasmid SCP1, containing the biosynthetic gene clusters for Methylenomycin and the MMF autoregulators which regulate the production of Methylenomycin. W81 is a derivative of A3(2) without MMF biosynthetic genes, retains Methylenomycin genes. |
| MmfR | 2021/ Zhou[14] | **MMF1** | | EMSA | 0.6± 0.04 nM | |
| | 2011/ Kitani[15] | *S. avermitilis* KA320 (*Δaco*)/ **Avenolide** | Liquid Production Media | Liquid cultures of strain were supplemented with various concentrations of pure avenolide, supernatant analyzed by HPLC for production of avermectin | 4 nM | Strain is an avenolide synthase knockout<br>Media is Medium B from Burg et al. 1979[16]<br>Saturation of NP induction at 50–150 nM of avenolide |
| AvaR1 | 2017/ Thao[9] | *S. avermitilis* KA320 (*Δaco*)/ **Avenolide** | YMS-MC Agar | Strain spores were spread onto media pre-mixed with avenolide and incubated for 8 days. The agar culture was analyzed by HPLC to evaluate avermectin production | 8 nM | Strain is an avenolide synthase knockout |
| | 2020/ Nair[17] | **Avenolide** | | Isothermal Titration Calorimetry | 42.5 ± 2.1 nM | |

| Individual Cultures | OD600 at induction | Cocultures (GFP/mCherry) | OD600 at induction |
|---|---|---|---|
| pTOTAL-ScbR-GFP | 0.273 | ScbR/FarA | 0.251 |
| pTOTAL-ArpA-GFP | 0.176 | ScbR/MmfR | 0.196 |
| pTOTAL-BarA-GFP | 0.178 | ScbR/SrrA | 0.238 |
| pTOTAL-FarA-GFP | 0.235 | | |
| | | ArpA/BarA | 0.215 |
| pTOTAL-FarA-mCherry | 0.235 | ArpA/FarA | 0.215 |
| pTOTAL-BarA-mCherry | 0.210 | ArpA MmfR | 0.151 |
| pTOTAL-MmfR-mCherry | 0.118 | ArpA/SrrA | 0.184 |
| pTOTAL-SrrA-mCherry | 0.203 | | |
| DH10B Control | 0.242 | BarA/FarA | 0.221 |
| | | BarA/MmfR | 0.167 |
| | | BarA/SrrA | 0.194 |
| | | FarA/MmfR | 0.166 |
| | | FarA/SrrA | 0.225 |

**Supplementary Table 4: $OD_{600}$ at time of induction for CSR biosensor coculture induction assay.** Each individual culture was grown overnight, diluted 1:100, then one pTOTAL vector with GFP was combined 1:1 with a pTOTAL vector with mCherry. Combined cultures were aliquoted into a microplate for outgrowth to $OD_{600}$ ~0.15-0.25. For the control, individual pTOTAL-GFP and pTOTAL-mCherry cultures were plated, their $OD_{600}$ for a representative biological replicate at time of experimental induction is on the left. On the right is the $OD_{600}$ for the combined cultures at time of induction.

### Supplementary Figures

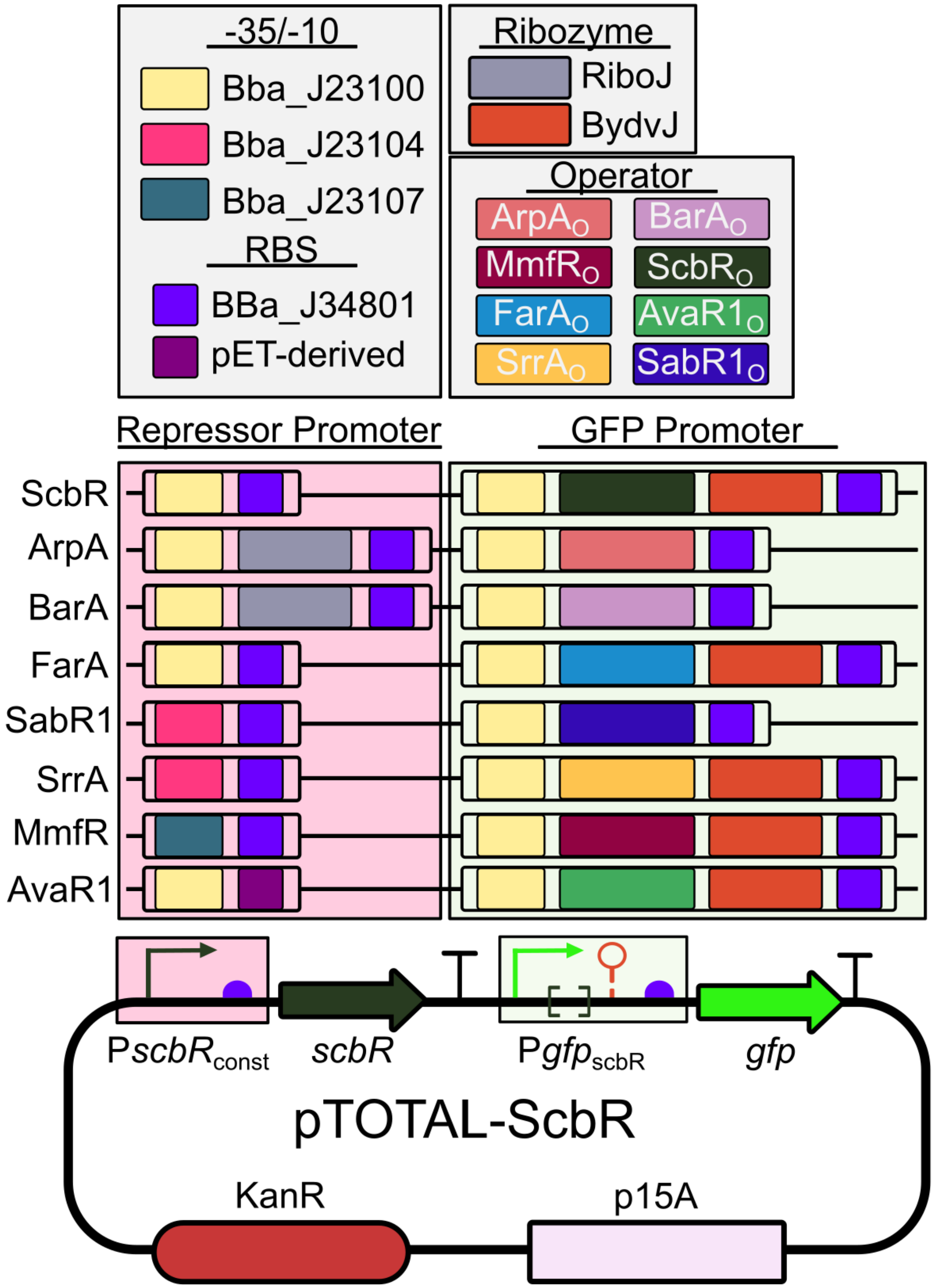

**Supplementary Figure 1: Detailed parts description of pTOTAL vectors.**
The promoter region that expresses each CSR was uniquely tailored to maximize protein production and minimize toxicity, while also reducing the baseline fluorescence of each CSR biosensor vector. Parts are identified by either their Registry of Standard Biological Parts identifier or relevant name. The *gfp* promoter for each vector consists of a strong -35/-10, the cognate operator to the CSR, a Ribosomal Binding Site (RBS), and for some vectors the genetic insulator BydvJ. Promoter notation identifies the gene the promoter regulates, subscript is constitutive (const) or the name of the regulator that acts on the promoter.

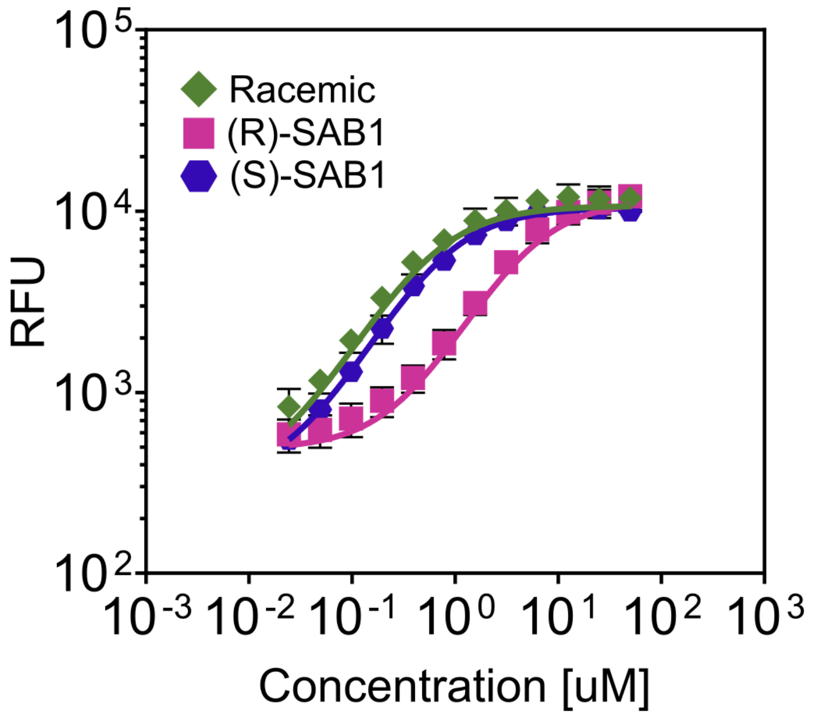

| Molecule | $K_D$ (μM) | Hill Coefficient | Ymax | Ymin | Average Experimental Maximum | Average Baseline RFU | Average Maximum Fold change |
|---|---|---|---|---|---|---|---|
| Racemic | 0.7174 | 1.108 | 10424 | 317.1 | 10394.2 ± 787.1 | 359.0 ± 25.1 | 29.2 ± 4.0 |
| (*R* )-SAB1 | 4.897 | 1.146 | 11464 | 488.4 | 10596.4 ± 1085.4 | 359.0 ± 25.1 | 29.8 ± 5.1 |
| (*S* )-SAB1 | 0.5522 | 1.074 | 10696 | 305 | 10886.5 ± 2084.4 | 359.0 ± 25.1 | 30.9 ± 8.4 |

**<u>Supplementary Figure 2:</u> SAB1 Epimers with SabR1 Biosensor Vector.**
The SabR1 CSR biosensor vector was induced with (*R*)- and (*S*)-SAB1, as well as the racemic (reproduced from **Supplementary Table 1**). The table below shows the response function parameters for each derivative tested in the assay.

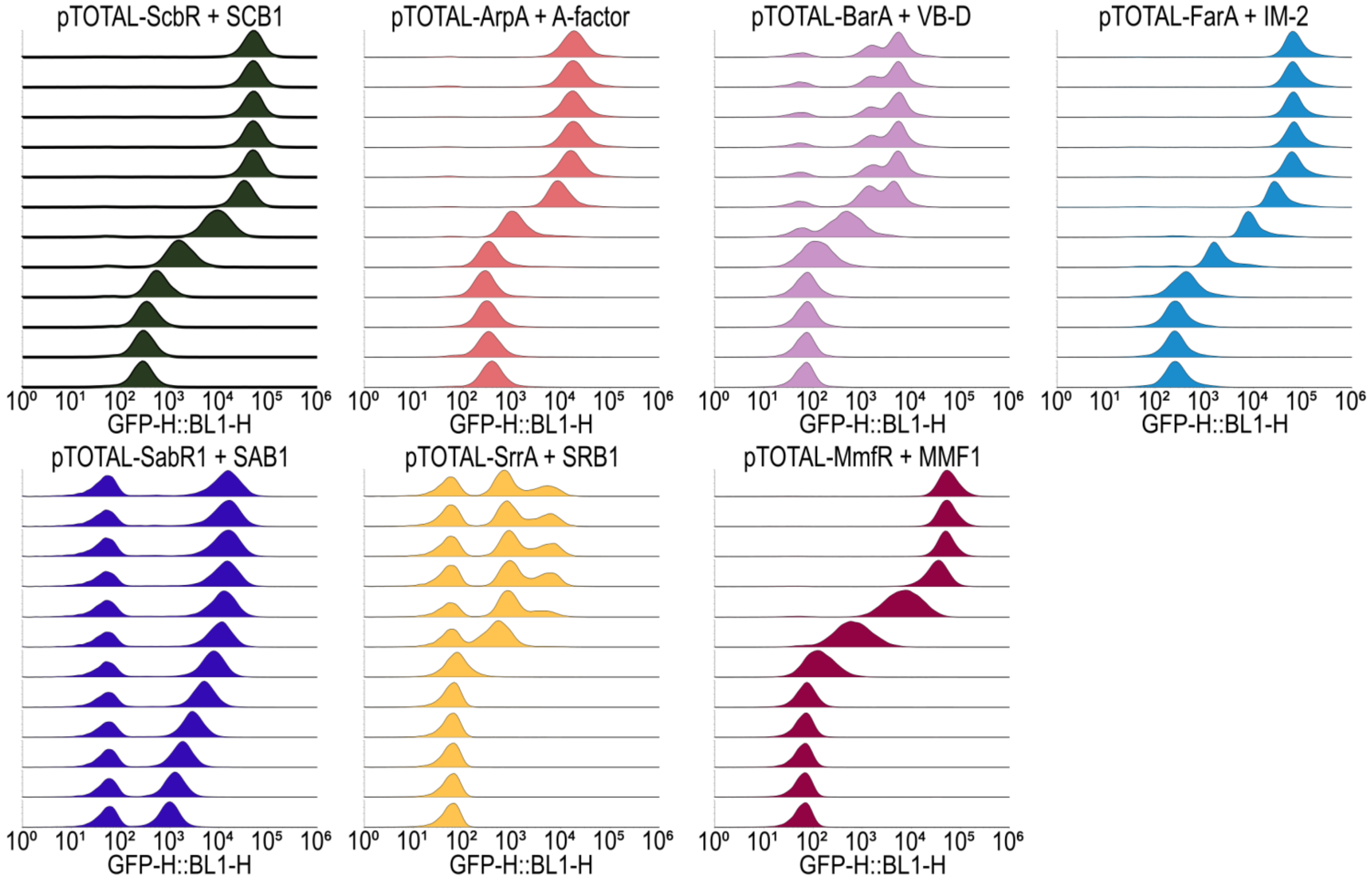

**<u>Supplementary Figure 3:</u> Flow histogram plots for the synthesized molecule induction assay.**
Histograms of GFP RFU from one biological replicate of flow cytometry results are shown. Histograms are ordered with highest concentration of inducing molecule at the top, descending in concentration. pTOTAL-BarA, pTOTAL-SabR1, and pTOTAL-SrrA flow histograms show subpopulations of GFP expression within the cultures. One explanation for the variation would be the CSR expression affecting the retention of, or GFP expression from, the plasmid[18]. However, there is not a perfect link between the burden imposed by the plasmid (be it plasmid replication, CSR expression, or GFP expression) and exhibiting subpopulations of GFP expression. Although pTOTAL-MmfR shows a significant reduction in growth rate (41%), events of the induction curve exhibit a normal distribution; pTOTAL-ArpA also shows a reduction in growth but normal histograms. pTOTAL-BarA has a marginal reduction of 13%, and pTOTAL-SrrA a small 3% reduction, but both have multimodal GFP induction responses. These cultures also have a subpopulation of cells exhibiting the baseline fluorescence of a vectorless culture, indicating loss of plasmid or mutation of GFP expression.

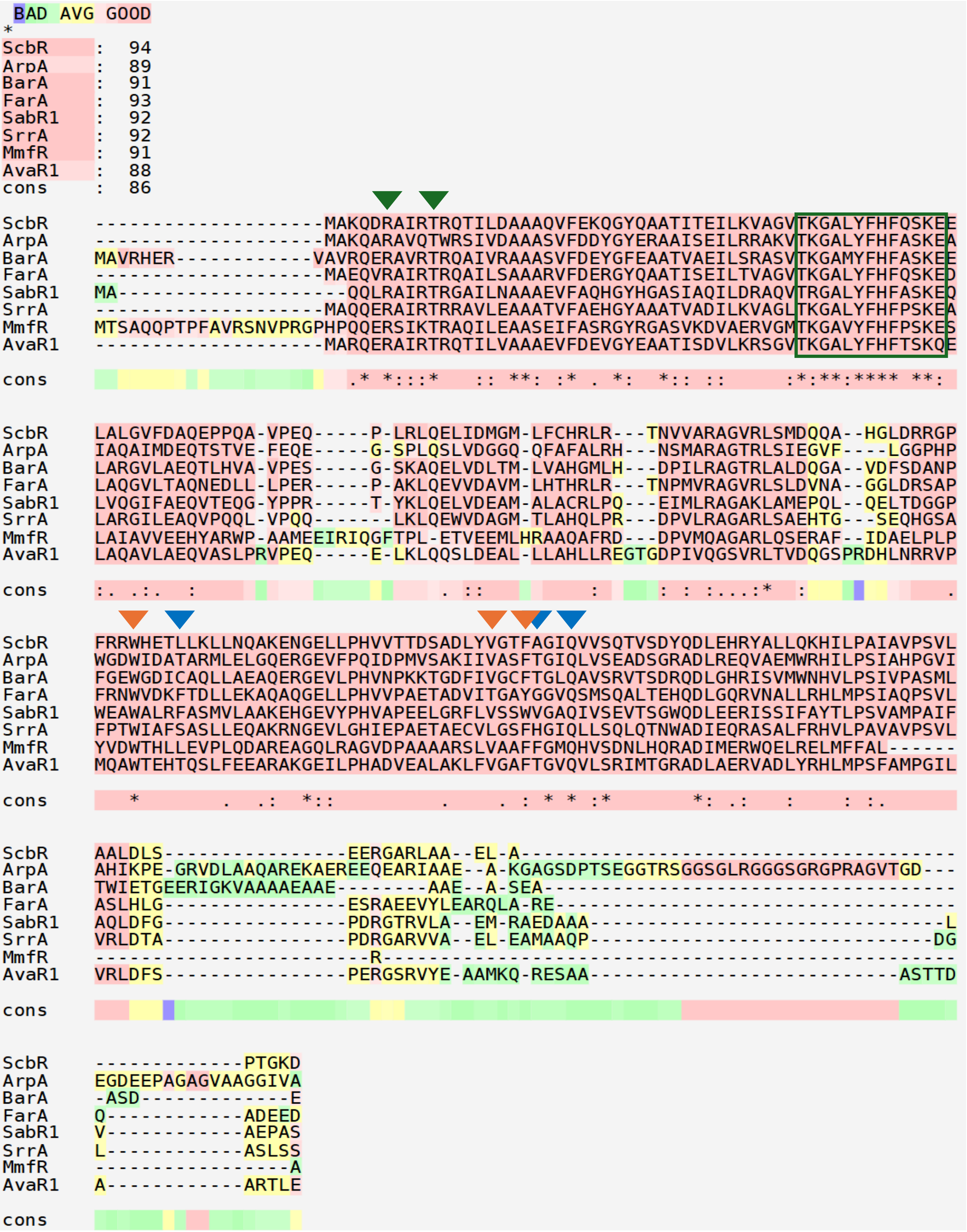

**<u>Supplementary Figure 4:</u> Multiple Sequence Alignment of CSRs.**
The CSRs demonstrate a high level of sequence similarity, especially in the N-terminus DNA binding domain. Conserved DNA binding motifs and residues are indicated by the green arrow and square. Ligand binding residues are indicated by the blue arrow for lactone ring and orange arrow for chain[14,17]. The ArpA coding sequence used for this study is as previously reported[7,19]. One unique characteristic of ArpA, relative to the other CSRs, is its extended C terminus sequence. ArpA in most literature is this ~29 kDa protein, however, it was first identified as 26 kDa protein, much more similar in size to the other CSRs in this studies' cohort[20]. This extended region is possibly the result of sequencing error; if this region is inaccurate, it could affect the ability of the ArpA protein to dimerize, interact with DNA, or bind to its autoregulator.

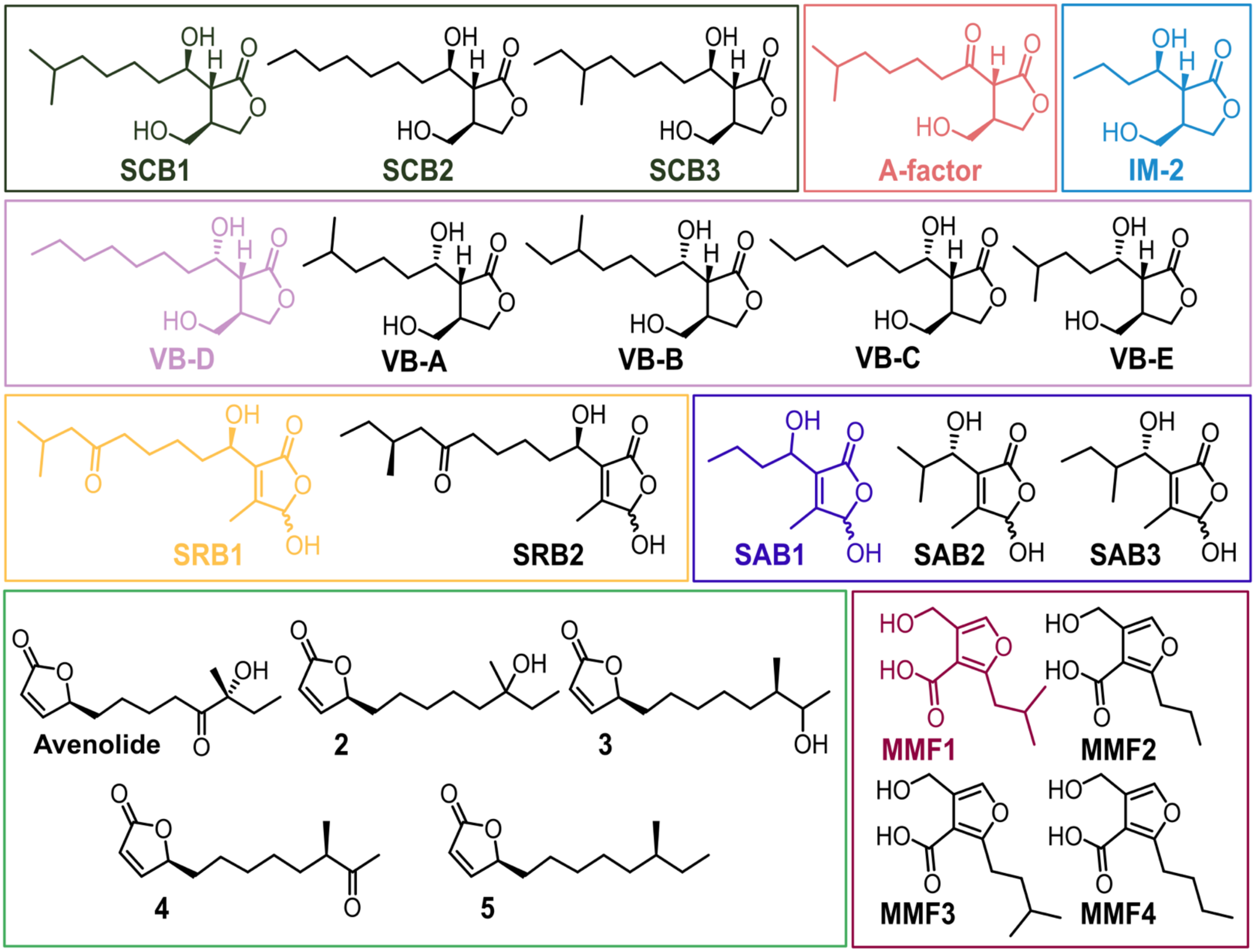

**<u>Supplementary Figure 5:</u> Natural *Streptomyces* sp. autoregulators.**
Natural variations in *Streptomyces* autoregulators occur as a result of the different β-keto ester precursors utilized during the first step of autoregulator synthesis. For example, the SCBs of *Streptomyces coelicolor* A3(2) have been identified in the isobutyl, n-butyl, and secbutyl derivatives as presented here. The colored derivatives indicate the structure of the autoregulator used in the synthetic molecule induction assay.

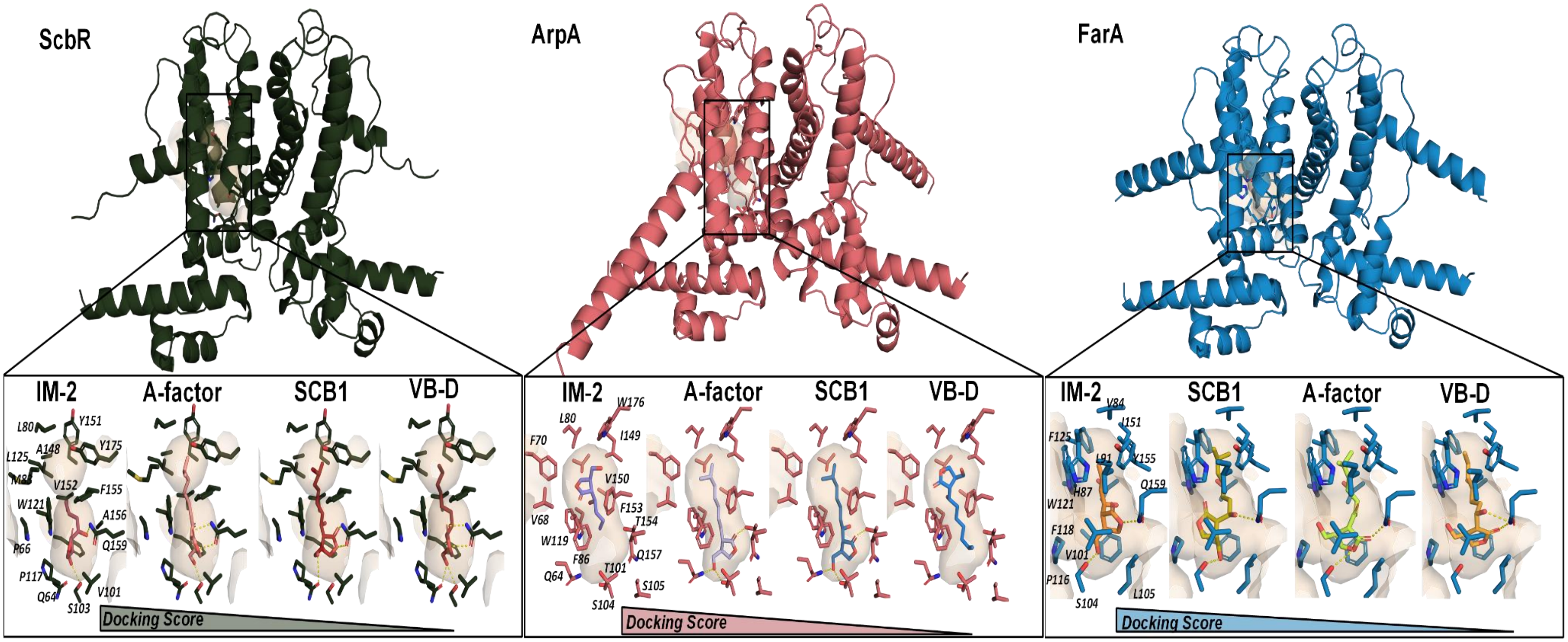

**<u>Supplementary Fig 6:</u> Results of docking autoregulators into CSRs.**

The gamma-butyrolactones SCB1, A-factor, VB-D, and IM-2 were docked into AlphaFold3 models of the three gamma-butyrolactone receptor CSRs: ScbR, ArpA, and FarA. Although IM-2 did not induce GFP expression (i.e. derepression or repression inhibition) for ScbR or ArpA, it was the molecule with the best docking score, suggesting that docking score is not reflective of bioactivity. For FarA, VB-D was ranked lower than both SCB1 and A-factor although it was the only molecule amongst the three that was active. Notably, SCB1 was ranked third overall in binding to its cognate CSR ScbR. All of this data indicates molecular docking is not yet an effective tool for accurately identifying biologically active *Streptomyces* CSR/autoregulator pairs.

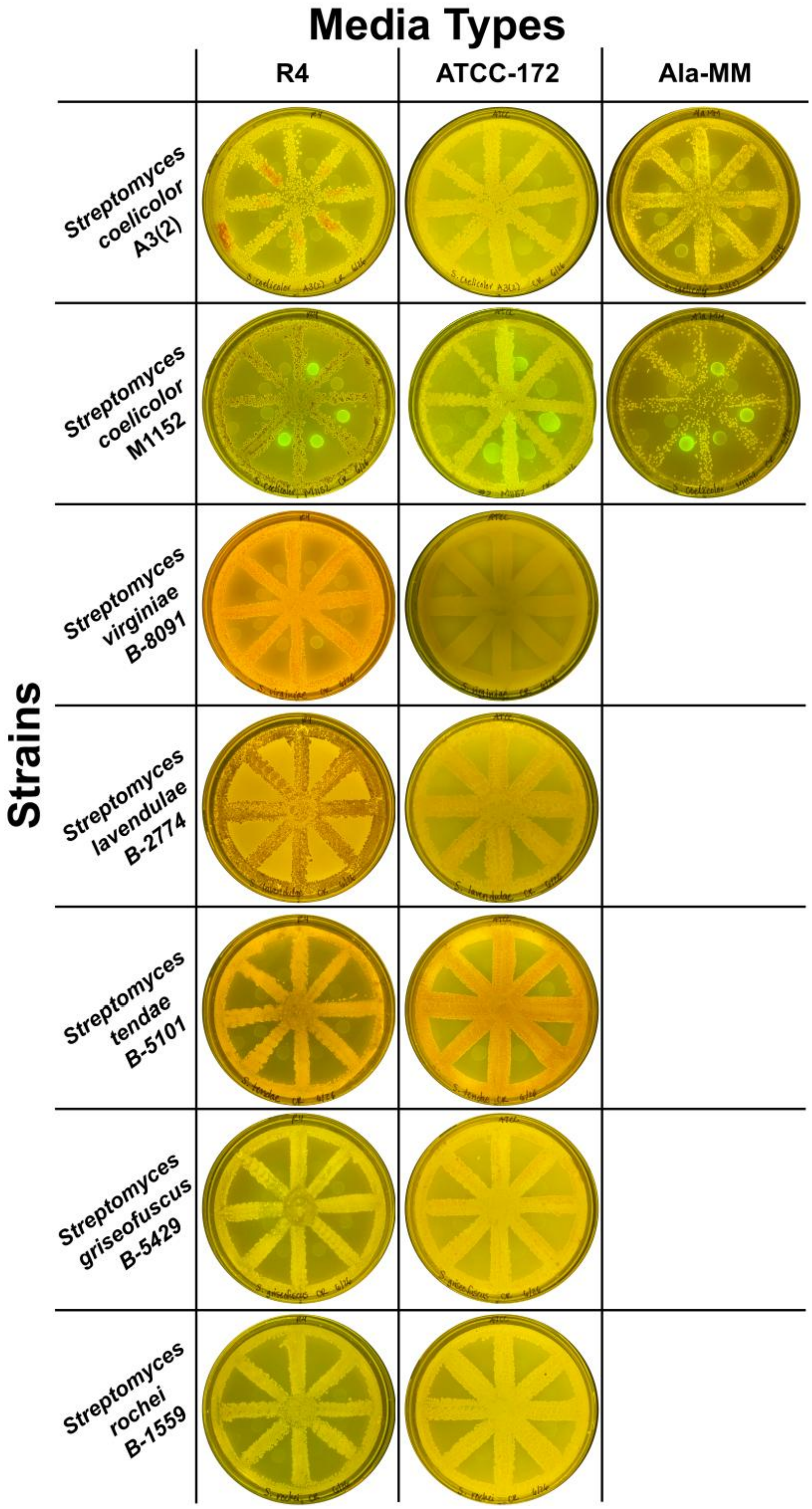

**<u>Supplementary Figure 7:</u> *Streptomyces* and *E. coli* based biosensor coculture on solid media.** Plates are grouped horizontally by *Streptomyces* strain and vertically by media used. For certain CSRs, their *Streptomyces* strain of origin is not commercially available. For these *Streptomyces*, we selected a related, commercially available strain that possessed an identical copy of the CSR within its genome. For FarA: *S. lavendulae* B-2774, SabR1: *S. tendae* B-5101, *S. griseofuscus* B-5429, and for SrrA: *S. rochei* B-1559. Signal was not observed from these combinations of CSR biosensors and *Streptomyces* during either the solid media coculture or the crude extract induction. The inability for the biosensor strains to fluoresce indicates the cognate autoregulator is not synthesized under the given condition (media, solid culture) or is not made in high enough quantities to activate GFP production.

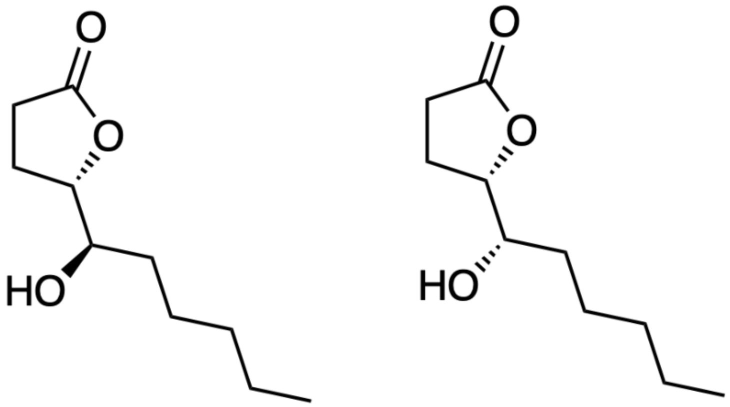

**<u>Supplementary Figure 8:</u> Additional autoregulators made by *S. griseus.***
Structures of two autoregulators made by *S. griseus* in addition to A-factor. As of yet, no cognate CSR has been identified for these molecules[9,21,22].

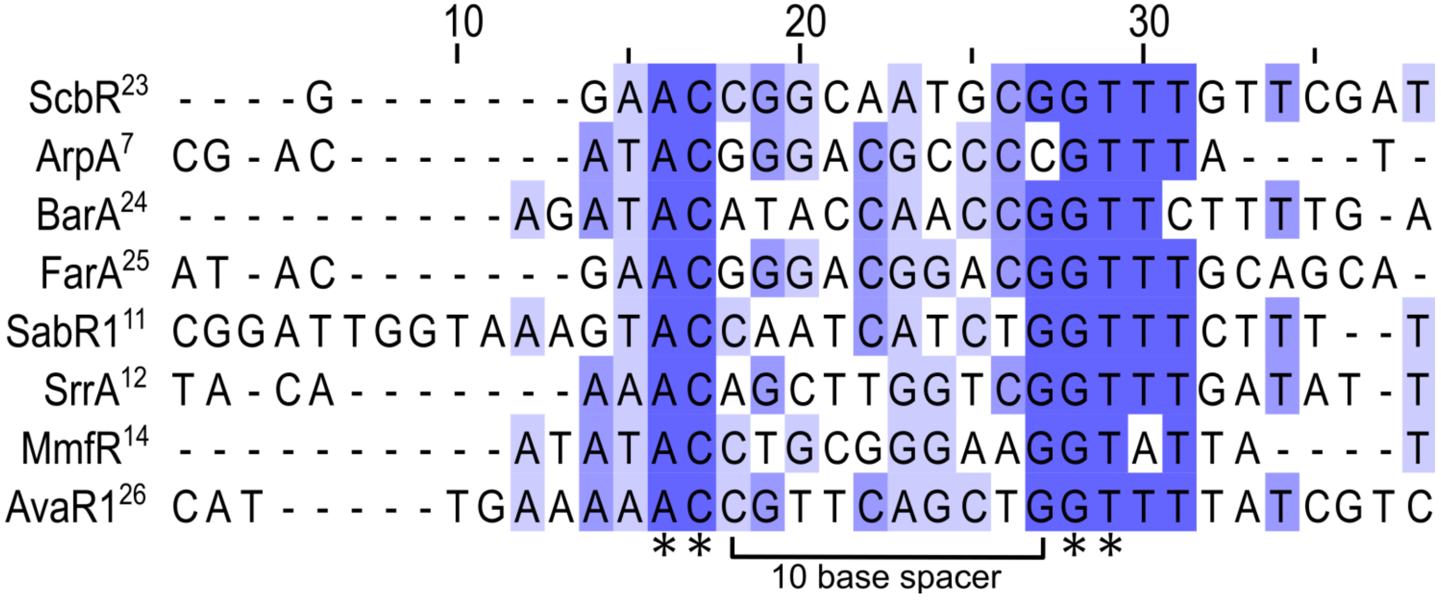

**<u>Supplementary Figure 9:</u> Alignment of native operator sequences.**
Sequences were sourced from indicated literature (superscript). All operator sequences possess a conserved 'AC-GT' motif with an unconserved 10 base spacer between the conserved bases. Interestingly, this spacer is much larger than those observed in other TetR-like repressors[27,28].

Cotransformed Fold Repression Average

| | $ScbR_R$ | $AvaR1_R$ | $ArpA_R$ | $BarA_R$ | $FarA_R$ | $SabR1_R$ | $MmfR_R$ | $SrrA_R$ |
|---|---|---|---|---|---|---|---|---|
| $ScbR_O$ | 67.33 | 37.60 | 6.26 | 25.26 | 73.14 | 2.23 | 1.07 | 39.41 |
| $AvaR1_O$ | 3.61 | 9.51 | 2.14 | 3.00 | 25.03 | 10.63 | 5.21 | 6.27 |
| $ArpA_O$ | 2.66 | 6.39 | 22.80 | 9.10 | 27.07 | 1.90 | 0.88 | 1.47 |
| $BarA_O$ | 4.88 | 10.73 | 27.06 | 63.57 | 82.81 | 2.47 | 1.49 | 32.35 |
| $FarA_O$ | 9.04 | 22.09 | 44.77 | 54.93 | 80.58 | 1.87 | 0.98 | 10.01 |
| $SabR1_O$ | 0.93 | 0.96 | 0.82 | 1.24 | 0.87 | 1.47 | 0.94 | 1.00 |
| $MmfR_O$ | 12.94 | 13.35 | 3.47 | 1.37 | 28.44 | 1.93 | 103.80 | 14.61 |
| $SrrA_O$ | 97.05 | 51.03 | 99.12 | 144.67 | 134.36 | 3.03 | 1.57 | 65.00 |

pGFP Max RFU Average

| | pGFP Max RFU Average |
|---|---|
| $ScbR_O$ | 92758.22 |
| $AvaR1_O$ | 88520.73 |
| $ArpA_O$ | 67166.08 |
| $BarA_O$ | 55079.07 |
| $FarA_O$ | 100996.28 |
| $SabR1_O$ | 82900.13 |
| $MmfR_O$ | 75053.38 |
| $SrrA_O$ | 114205.64 |

Cotransformed Fold Repression Standard Deviation

| | $ScbR_R$ | $AvaR1_R$ | $ArpA_R$ | $BarA_R$ | $FarA_R$ | $SabR1_R$ | $MmfR_R$ | $SrrA_R$ |
|---|---|---|---|---|---|---|---|---|
| $ScbR_O$ | 11.70 | 3.55 | 1.39 | 1.89 | 3.32 | 0.65 | 0.09 | 5.84 |
| $AvaR1_O$ | 0.64 | 0.82 | 0.65 | 0.48 | 3.13 | 2.63 | 0.70 | 0.73 |
| $ArpA_O$ | 0.57 | 0.81 | 2.41 | 1.71 | 4.30 | 0.60 | 0.09 | 0.12 |
| $BarA_O$ | 0.84 | 1.15 | 2.96 | 4.10 | 1.66 | 0.52 | 0.18 | 4.35 |
| $FarA_O$ | 1.46 | 4.17 | 10.47 | 9.30 | 10.61 | 0.33 | 0.16 | 2.24 |
| $SabR1_O$ | 0.08 | 0.06 | 0.12 | 0.11 | 0.05 | 0.14 | 0.03 | 0.09 |
| $MmfR_O$ | 2.22 | 1.42 | 0.46 | 0.10 | 4.58 | 0.55 | 3.82 | 0.97 |
| $SrrA_O$ | 13.46 | 3.82 | 23.22 | 16.86 | 4.60 | 0.85 | 0.19 | 17.85 |

pGFP Max RFU SD

| | pGFP Max RFU SD |
|---|---|
| $ScbR_O$ | 1189.67 |
| $AvaR1_O$ | 6494.56 |
| $ArpA_O$ | 4875.60 |
| $BarA_O$ | 14757.45 |
| $FarA_O$ | 18627.15 |
| $SabR1_O$ | 7172.16 |
| $MmfR_O$ | 8580.44 |
| $SrrA_O$ | 7047.92 |

**Supplementary Figure 10: Average and standard deviation of fold repression of cotransformed pREP/pGFP vectors.**

For 'Cotransformed Fold Repression Average' and 'Cotransformed Fold Repression Standard Deviation', labels across the horizontal axis (i.e. $ScbR_R$) indicate the CSR constitutively expressed by the pREP vector. Vertical labels indicate which operator is within the *gfp* promoter of the pGFP (i.e. $ScbR_O$). Fold repression is calculated by dividing the RFU of the cotransformed culture by the RFU of the culture that only possesses the pGFP vector. The 'pGFP Max RFU Average' and 'pGFP Max RFU SD' are the average and standard deviation of the pGFP vector that possesses each operator in the *gfp* promoter. Note that for the pREP/pGFP cotransformation experimentation, the pGFP plasmid possesses a ColE1 origin (~25 copy number), whereas the pTOTAL plasmids have a p15A origin of replication (~5 copy number).

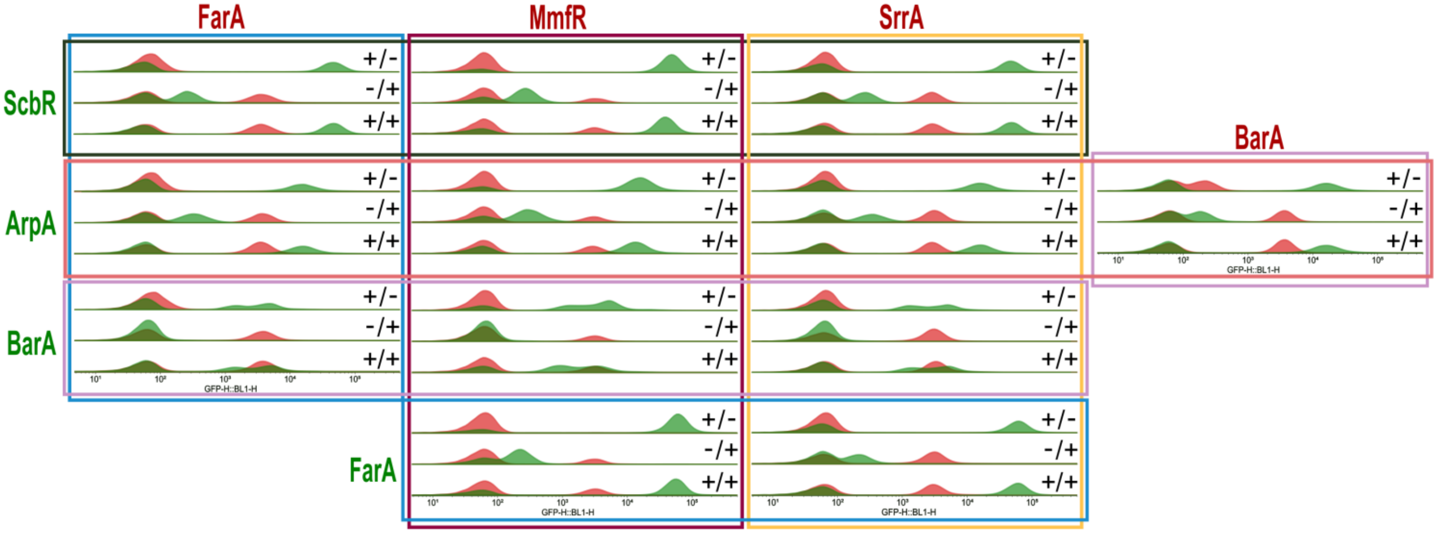

**<u>Supplementary Figure 11:</u> Histogram of biosensor coculture results.**
Cocultures of pTOTAL-CSR-GFP and pTOTAL-CSR-mCherry were induced with either the GFP-inducing autoregulator, the mCherry-inducing autoregulator, or both autoregulators. Vertical, green CSRs represent the biosensor vector in the coculture that expresses GFP when induced. Horizontal, red CSRs represent the biosensor vector in the coculture that expresses mCherry when induced. Each histogram plot was generated by combining GFP (green) and mCherry (red) fluorescence results of a single induced coculture sample. For each coculture the top histogram represents the GFP and mCherry results of the coculture replicate that was induced with the GFP-inducing autoregulator (+/-). Middle histogram represents results of the coculture sample that was induced with the mCherry-inducing regulator (-/+). Bottom histogram represents the results of the coculture sample that was indued with both autoregulators (+/+).

### Biology Materials and Methods

Media, *E. coli* Strains, Cloning Conditions

All *E. coli* and *Streptomyces* media utilized in this work and discussed below are listed here: **(1) Luria-Bertani (LB) Media** (10 g/L tryptone, 5 g/L yeast extract, 10 g/L NaCl); **(2) Supplemented M9 Media** (0.241 g/L $MgSO_4$, 0.111 g/L $CaCl_2$, 2 g/L casamino acids, 0.337 g/L thiamine hydrochloride, 4 g/L glucose, 100 mL 10X M9 salts [64 g/L $Na_2HPO_4$•7H2O, 15 g/L $KH_2PO_4$, 2.5 g/L NaCl, 5.0 g/L $NH_4Cl$]); **(3) ATCC-172 Media** (10 g/L glucose, 20 g/L soluble starch, 5 g/L yeast extract, 5 g/L N-Z amine Type A, 1 g/L $CaCO_3$, 15 g/L agar, adjusted to pH 7.3); **(4) R4 Media** (5 g/L glucose, 1 g/L yeast extract, 5 g/L $MgCl_2$•6$H_2O$, 2 g/L $CaCl_2$•2$H_2O$, 1.5 g/L L-proline, 1.2 g/L L-valine, 2.8 g/L TES, 50 mg/L casamino acid, 100 mg/L $K_2SO_4$, 15 g/L agar and 1 mL 1x trace element solution [40 mg/L $ZnCl_2$, 200 mg/L FeCl3•6$H_2O$, 10 mg/L $CuCl_2$•2$H_2O$, 10 mg/L $MnCl_2$•4$H_2O$, 10 mg/L $Na_2B_4O_7$•10$H_2O$, 10 mg/L $(NH_4)_6Mo_7O_{24}$•4$H_2O$], adjusted to pH 7.0); **(5) Alanine-Minimal Media (Ala-MM)** (2.67 g/L L-alanine, 871 mg/L $K_2HPO_4$, 1.23 g/L $MgSO_4$•7$H_2O$, 10 g/L glycerol, and 15 g/L agar, adjusted to pH 5.0).

All primers, plasmids, strains and DNA sequences are present in **Methods Tables 1-3**. During the cloning process, NEB® 5-alpha Chemically Competent *E. coli* cells (NEB, C2987H) were used with NEB's 'High Efficiency Transformation Protocol' for cloning transformation and plasmid propagation. Cells were grown in LB broth with kanamycin and carbenicillin, or in combination when appropriate. *E. coli* strain DH10B (Thermo Scientific, EC0113) was used for all fluorescence assays; transformations into this strain were performed according to manufacturer's protocol. All PCR amplification was performed using Q5 High-Fidelity 2x Master Mix (NEB, M0492S) with the following general methodology, adjusting the annealing temperature and the amplification time in accordance with the manufacturer's protocols. For each amplicon, triplicate PCR reactions were performed. Each of the three PCR tubes receives 10 µL of 2x Q5 Master Mix, 2 µL of mixed F/R primers at 500 µM each, 1 µL of 1 ng/µL plasmid template, and 7 µL of nuclease free water. Tube 1 was amplified at the annealing temperature recommended by NEBs TM calculator for Q5. Tube 2 received 3% DMSO and was amplified at the recommended annealing temperature. PCR for tube 3 without DMSO was performed using the stepdown/touchdown PCR method. The annealing temperature was set 2 °C over manufacturer guidelines and the thermocycler was programed to reduce the annealing temperature by 0.3 °C every round for 24 rounds of PCR. Thermocyclers were operated under the following general program: (1) initial denaturation at 98 °C for 30s; (2) 98 °C for 20s; (3) 60-70 °C for 15s (stepdown when appropriate) (4) 72 °C (template dependent, 20 sec per 1kb); (5) repeat steps 2–4 for a total of 25 cycles; (6) 72 °C for 2 min and (7) hold at 4 °C. All reactions were run on a 0.8% agarose gel with the GeneRuler 1kb plus ladder (Thermo Scientific, SM1331); successful amplification were gel excised, pooled, and DNA was extracted using the Freeze 'N Squeeze™ DNA Gel Extraction Spin Columns (Bio-Rad, 7326165) adhering to manufacturer protocol. DNA from the spin column was aliquoted into ice cold 900 µL 100% EtOH with 30 µL 3M Sodium acetate for ethanol precipitation. 1 µL of Glycoblue (Invitrogen, AM9516) was added to the mix for DNA visualization. The DNA precipitation mix was centrifuged at 15,000 rpm, washed twice with 700 µL of 70% EtOH, then the DNA pellet was resuspended in nuclease free water. For isothermal assembly, a 3:1 molar backbone to insert ratio was used with NEBuilder® HiFi DNA Assembly Cloning

Kit (NEB, E5520S). When parts were +/- 500bp of each other, a 1:1 ratio was used. For restriction digest cloning, plasmids were digested according to manufacturer protocol (New England Biolabs). Briefly, 1 µg plasmids or inserts were combined with 5 µL 10X CutSmart buffer, 1 µL of each enzyme, and nuclease free water was added for a total volume of 50 µL. These reactions were incubated for 1 hour at 37 °C, then separated by DNA gel electrophoresis and gel extracted as previously described. The T4 ligation reaction was prepared according to manufacturer protocol (NEB, M0202S), except that a 5:1 insert to backbone ratio was used for the ligation reaction. DNA for blunt end ligation was amplified and isolated using the aforementioned protocols; reactions were performed combining 150 ng of template, 1 µL DpnI (NEB, R0176S), 1 µL T4 ligase (NEB, M0202S), 1 µL T4 PNK (NEB, M0201S), 1 µL 10x T4 ligase reaction buffer (NEB, M0202S), and $H_2O$ for a total of 10 µL. Independent of cloning strategy, transformations were performed according to manufacturer protocol for chemically competent NEB® 5-alpha (NEB, C2987H). All clones were screened by restriction digest, then confirmed by Sanger sequencing (Azenta).

pTOTAL Vector Cloning

pLW0003 from our previous publication was used as the initial vector backbone (**Table 3**)[29]. Primers 1/2 (**Table 1**) amplified pLW0003 to generate a backbone to replace chloramphenicol resistance with kanamycin resistance. The kanamycin resistance was amplified from pET28b with primers 3/4. PCR reaction, gel extraction, clean up, isothermal assembly, and transformation were performed as described previously. The resulting vector was amplified with primers 5/6 to change the origin of replication to p15A. p15A insert was amplified from plasmid pRF-ScbR (**Table 3**) with 7/8. PCR reaction, gel extraction, clean up, isothermal assembly, transformation, and validation were performed as described previously.

From the resulting mother plasmid, CSR vectors were made in three sequential steps. First, the promoter region of GFP was changed to possess the BBa_J23100 promoter -10/-35 sequence, a CSR operator, the BydvJ ribozyme sequence, and RBS BBa_J34801. A GFP promoter dsDNA fragment was ordered for each of the operator from Integrated DNA Technologies (IDT) (**Table 2**). Backbone and GFP promoter dsDNA fragments were digested with restriction enzymes SalI and NdeI, then ligated and transformed as described. Second, the repressor promoter region was cloned by digesting the previously generated plasmid and a purchased CSR promoter DNA fragment (IDT) with EcoRI and AatII as described. Third, the cognate CSR coding sequence was cloned into the backbone containing its operator. The CSR sequence was ordered from IDT with homologous overhangs for isothermal assembly. For amplification of backbone to clone in CSR, forward primer 9 is used with primer 10, 11, or 12 depending on CSR. PCR reaction, gel extraction, clean up, isothermal assembly, and transformation were performed as described. For the AvaR1, ArpA, and BarA vectors the BydvJ ribozyme sequence was removed by amplifying the vector from step three with primers depending on the operator (14 for $AvaR1_O$, 15 for $ArpA_O$, 16 for $BarA_O$). Electrophoresis, gel extraction, and blunt-end ligation were performed as described above.

Synthetic Molecule Induction Assay Protocol

A single colony was used to inoculate a 14 mL disposable culture tube (USA Scientific, #5618-7262) containing 5 mL LB media with 50 µg/mL kanamycin; a vectorless DH10B control culture was also started in LB without antibiotics. The cultures were incubated for 16-18 hours at 37 °C in a Cole-Parmer TR-200D-120 Tube Rotator with rotation (40 rpm). The overnight cultures were diluted 1:100 into supplemented M9 and 198 µL of the cultures were dispensed into the wells of a round bottom Corning Clear Polystyrene 96-Well Microplate (Corning, 3797) and covered with AeraSeal film (Sigma Aldrich, A9224-50EA). The cultures were grown on an ELMI DTS-4 Digital Thermo Shaker at 1000 rpm, 37 °C to an $OD_{600}$ of 0.1-0.2 (approximately 3-4 hours) and induced with 2 µL (1% v/v) of each respective molecule or DMSO vehicle. After induction, the cultures were covered with a new AeraSeal film, and the plate was incubated under the same conditions for an additional 5 hours. After incubation, 1 µL of each culture was diluted into the wells of a Corning Clear Polystyrene 96-well microplate (Corning, 9017) filled with 199 µL cold, filter-sterilized PBS with 2 mg/mL streptomycin to arrest cell growth. These culture dilution plates were then vortexed thoroughly and analyzed using a ThermoFisher Attune NxT Flow Cytometer. Culture dilution plates that were not immediately analyzed were stored at 4 °C and then vortexed for 1 minute before flow cytometry analysis.

Flow Cytometry Analysis

The following protocol was used for analyzing flow cytometry data for the synthesized molecule induction, crude extract induction, and pREP/pGFP cotransformation assays. An injection volume of 50 µL and a flow rate of 25 µL/min were used, and 18,000 events were collected. This protocol was adjusted for the pTOTAL-CSR *E. coli* coculture assay (see ‘pTOTAL-CSR *E. coli* Coculture Cloning, Protocol, and Analysis’). Events were gated on a range of X axis FSC-W 1 to 100 using floreada.io, and mean fluorescence was calculated from at least 10,000 events. The mean autofluorescence of a DH10B vector-empty control was subtracted from each fluorescent mean. Fold change for each pTOTAL-CSR culture was calculated by dividing the maximum induced fluorescence from each biological replicate by the fluorescence of an average of technical replicates of DMSO supplemented pTOTAL-CSR culture, then averaging the fold changes of each biological replicate. Data presented is an average of biological triplicate collected on separate days, error bars are +/1 standard deviation. $K_D$ and Hill coefficients were determined by fitting the data to a Hill function below using a weighted least of squares adjustment.

$$f(x) = y = Y_{min} + (Y_{max} - Y_{min}) \frac{x^n}{x^n + K_D^n}$$

pREP/pGFP Vector Cloning and Operator Crosstalk Analysis

To generate the pREP vectors for the operator crosstalk analysis, each of the pTOTAL vectors were amplified with primers 17/18 and blunt-end ligated; this removed the GFP cassette from the pTOTAL backbone. To generate the pGFP vectors a three-part assembly was performed: the ColE1 origin of replication was generated by amplifying the pTU2S-a plasmid with primers 19/20, ampicillin resistance was amplified from pGEX-6P-1 with primers 21/22; and the GFP cassette including promoter, operator, and *gfp* was amplified from each pTOTAL vector using primers 23/24. All cloning was performed as described above. After verification of the pREP and pGFP plasmids, each pREP vector (8 total) was cotransformed with every pGFP vector (8 total) to create 64 unique combinations. For this transformation, one microliter of each plasmid at ~50 ng/µL was added to 50 µL of competent cells, all other steps were according to manufacturer protocol.

Each cotransformed strain was streaked from a pure culture to LB with 50 µg/mL kanamycin and 100 µg/mL carbenicillin, pREP cultures were streaked to LB with 50 µg/mL kanamycin and pGFP cultures were streaked to LB with 100 µg/mL carbenicillin. A single culture from each of the 64 cotransfomed strains, 8 pREP, and 8 pGFP were inoculated into 1 mL of supplemented M9 with respective antibiotics in a 96-well V-bottom deep well plate (Stellar Scientific, IP-DP22VS-9-N). This plate was covered with an AeraSeal film and incubated at 16 hours at 30 °C on a Fisherbrand 4-Place Platform Shaker (FisherScientific, 88-861-023) shaking at 1000 rpm. After incubation, each culture was diluted 1:4, then further diluted 1:200 in cold, filter-sterilized PBS with 2 mg/mL streptomycin to arrest cell growth. Culture dilution plates were then vortexed thoroughly and analyzed using a ThermoFisher Attune NxT Flow Cytometer.

<u>MSA and Alpha-Fold Model Generation</u>

Amino acid sequences were aligned using the default T-coffee multiple sequence alignment[30]. Alpha Fold models of the dimers for ScbR, ArpA, and FarA using the standard settings for AlphaFold3[31]. For the docking simulation, the AlphaFold3 structures of proteins were refined using the Protein Preparation Wizard in Maestro (Schrödinger Suite 2024-2, Schrödinger LLC), using PROPKA in the hydrogen bond assignment step with the pH set at 7.0. proteins were docked using Glide through standard precision (SP) mode. The center of the grid box for the simulation was set around the ligand binding domains and post-docking minimization was conducted for 200 poses. The top 5 poses (by docking score) for each docked ligand were chosen and analyzed using PyMOL.

<u>*Streptomyces/E. coli* Biosensor Plate Coculture</u>

*Streptomyces* strains (**Methods Table 3**) were streaked frozen glycerol stocks to solid ATCC-172. Plates were grown inverted at 30 °C in a closed container for 5 days with MilliQ water available for humidity. After 5 days of growth on solid media, one colony of each *Streptomyces* strain was used to inoculate a glass culture tube with 5mL of liquid ATCC-172. These tubes were incubated at 30 °C on a New Brunswick TC-7 Tissue Culture Roller Rotator (40 rpm) for 3 days. After 3 days of growth, a sterile cotton-tipped swab was used to inoculate solid agar plates in the "sunburst" pattern on both ATCC-172 and R4 plates for all strains. *Streptomyces* sp. that produce MMFs were also plated on Ala-MM. These were grown inverted at 30 °C for 2 days. 5 mL overnights of each pTOTAL CSR biosensor (ScbR, AvaR1, ArpA, BarA, FarA, SabR1, MmfR, and SrrA) and a vectorless DH10B control were started in LB in disposable 14 mL polypropylene culture tubes, with 50 µg/mL kanamycin for the biosensor cultures. On day 2 of the *Streptomyces* sunburst plate growth, 20 µL of each biosensor vector culture were aliquoted on to a 1/8th section of the plate and allowed to dry briefly in a biosafety cabinet. Plates were grown inverted overnight for approximately 16-18 hours at 30 °C in a closed container with

MilliQ water for humidity. Plates were visualized with UV transillumination; pictures are a representation of one of three biological replicates performed.

*Streptomyces* Crude Extract Isolation

*Streptomyces* strains (**Methods Table 3**) were started from frozen glycerol stocks streaked to solid ATCC-172. Plates were grown inverted at 30 °C in a closed container for 5 days with MilliQ water available for humidity. A single colony was transferred to 5mL liquid broth culture of ATCC-172; these tubes were incubated at 30 °C on a New Brunswick TC-7 Tissue Culture Roller Rotator for 7 days (40 rpm). Then, 750 µL of culture was plated per plate using glass beads to create a dense lawn. Each strain was cultivated on 15 solid agar plates. These plates were grown for 5 days at 30 °C in the same conditions described above, sliced into approximately 1 $cm^3$ cubes with a razor blade and soaked in 350-400 mL ethyl acetate overnight. The overnight extraction was filtered through filter paper into a round bottom flask and dried via rotary evaporation until no solvent was visible, then placed under high vacuum for 30 minutes for removal of trace solvent. The crude extract was weighed then resuspended in DMSO to 100 mg/mL

Crude Extract Induction Assay

For the crude extract induction assay, a single isolated colony from each pTOTAL vector was used to inoculate 5 mL LB with 50 µg/mL kanamycin; a 5 mL vectorless DH10B control was started in LB without antibiotics. The overnight liquid culture was incubated for 16-18 hours at 37 °C in a Cole-Parmer TR-200D-120 Tube Rotator with rotation (40 rpm). The overnight culture was diluted 1:100 into supplemented M9 media and 990 µL was dispensed into 5 mL Falcon™ Round-Bottom Polystyrene Test Tubes. These tubes were incubated under the same conditions as the overnights. At $OD_{600}$ 0.1-0.2, each tube was inoculated with 10 µL of the respective crude extract for a final concentration of 1 mg/mL; 10 µL DMSO was added to the control culture tubes. The tubes were returned to the incubator and grown for an additional 5 hours under the same conditions. After induction, 1 µL of each culture was diluted into 199 µL cold, filter-sterilized PBS with 2 mg/mL streptomycin to arrest cell growth, vortexed, then analyzed using a ThermoFisher Attune NxT Flow Cytometer according to the protocol in ‘Flow Cytometry Analysis’.

pAND-SrrA.MmfR Cloning and Protocol

The pTOTAL-SrrA plasmid was used as the starting backbone for cloning the pAND-SrrA.MmfR vector. Primers 25/26 were used to amplify pTOTAL-SrrA to add the $MmfR_O$ upstream of the *gfp* promoter. This linear DNA was ligated using blunt end ligation as previously described. The resulting plasmid was then amplified with primers 27/28 to allow for insertion of the MmfR cassette; the MmfR cassette to be inserted was amplified with primers 29/30. Isothermal assembly, transformation, and verification were all performed as previously described.

For the assay, a single isolated pAND-SrrA.MmfR colony was used to inoculate 5 mL LB with 50 µg/mL kanamycin, along with a 5 mL overnight in LB without antibiotics for a vectorless DH10B control. The overnight liquid cultures were incubated for 16-18 hours at 37 °C in a tube rotator with rotation (40 rpm). Overnight cultures were diluted 1:100 into supplemented M9 media and 198 µL of experimental and control cultures were dispensed into a round bottom clear polystyrene 96-well microplate. The cultures were grown on an ELMI shaker at 1000 rpm, 37 °C to an $OD_{600}$ of 0.1-0.2 (approximately 3-4 hours) and induced with 2 µL of each respective individual molecule, molecule combination, or vehicle. The pAND-SrrA.MmfR cultures were induced with a final concentration of 500 µM MMF1, 7.8 µM SRB1, or a combination of those molecules at the same final concentration in culture. After induction, the cultures were returned to the shaker plate incubated with the same conditions for an additional 5 hours. 1 µL of each culture was diluted into 199 µL cold, filter-sterilized PBS with 2 mg/mL streptomycin to arrest cell growth, vortexed, then analyzed using a ThermoFisher Attune NxT Flow Cytometer according to the protocol in 'Flow Cytometry Analysis'.

pTOTAL-CSR *E. coli* Coculture Cloning, Protocol, and Analysis

Using the pTOTAL vectors as template, the *gfp* coding sequence was replaced with mCherry. pTOTAL-BarA, pTOTAL-FarA, pTOTAL-MmfR, pTOTAL-SrrA were each amplified with primers 31/32 and the mCherry coding region was amplified from vector pRSET-B mCherry (**Methods Table 3**) with primers 33/34. Isothermal assembly, transformation, and validation were performed according to previous protocol, generating pTOTAL-FarA-mCherry, pTOTAL-BarA-mCherry, pTOTAL-MmfR-mCherry, pTOTAL-SrrA-mCherry.

For the analysis, a single colony of all mCherry pTOTAL vectors, as well as the GFP-containing vectors (denoted with '-GFP' for this protocol), pTOTAL-ScbR-GFP, pTOTAL-ArpA-GFP, pTOTAL-BarA-GFP, pTOTAL-FarA-GFP and DH10B controls were started in 5 mL LB, with 50 µg/mL kanamycin for experimental samples. The cultures were incubated for 16-18 hours at 37 °C in a Cole-Parmer TR-200D-120 Tube Rotator with rotation (40 rpm). All overnight cultures were diluted 1:100 into supplemented M9 media and each experimental combination of pTOTAL GFP and pTOTAL mCherry cultures were mixed 1:1. 198 µL of each coculture, as well as each individual pTOTAL-GFP/ pTOTAL-mCherry culture, was dispensed into the wells of a round bottom clear polystyrene 96-well microplate and covered with AeraSeal film. The cultures were grown on an ELMI Shaker at 1000 rpm, 37 °C to an $OD_{600}$ of 0.15-0.25 (approximately 4 hours) and each coculture replicate was induced with 2 µL of a single molecule, a molecule combination, or DMSO vehicle. For all cocultures with pTOTAL-ScbR-GFP, a final concentration of 31.5 µM of SCB1 was used as the induction concentration for both the single molecule (SCB1 alone) and dual molecule (SCB1+ a second inducer) conditions. For pTOTAL-ArpA-GFP 0.78 µM A-factor, for pTOTAL-BarA (both GFP and mCherry vectors) 6.25 µM VB-D, for FarA (both GFP and mCherry vectors) 0.625 µM IM-2, for pTOTAL-MmfR-mCherry 500 µM MMF1, and for pTOTAL-SrrA-mCherry 7.8 µM SRB1. After induction, the cultures were covered with a new AeraSeal film and the plate was incubated under the same conditions for an additional 5 hours. After incubation, 1 µL of each culture was diluted into the wells of a clear polystyrene 96-well microplate filled with 199 µL cold, filter-sterilized PBS with 2 mg/mL streptomycin to arrest cell growth. These culture dilution plates were then vortexed thoroughly and analyzed using a ThermoFisher Attune NxT Flow Cytometer using the previously described protocol, only increasing the number of events analyzed to 40,000. For flow cytometry data analysis, the events were first gated on a range of X axis FSC-W 1 to 100. To minimize the baseline signal from the *E. coli* strain in the coculture expressing the other fluorophore, gates were created to calculate the mean signal only including the induced cells. To calculate the fold change, the mean induced signal was divided by the mean fluorescence of corresponding the pTOTAL-GFP or pTOTAL-mCherry culture grown alone. Data presented is an average of biological triplicate collected on separate days, error bars are +/1 standard deviation.

**Methods Table 1: Primers**

| Primer Use | Primer Number | Sequence (5' -> 3') |
|---|---|---|
| **pTOTAL Cloning** | | |
| Amplify pLW0003 to replace chloramphenicol with kanamycin resistance | 1 | tttgatatcgagctcgcttgg |
| | 2 | gagtgccacctgacgtc |
| Amplification of kanamycin cassette from pET28b(-) | 3 | gaaccgtaaaaaggccgcgttgctggcgtttttccataggctccgcc |
| | 4 | agatcaaaggatcttcttgagatcctttttt |
| Amplify pLW0003 for origin replacement with p15a | 5 | gaagatcctttgatcttttctacggg |
| | 6 | aaacgccagcaacgc |
| Amplify p15a origin | 7 | gaaccgtaaaaaggccgcgttgctggcgtttttccataggctccgcc |
| | 8 | tgaccccgtagaaaagatcaaaggatcttcttgagatcgttttggtctgcg |
| Amplify backbone to replace repressor sequence Forward | 9 | gaattctttctcctcttttctagtttaaac |
| Reverse for ArpA, BarA insertion | 10 | taataagcggccgccac |
| Reverse for ScbR, FarA, SabR1, MmfR, or SrrA insertion | 11 | gaattctttctcctctttggatccg |
| Reverse for AvaR1 insertion | 12 | gaattcctccttcttaaagttaaacaaaattatttctagacc |
| Remove BydvJ Forward | 13 | taaagaggagaaatagcatatgcgtaaagg |
| Remove BydvJ for AvaR1 Reverse | 14 | ctagaagcttgacgataaaaccagc |
| Remove BydvJ for ArpA Reverse | 15 | ctagaagcttataaacggggcgt |
| Remove BydvJ BarA Reverse | 16 | caaccggttcttttgaaagcttcta |
| **pREP/pGFP Cloning** | | |
| Remove GFP cassette from pTOTAL, generate pREP | 17 | taatctagacactgatagtgctagtg |
| | 18 | gtcgacacctcttaagaggtaag |
| Amplify ColE1 origin of replication for pGFP | 19 | atttccccgaaaagtgccacctgatttatacccgggtactagaggtc |
| | 20 | atgataataatggtttcttattgagatcctttttttctgcg |
| Amplify amp/carb resistance for pGFP | 21 | aaggccgcgttgctggcgttttaccaatgcttaatcagtgagg |
| | 22 | gtacccgggtataaatcaggtggcacttttcgg |
| Amplify GFP cassette from pTOTAL for pGFP | 23 | gcagaaaaaaaggatctcaataagaaaccattattatcatgacattaacc |
| | 24 | aaacgccagcaacgc |
| **pAND-SrrA.MmfR Cloning** | | |
| Amplify pTOTAL-SrrA to add $MmfR_O$ upstream of promoter | 25 | atatacctgcgggaaggtattatttgacggctagctcagtc |
| | 26 | gtcgacacctcttaagaggtaag |
| Amplify pTOTAL-SrrA[$MmfR_O$] backbone to add in MmfR cassette | 27 | ctgcagccagaaatcatcgatgatttctggctgcagttgac |
| | 28 | tttataggtaccaatcaatctaaagtttataggttaatgtcatgataataatggtttcttagacg |
| Amplify MmfR cassette | 29 | tattatcatgacattaacctataaagtttactcatatatactttagattgattggtacctataaa |
| | 30 | ctgcagccagaaatcatcgatgatttctggctgcagtttacg |
| **pTOTAL – mCherry Cloning** | | |
| Amplify backbone of pTOTAL to swap GFP with mCherry | 31 | taatctagacactgatagtgctagtg |
| | 32 | atgctatttctcctctttagatctttaaac |
| Amplify mCherry coding region | 33 | aactagtaaagaggagaaatagcatatggtgagcaagggcga |
| | 34 | actagcactatcagtgtctagattattacttgtacagctcgtccatgc |

**<u>Methods Table 2:</u> DNA Sequences**

| CSR | Promoter Region of CSR | CSR Coding Sequence | GFP Promoter Region with CSR Operator |
|---|---|---|---|
| | AatII/EcoR1 | Homologous Overhang | SalI/NdeI |
| ScbR | ccacctgacgtctaa<br>gaaaccattattatc<br>atgacattaacctat<br>aaaaataggcgtat<br>cacgaggcagaatt<br>tcagataaaaaaaa<br>tccttagctttcgcta<br>aggatgatttctggct<br>gcagttgacggcta<br>gctcagtcctaggta<br>cagtgctagcggat<br>ccaaagaggagaa<br>agaattctggcca | ccaaagaggagaaagaattcATGGCCAAGCAGGACCGGGCGATC<br>CGCACGCGGCAGACGATCCTGGACGCCGCGGCGCAGGT<br>CTTCGAGAAGCAGGGCTACCAAGCTGCCACGATCACGGA<br>GATCCTCAAGGTGGCCGGGGTGACCAAGGGAGCCCTCTA<br>CTTCCACTTCCAGTCCAAGGAAGAACTGGCGCTGGGCGT<br>CTTCGACGCCCAGGAACCACCACAGGCCGTTCCGGAGCA<br>ACCCCTCCGGCTGCAAGAACTCATCGACATGGGCATGTTG<br>TTCTGTCACCGCTTGCGCACGAACGTCGTGGCCCGGGCC<br>GGCGTGCGCCTCTCCATGGACCAGCAGGCGCACGGTCTC<br>GATCGCCGAGGACCCTTCCGTCGCTGGCACGAGACACTC<br>CTGAAGCTGCTGAACCAGGCCAAGGAGAACGGTGAGTTG<br>CTGCCCCATGTGGTCACCACCGACTCGGCCGATCTCTAC<br>GTGGGCACGTTCGCCGGGATACAGGTCGTGTCCCAGACG<br>GTCAGCGACTACCAGGACCTCGAACACCGCTACGCGCTG<br>CTGCAGAAGCACATCCTGCCCGCCATCGCGGTTCCCTCC<br>GTGCTGGCCGCGCTCGATCTCTCCGAGGAGCGCGGAGCA<br>CGCCTCGCGGCCGAACTGGCACCGACCGGGAAGGACTG<br>Ataataagcggccgccactga | agaggtgtcgacttg<br>acggctagctcagtc<br>ctaggtacagtgcta<br>gcaggaaggaacc<br>ggcaatgcggtttgtt<br>cgataagcttagggt<br>gtctcaaggtgcgtac<br>cttgactgatgagtcc<br>gaaaggacgaaac<br>accctctacaaata<br>attttgtttaaagatcta<br>aagaggagaaatag<br>catatgcgtaaa |
| ArpA | ccacctgacgtctaa<br>gaaaccattattatc<br>atgacattaacctat<br>aaaaataggcgtat<br>cacgaggcagaatt<br>tcagataaaaaaaa<br>tccttagctttcgcta<br>aggatgatttctggct<br>gcagcgaggaagc<br>ggggccggccttga<br>cggctagctcagtcc<br>taggtacagtgtagc<br>ttgatcattataacag | aaagaggagaaagaattcATGGCGAAGCAGGCTCGCGCAGTCCA<br>GACGTGGCGGTCGATCGTGGATGCCGCGGCGAGTGTCTT<br>CGACGACTACGGCTACGAGCGTGCCGCCATCTCGGAGAT<br>TCTGCGCCGCGCCAAGGTCACCAAGGGGGCCTTGTACTT<br>CCACTTCGCCTCCAAGGAGGCCATCGCCCAGGCGATCAT<br>GGACGAGCAGACGTCCACGGTGGAGTTCGAGCAGGAGG<br>GCTCGCCGCTTCAGTCCCTGGTGGACGGGGGCCAGCAGT<br>TCGCTTTCGCCCTGCGCCACAACTCGATGGCCCGGGCCG<br>GTACCAGGCTCTCCATCGAGGGCGTCTTCCTCGGCGGGC<br>CGCACCCCTGGGGCGACTGGATCGACGCGACGGCCCGG<br>ATGCTGGAGCTGGGCCAGGAGCGCGGCGAGGTGTTCCC<br>GCAGATCGACCCGATGGTGTCAGCCAAAATCATCGTCGCT<br>TCGTTCACCGGTATCCAGCTGGTCTCGGAGGCCGATTCC | agaggtgtcgacttg<br>acggctagctcagtc<br>ctaggtacagtgcta<br>gcaggaacgacata<br>cgggacgccccgttt<br>ataagcttctagtaaa<br>gaggagaaatagca<br>tatggagcaa |

| | | | |
|---|---|---|---|
| | ctgtcaccggatgtgctttccggtctgatgagtccgtgaggacgaaacagcctctacaaataattttgtttaaactagaaaagaggagaaagaattctggcga | GGCCGGGCCGATCTCCGCGAGCAGGTGGCGGAGATGTGGCGGCACATCCTGCCGTCGATCGCCCACCCCGGGGTCATCGCCCACATCAAACCGGAGGGCCGGGTGGATCTGGCGGCCCAGGCGCGCGAGAAGGCCGAGCGTGAGGAGCAGGAGGCCAGGATCGCCGCGGAGGCCAAGGGGGCGGGCTCCGATCCCACGAGCGAGGGCGGCACCCGGTCGGGCGGGTCGGGGCTGCGGGGCGGGGGATCCGGCCGTGGTCCGCGCGCCGGGGTGACCGGCGACGAGGGCGACGAGGAGCCCGCAGGTGCCGGGGTGGCGGCCGGGGGGATCGTCGCCTGAtaataagcggccgccactgatagt | |
| BarA | gccactgacgtctaagaaaccattatatcatgacattaacctataaaataggcgtatcacgaggcagaatttcagataaaaaaatccttagctttcgctaaggatgatttctggctgcagcgaggaagcggggccggccttgacggctagctcagtcctaggtacagtgtagcttgatcattataacagctgtcaccggatgtgctttccggtctgatgagtccgtgaggacgaaacagcctctacaaataattttgtttaaactagaaaagaggagaaagaattctggcga | tagaaaagaggagaaagaattcATGGCAGTTCGTCATGAACGTGTTGCCGTGCGTCAAGAACGTGCAGTTCGTACCCGTCAGGCAATTGTTCGTGCAGCAGCAAGCGTTTTTGATGAATATGGTTTTGAAGCAGCAACCGTTGCAGAAATTCTGAGCCGTGCAAGCGTTACCAAAGGTGCAATGTATTTTCATTTTGCCAGCAAAGAAGAACTGGCACGCGGAGTTCTGGCAGAACAGACCCTGCATGTTGCAGTTCCGGAAAGCGGTAGCAAAGCACAAGAACTGGTTGATCTGACCATGCTGGTTGCACATGGTATGCTGCATGATCCGATTCTGCGTGCAGGCACCCGTCTGGCACTGGATCAGGGTGCAGTTGATTTTTCAGATGCAAATCCGTTTGGTGAATGGGGTGATATTTGTGCACAGCTGCTGGCCGAAGCGCAAGAACGCGGTGAAGTTCTGCCGCATGTTAATCCGAAAAAGACCGGTGATTTTATTGTGGGTTGTTTTACCGGTCTGCAGGCAGTTAGCCGTGTTACCAGCGATCGTCAGGATCTGGGTCATCGTATTAGCGTTATGTGGAATCATGTTCTGCCGAGCATTGTTCCGGCAAGCATGCTGACCTGGATTGAAACCGGTGAAGAACGTATTGGTAAAGTTGCAGCCGCAGCAGAAGCAGCCGAAGCCGCAGAAGCATCTGAAGCAGCCAGTGATGAATAATAAtaagcggccgccactgatag | cttaatgtcgacttgacggctagctcagtcctaggtacagtgctagcaggaaagatacataccaaccggttcttttgaaagcttctagtaaagaggagaaatagcatatggtacga |

| | | | |
|---|---|---|---|
| FarA | ccacctgacgtctaa<br>gaaaccattattatc<br>atgacattaacctat<br>aaaaataggcgtat<br>cacgaggcagaatt<br>tcagataaaaaaa<br>tccttagctttcgcta<br>aggatgatttctggct<br>gcagttgacggcta<br>gctcagtcctaggta<br>cagtgctagcggat<br>ccaaagaggagaa<br>agaattctggcca | atccaaagaggagaaagaattcATGGCTGAACAGGTCCGAGCCAT<br>CCGCACGCGCCAGGCGATCCTGAGCGCGGCGGCCAGGG<br>TGTTCGACGAACGGGGCTACCAGGCGGCCACCATCTCGG<br>AGATCCTCACCGTCGCGGGGGTGACCAAGGGCGCCCTGT<br>ACTTCCACTTCCAGTCCAAGGAAGACCTGGCCCAGGGCG<br>TCCTCACCGCGCAGAACGAGGATCTTCTGCTCCCTGAGC<br>GCCCCGCCAAACTCCAGGAGGTGGTGGATGCCGTCATGC<br>TGCACACCCACCGCCTGCGGACCAACCCCATGGTCCGCG<br>CGGGCGTCCGGCTGAGCCTCGACGTGAACGCCGGGGGG<br>CTCGACCGCAGCGCTCCGTTCCGCAACTGGGTCGACAAG<br>TTCACCGACCTCCTGGAGAAGGCCCAGGCCCAGGGCGAA<br>CTGCTGCCCCACGTGGTACCGGCCGAGACCGCCGACGTC<br>ATCACCGGTGCCTACGGCGGCGTCCAGTCCATGTCCCAG<br>GCGCTCACCGAGCACCAGGACCTCGGGCAGCGGGTCAA<br>CGCGCTGCTGCGCCACCTCATGCCGAGCATCGCCCAGCC<br>CTCGGTCCTCGCCTCCCTGCACCTCGGCGAGAGCCGGGC<br>GGAAGAGGTCTACCTCGAAGCCCGGCAGCTGGCCCGGG<br>AGCAGGCGGACGAGGAAGACTGAtaataagcggccgccactgatag<br>tg | agaggtgtcgacttg<br>acggctagctcagtc<br>ctaggtacagtgcta<br>gcaggaagatacga<br>acgggacggacggt<br>ttgcagcaaagctta<br>gggtgtctcaaggtg<br>cgtaccttgactgatg<br>agtccgaaaggacg<br>aaacacccctctaca<br>aataattttgtttaaag<br>atctaaagaggaga<br>aatagcatatgcgta<br>ag |
| SabR1 | agtcctgacgtctaa<br>gaaaccattattatc<br>atgacattaacctat<br>aaaaataggcgtat<br>cacgaggcagaatt<br>tcagataaaaaaa<br>tccttagctttcgcta<br>aggatgatttctggct<br>gcagttgacagcta<br>gctcagtcctaggta<br>ttgtgctagcggatc<br>caaagaggagaaa<br>gaattcatggca | aagaggagaaagatttcATGGCACAGCAGTTGCGCGCGATACGT<br>ACCCGTGGCGCGATCCTGAACGCCGCGGCGGAAGTTTTC<br>GCTCAGCACGGCTATCATGGTGCAAGTATTGCGCAAATTT<br>TAGATAGAGCCCAGGTCACTCGTGGGGCTCTTTACTTTCA<br>TTTTGCTTCCAAAGAGCAGTTGGTGCAAGGTATCTTCGCC<br>GAGCAGGTGACGGAACAGGGTTATCCGCCGCGCACCTAC<br>AAATTGCAGGAATTAGTTGATGAGGCCATGGCACTGGCCT<br>GCCGACTGCCGCAAGAAATTATGCTGCGGGCCGGCGCCA<br>AGCTGGCTATGGAGCCTCAGCTCCAGGAACTGACCGACG<br>GAGGCCCATGGAAGCTTGGGCGCTGCGGTTCGCCTCTA<br>TGGTACTAGCGGCCAAAGAACACGGCGAAGTCTATCCACA<br>TGTGGCGCCCGAAGAATTAGGTCGTTTTCTTGTTTCATCCT<br>GGGTTGGCGCACAAATTGTCTCGGAAGTGACGAGTGGTT<br>GGCAGGACCTGGAAGAACGCATTAGCTCGATCTTCGCATA<br>CACTCTGCCTAGCGTAGCCATGCCGGCAATCTTTGCTCAA<br>CTCGACTTTGGACCGGATCGCGGGACACGTGTGCTGGCA<br>GAAATGCGCGCGGAGGATGCGGCAGCGCTGGTGGCGGA<br>ACCGGCGAGCTAAtaataagcggccgcc | gaggtagtcgacttg<br>acggctagctcagtc<br>ctaggtacagtgcta<br>gcaggaattcacgg<br>attggtaaagtacca<br>atcatctggtttcttttgt<br>aagcttagggtgtctc<br>aaggtgcgtaccttg<br>actgatgagtccgaa<br>aggacgaaacaccc<br>ctctacaaataatttg<br>tttaaagatctaaaga<br>ggagaaatagcatat<br>gtaaagg |

| | | | |
|---|---|---|---|
| SrrA | tcggatgacgtctaa<br>gaaaccattattatc<br>atgacattaacctat<br>aaaaataggcgtat<br>cacgaggcagaatt<br>tcagataaaaaaaa<br>tccttagctttcgcta<br>aggatgatttctggct<br>gcagttgacagcta<br>gctcagtcctaggta<br>ttgtgctagcggatc<br>caaagaggagaaa<br>gaattcatggca | caaagaggagaaagaattcATGGCACAACAGGAACGTGCGATTC<br>GTACCCGCCGCGCGGTTCTAGAAGCGGCAGCAACGGTAT<br>TTGCCGAGCACGGTTATGCTGCCGCTACCGTGGCGGATA<br>TTTTAAAGGTGGCCGGACTGACTAAAGGCGCCCTGTACTT<br>CCACTTTCCCAGTAAAGAAGCGCTTGCACGTGGCATTCTG<br>GAGGCACAGGTGCCGCAACAGCTCGTTCCGCAGCAGCTG<br>AAACTCCAGGAATGGGTAGATGCGGGTATGACTCTGGCTC<br>ATCAGCTGCCGCGCGACCCTGTTCTGCGCGCGGGCGCCC<br>GTTTGTCGGCTGAACATACCGGTAGCGAACAACATGGCAG<br>CGCCTTCCCAACCTGGATCGCCTTTTCTGCCTCATTGCTT<br>GAACAGGCGAAACGGAATGGTGAGGTCCTGGGGCACATA<br>GAACCGGCCGAAACCGCCGAATGCGTCCTCGGCTCCTTC<br>CACGGCATTCAACTTTTATCCCAATTGCAGACGAACTGGG<br>CGGATATCGAGCAGCGTGCCAGCGCGCTGTTTAGGCATG<br>TATTACCGGCAGTGGCGGTGCCAAGTGTGCTGGTGCGCT<br>TAGACACAGCGCCTGATCGAGGAGCCCGCGTCGTTGCAG<br>AACTGGAGGCGATGGCGGCGCAGCCGGATGGGCTGGCA<br>TCGCTGTCAAGCTAAtaataagcggccgccac | agaggtgtcgacttg<br>acggctagctcagtc<br>ctaggtacagtgcta<br>gcaggaatacaaaa<br>cagcttggtcggtttg<br>atattaagcttagggt<br>gtctcaaggtgcgtac<br>cttgactgatgagtcc<br>gaaaggacgaaac<br>acccctctacaaata<br>attttgtttaaagatcta<br>aagaggagaaatag<br>catatgcgtaaa |
| MmfR | ccacctgacgtctaa<br>gaaaccattattatc<br>atgacattaacctat<br>aaaaataggcgtat<br>cacgaggcagaatt<br>tcagataaaaaaaa<br>tccttagctttcgcta<br>aggatgatttctggct<br>gcagtttacggctag<br>ctcagccctaggtat<br>tatgctagcggatcc<br>aaagaggagaaa<br>gaattccgagcg | gaggagaaagaattcATGACGAGCGCCCAACAACCAACTCCTTT<br>CGCGGTCCGGTCCAACGTTCCGCGTGGACCTCATCCGCA<br>GCAAGAGAGGTCGATTAAAACCCGTGCCCAGATTCTGGAA<br>GCGGCGTCGGAAATCTTTGCGAGTCGCGGTTACCGAGGG<br>GCATCCGTCAAGGATGTTGCCGAACGTGTCGGCATGACC<br>AAAGGCGCGGTTTATTTCCACTTTCCCAGCAAAGAATCAC<br>TAGCTATCGCTGTGGTGGAAGAACATTATGCGCGCTGGCC<br>AGCAGCGATGGAAGAAATCCGCATTCAGGGCTTCACACC<br>GCTGGAGACGGTAGAGGAGATGTTACATCGCGCGGCGCA<br>GGCATTCCGCGATGATCCGGTGATGCAGGCGGGTGCCCG<br>TCTGCAGAGTGAACGCGCCTTTATAGACGCTGAGCTGCCC<br>CTGCCGTACGTGGACTGGACCCACCTGCTGGAAGTGCCG<br>TTGCAGGACGCACGTGAAGCCGGGCAGTTGCGTGCGGGT<br>GTTGATCCGGCAGCAGCTGCACGTAGCTTAGTGGCCGCC<br>TTTTTCGGCATGCAACATGTATCTGATAATCTTCACCAAAG<br>AGCGGATATTATGGAGCGCTGGCAGGAACTTCGGGAACT<br>GATGTTTTTTGCTCTCCGCGCCTAAtaataagcggccgccactgata<br>gtgctagtgt | aagggtgtcgacttg<br>acggctagctcagtc<br>ctaggtacagtgcta<br>gcaggaaatatacct<br>gcgggaaggtattat<br>aagcttagggtgtctc<br>aaggtgcgtaccttg<br>actgatgagtccgaa<br>aggacgaaacaccc<br>ctctacaaataattttg<br>tttaaagatctaaaga<br>ggagaaatagcatat<br>gcgtaaa |

| | | | |
|---|---|---|---|
| AvaR1 | ccgactgacgtcta agaaaccattattat catgacattaaccta taaaaataggcgtat cacgaggcagaatt tcagataaaaaaa tccttagctttcgcta aggatgatttctggct gcagcgaggaagc ggggccggccttga cggctagctcagtcc taggtacagtgtagc ttgatcattataacca cacaggaaacagc cagtccgtttaggtct agaaataattttgttt aactttaagaagga ggaattcatgggc | actttaagaaggaggaattcATGGCGCGGCAGGAGCGAGCCATTC GGACGCGGCAGACGATTCTGGTCGCCGCGGCCGAGGTG TTCGACGAGGTGGGATACGAGGCGGCAACCATCTCCGAC GTGCTGAAGCGCTCGGGGGTCACCAAGGGGGCCCTCTAC TTCCACTTCACGTCGAAGCAGGAGCTGGCCCAGGCCGTG CTGGCCGAGCAGGTCGCCTCCCTTCCGCGCGTCCCCGAG CAGGAGCTGAAGCTCCAGCAGTCGCTGGACGAGGCGCTG CTGCTCGCCCATCTGCTCAGGGAAGGCACCGGCGATCCG ATCGTCCAGGGCAGTGTGCGGCTGACCGTGGACCAGGGC TCGCCCAGGGACCATCTCAACCGGCGGGTCCCGATGCAG GCCTGGACCGAGCACACGCAGTCCCTCTTCGAAGAGGCC AGGGCCAAGGGCGAGATCCTGCCCCACGCCGATGTGGAA GCGCTCGCCAAGCTGTTCGTGGGCGCGTTCACCGGCGTG CAGGTCCTCTCGAGGATCATGACCGGGCGCGCGGACCTG GCGGAGCGGGTGGCCGACCTCTACCGCCATCTGATGCCG TCCTTCGCCATGCCGGGGATCCTGGTCCGCCTGGACTTC TCCCCGGAGCGGGGCTCGCGGGTGTACGAAGCCGCCAT GAAGCAGCGGGAGTCGGCGGCAGCGAGTACGACGGACG CGGCACGGACGTTGGAGCATCATCATCATCATCACTGAtG Gtaagcggccgccact | agaggtgtcgacttg acggctagctcagtc ctaggtacagtcattg aaaaaccgttcagct ggttttatcgtcaagct tctagtaaagaggag aaatagcatatgcgtt gc |

**Methods Table 3: Strains and Plasmids**

| | Strain or Plasmid | Relevant Characteristics | Source or Reference |
|---|---|---|---|
| **Strains** | NEB5-alpha | *fhuA2Δ(argF-lacZ)U169 phoA glnV44 Φ80Δ(lacZ)M15 gyrA96 recA1 relA1 endA1 thi-1 hsdR17* | New England Biolabs |
| | DH10B | *F– mcrA Δ(mrr-hsdRMS-mcrBC) φ80lacZΔM15 ΔlacX74 recA1 endA1 araD139 Δ (ara-leu)7697 galU galK λ– rpsL(StrR) nupG* | Novagen |
| | *Streptomyces coelicolor* A3(2) | | ATCC |
| | *Streptomyces coelicolor* M1152 | *ΔactΔredΔcpkΔcda rpoB[C1298T]* | Gomez-Escribano and Bibb, 2011[32] |
| | *Streptomyces coelicolor* MH1 mmfHLP | *S. coelicolor* M512 with pIJ6584::*attB* (MMF producer) | Corre, O'Rourke, Challis 2008[13] |
| | *Streptomyces griseus* NRRL F-5144 | | NRRL |
| | *Streptomyces rochei* ATCC B-1559 | | ATCC |
| | *Streptomyces avermitilis* ATCC 31267 | | ATCC |
| | *Streptomyces virginiae* NRRL B-8091 | | NRRL |
| | *Streptomyces lavendulae* NRRL B-2774 | | NRRL |
| | *Streptomyces tendae* ATCC ISP-5101 | | ATCC |
| | *Streptomyces griseofuscus* ATCC B-5429 | | ATCC |
| **Plasmids** | pLW0003 | | Wilbanks et al. 2023[29] |
| | pET-28b(-) | | Novagen |
| | pRF-ScbR | | Addgene #49370 Stanton et al. 2013[33] |
| | pGEX-6P-1 | | Millipore Sigma |
| | pTU2S-a | | Addgene #74091 Moore et al. 2016[34] |

| | | | |
|---|---|---|---|
| | pRSET-B mCherry | | Addgene #108857 |
| | pTOTAL-ScbR | | This study |
| | pTOTAL-ArpA | | This study |
| | pTOTAL-BarA | | This study |
| | pTOTAL-FarA | | This study |
| | pTOTAL-SabR1 | | This study |
| | pTOTAL-SrrA | | This study |
| | pTOTAL-MmfR | | This study |
| | pTOTAL-AvaR1 | | This study |
| | pREP-ScbR | | This study |
| | pREP-ArpA | | This study |
| | pREP-BarA | | This study |
| | pREP-FarA | | This study |
| | pREP-SabR1 | | This study |
| | pREP-SrrA | | This study |
| | pREP-MmfR | | This study |
| | pREP-AvaR1 | | This study |
| | pGFP-ScbR | | This study |
| | pGFP-ArpA | | This study |
| | pGFP-BarA | | This study |
| | pGFP-FarA | | This study |
| | pGFP-SabR1 | | This study |
| | pGFP-SrrA | | This study |
| | pGFP-MmfR | | This study |
| | pGFP-AvaR1 | | This study |
| | pTOTAL-FarA-mCherry | | This study |
| | pTOTAL-BarA-mCherry | | This study |
| | pTOTAL-MmfR-mCherry | | This study |
| | pTOTAL-SrrA-mCherry | | This study |
| | pAND-SrrA.MmfR | | This study |

**Chemistry Materials and Methods**

**General Remarks**

All reactions were performed in flame-dried glassware under an atmosphere of nitrogen unless otherwise noted. All reagents and solvents, unless otherwise stated, were purchased from Sigma Aldrich, Thermo Fisher, TCI, Oakwood, or Aaron Chemicals and used without further purification. The solvents THF, DCM, toluene, and DMF were dried with an Inert PurSolv solvent purifying system. The reactions were monitored using thin layer chromatography (TLC) plates, Silica gel 60 $F_{254}$ (20 cm x 20 cm), were purchased from Supelco. Purification via column chromatography was performed with silica gel (230-400 mesh) purchased from Silicycle. Proton and Carbon NMR spectra were obtained on a Bruker 300 mHz, Bruker 400 mHz or a Bruker 500 mHz spectrometer in chloroform-*d*. Chemical shifts (δ) are reported in parts per million (ppm) relative to an internal standard (TMS, 0.00 ppm). Multiplicities are reported as singlet (s), doublet (d), triplet (t), quartet (q), and multiplet (m). IR spectra were taken on a Nicolet FTIR spectrometer. Optical rotations were measured from a Rudolph Autopol III S2 polarimeter. Chiral HPLC data was obtained from an Agilent 1220 Infinity LC using a Chiralpak AD-H column with the given conditions.

**Aldol Reaction**

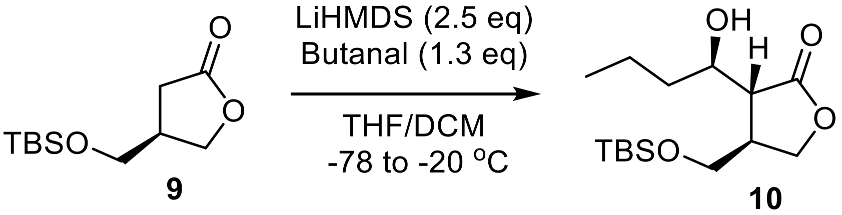

A solution of protected *R*-paraconyl alcohol (**9**, 200mg, 0.87mmol) in anhydrous THF (7.83mL) was added to DCM (0.87mL) at -78°C[35]. To the solution was added lithium(bistrimethylsilyl)amide (1.0M in THF, 2.18mL, 2.18mmol) dropwise. The reaction was allowed to stir for 1.5 hours then butanal (1.13mmol, 0.10mL) was added to the solution dropwise. The reaction continued to stir at -78°C for another hour, after which it was warmed to -20°C where it stirred overnight. The solution was then quenched with a saturated $NH_4Cl$ solution (7.83mL) then extracted with EtOAc (3x15mL). The organic layers were combined to dry with sodium sulfate and concentrated in vacuo. The resulting clear oil was then purified via flash chromatography (5% Petroleum Ether and 5% EtOAc in DCM) to afford TBS protected IM-2 (**10,** 151mg, 56%) as only one diastereomer observed via NMR and polarity. Spectra matched previously reported data[36].

**$^1$H NMR** (500 MHz, $CDCl_3$) δ 4.37 (t, *J* = 8.6 Hz, 1H), 4.04 (t, *J* = 8.4 Hz, 1H), 3.87 (m, 1H), 3.68 (m, 2H), 2.97 (d, J=5.47 Hz, 1H), 2.66 (m, 1H), 2.60 (dd, J=8.7, 5.3 Hz, 1H), 1.34–1.66 (m, 4H), 0.95 (t, *J* = 7.0 Hz, 3H), 0.90 (s, 9H), 0.07 (s, 6H)

**$^{13}$C NMR** (125 MHz, $CDCl_3$) δ 178.2, 70.9, 68.8, 62.7, 47.3, 40.5, 36.7, 25.8, 18.8, 18.2, 13.9, -f5.5

**Deprotection**

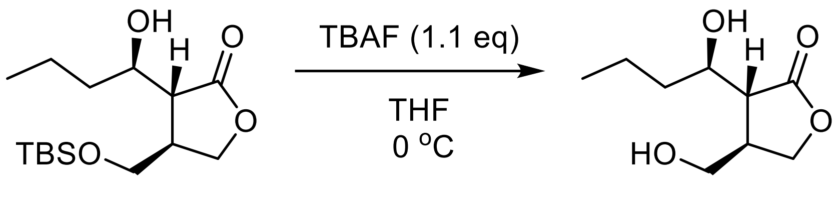

**10** **IM-2 (4)**

TBS protected IM-2 (**10,** 140mg, 0.45mmol ) in dry THF (4.5mL) was cooled to 0°C under nitrogen. TBAF (1.0M in THF, 0.49mL, 0.49mmol) was added dropwise. The reaction stirred for 2 hours at 0°C where TLC showed complete starting material consumption. The reaction was then quenched with $H_2O$ (4.5mL) and extracted three times with EtOAc (10mL each). The organics were combined and dried with sodium sulfate, filtered, and concentrated. The resulting oil was then purified via flash chromatography to afford purified IM-2 (**4,** 71mg, 80%). Spectra matched previously published data[36].

**$^1$H NMR** (500 MHz, $CDCl_3$) δ 4.42 (t, *J* = 8.7 Hz, 1H), 4.03 (m, 1H), 3.99 (t, J=8.8 Hz, 1H), 3.76 (dd, *J* = 10.7, 5.3 Hz, 1H), 3.64 (dd, *J* = 10.7, 6.5 Hz, 1H), 2.70–3.00 (br, 2H), 2.72–2.84 (m, 1H), 2.64 (dd, *J* = 4.8, 9.3 Hz, 1H), 1.35–1.67 (m, 4H), 0.96 (t, *J* = 7.1 Hz, 3H)

**$^{13}$C NMR** (125 MHz, $CDCl_3$) δ 177.3, 70.7, 68.4, 63.0, 49.0, 40.2, 36.1, 19.1,13.9

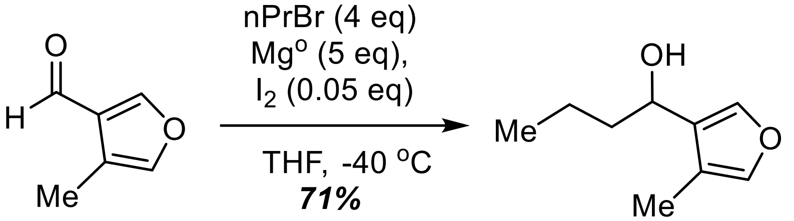

**11** **12**

**Alkylation of Furaldehyde**

Magnesium turnings (17.8 mmol, 4 eq) were crushed with a mortar and pestle, then transferred to a 2-neck conical flask equipped with a stir bar and fitted with a reflux condenser. The vessel was then flame-dried under vacuum for 5 minutes. Iodine (0.223 mmol, 0.05 eq) was added and placed under an atmosphere of nitrogen. THF (10 mL) was then added and brough to 65 °C. 1-bromopropane (13.4 mmol, 4 eq) was then added dropwise over 10 minutes. The reaction was allowed to stir at 65 °C for 2 hours before it was brough to room temperature. In a separate flame-dried flask, a solution of aldehyde (**11,** 4.46 mmol, 1 eq) in THF (10 mL) was brought to -40 °C in an acetonitrile-dry ice bath[37]. The freshly prepared Grignard reagent was then added dropwise at -40 °C and the reaction was allowed to slowly warm to room temperature. Upon consumption of the aldehyde starting material, the reaction was diluted with $Et_2O$ (20 mL) and quenched with saturated $NH_4Cl$ (20 mL). The aqueous layer was extracted with $Et_2O$ (3 x 20 mL). Combined organic layers were dried with magnesium sulfate, filtered, and concentrated. The crude oil was then purified on silica gel in 17:3 Hexanes:EtOAc to afford a clear oil, *71%*.

**TLC:** $R_f$ = 0.36 (17:3 Hexanes: EtOAc, CAN).

**$^1$H NMR (500 MHz, CDCl$_3$):** δ 7.30 (s, 1H), 7.16 (s, 1H), 4.63 (t, *J* = 5.8 Hz, 1H), 2.04 (d, *J* = 1.3 Hz, 3H), 1.81 – 1.70 (m, 2H), 1.54 – 1.44 (m, 1H), 1.44 – 1.33 (m, 1H), 0.96 (t, *J* = 7.3 Hz, 3H).

**$^{13}$C NMR (125 MHz, CDCl$_3$):** δ 140.31, 139.54, 128.92, 119.03, 66.61, 39.22, 19.15, 13.94, 8.55.

**FTIR (Neat, cm$^{-1}$):** *v* 3441.29 cm$^{-1}$, 1545.07 cm$^{-1}$, 1454.92 cm$^{-1}$.

**HRMS (+ESI):** calcd for $C_9H_{13}O$ [M-H$_2$O]$^-$ 137.0972, found 137.0956 (-11.7 ppm)

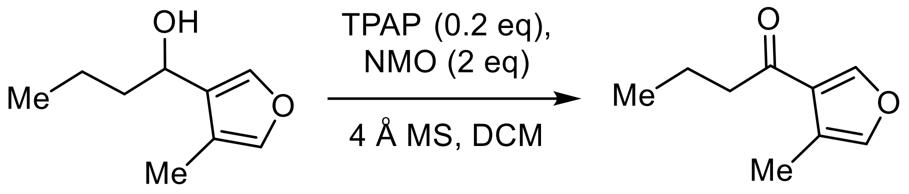

**12** **13**

**Ley Oxidation**

A flame-dried round bottom flask equipped with a stir bar was charged with TPAP (0.389 mmol, 0.2 eq) and NMO (3.891 mmol, 2 eq). The solids were stirred under vacuum for 30 min. Crushed 4 Å MS were then added and brought to room temperature under vacuum. The flask was then put under nitrogen and a solution of alcohol (1.945 mmol, 1 eq) in DCM (0.25 M) was added. Reaction was allowed to stir at room temperature. Following consumption of the alcohol starting material by TLC, the solution was filtered over a pre-equilibrated plug of silica gel. The silica plug was then washed with 200 mL of a solution of 5% EtOAC in Hexanes. Filtrate was concentrated to afford a clear oil, (*77%*).

**TLC:** $R_f$ = 0.61 (9:1 Hexanes:EtOAc, UV).

**$^1$H NMR (500 MHz, CDCl$_3$):** δ 7.95 (s, 1H), 7.20 (s, 1H), 2.69 (d, *J* = 7.3 Hz, 2H), 2.21 (s, 3H), 1.73 (d, *J* = 7.3 Hz, 2H), 0.98 (t, *J* = 7.4 Hz, 3H).

**$^{13}$C NMR (125 MHz, CDCl$_3$):** δ 196.47, 148.42, 141.41, 126.63, 120.47, 42.63, 17.83, 13.86, 9.51.

**FTIR (Neat, cm$^{-1}$):** *v* 1674.88 cm$^{-1}$, 1528.15 cm$^{-1}$.

**HRMS (+ESI):** calcd for $C_9H_{13}O_2$ [M+H]$^+$ 153.0916, found 153.0907 (-5.8 ppm)

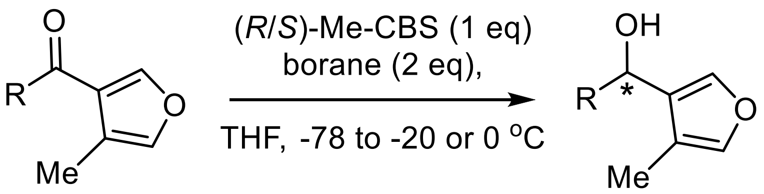

**13** **(*R*)-/(*S*)-12**

**General Procedure 1: CBS Reduction of 1-(4-methylfuran-3-yl)butan-1-one**

In a flame-dried round bottom flask equipped with a stirbar was dissolved (*R*)- or (*S*)-Me-CBS (0.657 mmol, 1 eq) in THF (5 mL) and brough to -78 °C in an isopropanol-dry ice bath. A solution of ketone (0.657 mmol, 1 eq) in THF (1 mL) was added to the vessel and stirred at -78 °C for 5 minutes. After which, a solution of borane, 1M in THF, (1.314 mmol, 2 eq) was added. The reaction was then warmed to the specified temperature. After consumption of the ketone by TLC, the reaction was quenched at 0 °C with MeOH (3 mL). A saturated solution of $NH_4Cl$ (10 mL) was then added and partitioned with $Et_2O$ (10 mL). The phases were mixed and separated, and the aqueous phase was further extracted with $Et_2O$ (2 x 10 mL). Combined organic layers were dried with $MgSO_4$, gravity filtered, and concentrated. The resultant crude oil was purified on silica gel.

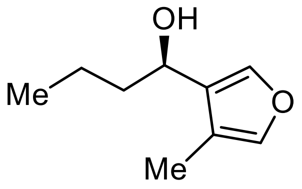

**(*R*)-12**

**(*R*)-1-(4-methylfuran-3-yl)butan-1-ol** prepared with 1-(4-methylfuran-3-yl)butan-1-one (**13**), (*S*)-Me-CBS and catecholborane, warmed to -20 °C and purified in 17:3 Hexanes:EtOAc to afford a clear oil, *75%.* The enantiomeric excess was determined by HPLC using a Chiralpak AD-H column in 5% isopropanol in hexanes at 1mL/min at 30 °C with detection at 217 nm ($t_{major}$ =6.55 min, $t_{minor}$ = 7.11 min, 90% ee).

**TLC:** $R_f$ = 0.36 (17:3 Hexanes: EtOAc, CAN).

**$^1$H NMR (500 MHz, $CDCl_3$):** δ 7.30 (s, 1H), 7.16 (s, 1H), 4.62 (t, *J* = 6.5 Hz, 1H), 2.05 (s, 3H), 1.82 – 1.70 (m, 2H), 1.52 – 1.42 (m, 1H), 1.43 – 1.30 (m, 1H), 0.96 (t, *J* = 7.1 Hz, 3H).

**$^{13}$C NMR (125 MHz, $CDCl_3$):** δ 140.31, 139.54, 128.93, 119.02, 66.62, 39.22, 19.14, 13.93, 8.55.

**FTIR (Neat, cm$^{-1}$):** *v* 3353.61 cm$^{-1}$, 1541.15 cm$^{-1}$, 1457.35 cm$^{-1}$.

**HRMS (+ESI):** calcd for $C_9H_{13}O$ [M-$H_2O$]$^-$ 137.0972, found 137.0955 (-12.4 ppm)

$[\alpha]_D^{21}$ **:** +15.55 ° (*c* = 4.5 mg/mL, $CHCl_3$) for 90% *ee*.

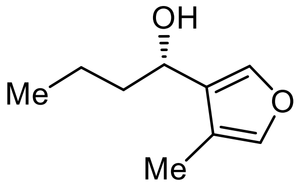

**(*S*)-12**

**(*S*)-1-(4-methylfuran-3-yl)butan-1-ol** prepared with 1-(4-methylfuran-3-yl)butan-1-one (**13**), (*R*)-Me-CBS and catecholborane, warmed to -20 °C and purified in 17:3 Hexanes:EtOAc to afford a clear oil, *73%.* The enantiomeric excess was determined by HPLC using a Chiralpak AD-H column in 5% isopropanol in hexanes at 1mL/min at 30 °C with detection at 217 nm ($t_{major}$ =7.12 min, $t_{minor}$ = 6.55 min, 86% ee).

**TLC:** $R_f$ = 0.36 (85:15 Hexanes: EtOAc, CAN).

**$^1$H NMR (500 MHz, $CDCl_3$)** δ 7.30 (s, 1H), 7.16 (s, 1H), 4.62 (d, *J* = 6.9 Hz, 1H), 2.05 (s, 3H), 1.88 – 1.64 (m, 2H), 1.61 – 1.55 (br s, 1H), 1.54 – 1.42 (m, 1H), 1.44 – 1.24 (m, 1H), 0.96 (t, *J* = 7.5 Hz, 3H).

**$^{13}$C NMR (126 MHz, $CDCl_3$)** δ 140.31, 139.54, 128.91, 119.03, 66.62, 39.21, 19.15, 13.94, 8.56.

**FTIR (Neat, cm$^{-1}$):** $v$ 3389.70 cm$^{-1}$, 1545.93 cm$^{-1}$, 1458.07 cm$^{-1}$.

**HRMS (+ESI):** calcd for $C_9H_{13}O$ $[M-H_2O]^-$ 137.0972, found 137.0956 (-11.7 ppm)

$[\alpha]^{21}_{D}$ : -17.14 $^{O}$ ($c$ 0.07, $CHCl_3$) for 86% *ee*.

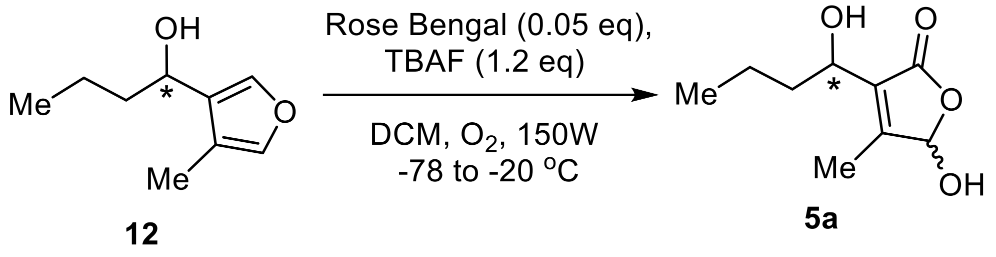

**General Procedure 2: Photo-oxidation**

A solution of furyl alcohol (0.162 mmol, 1 eq) in DCM (1.5 mL) was brough to -78 °C in an isopropanol-dry ice bath. A solution of TBAF, 1 M in THF (0.243 mmol, 1.5 eq) was added followed by Rose Bengal (0.003 mmol, 0.02 eq). The reaction turned vibrant pink and was shielded from light. The solution was sparged with a nitrogen balloon for 10 minutes, then sparged with an oxygen balloon for an additional 10 minutes. After which, a 150W white light bulb was introduced and the reactions were stirred in the light and the reactions were allowed to slowly warm to -20 °C. After 2 hours, or consumption of the furyl alcohol by TLC, the reaction was filtered over a short pad of silica. The silica cake was then washed with EtOAc (10 mL, 2x) and the filtrate was concentrated under reduced pressure. The resulting pale-pink crude oil was loaded onto a pre-equilibrated silica gel column and eluted in 1:1 Hexanes:EtOAc to afford a clear oil.

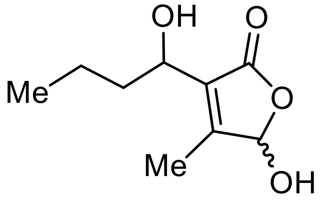

**5a**

**5-hydroxy-3-(1-hydroxybutyl)-4-methylfuran-2(5H)-one** (rac-SAB1), 78% as a 7:1 mix of regioisomers.

**TLC:** $R_f$ = 0.25 (1:1 Hexanes: EtOAc, CAN).

**$^1$H NMR (400 MHz, CDCl$_3$)** δ 5.84 (s, 1H), 5.28 (s, 1H), 4.48 (s, 1H), 3.28 (br s, 0.5H), 3.11 (br s, 0.5H) 2.06 (d, $J$ = 5.7 Hz, 3H), 1.86 – 1.73 (m, 1H), 1.70 – 1.55 (m, 1H), 1.53 – 1.23 (m, 3H, exp. 2H), 0.94 (dt, $J$ = 6.8, 1.5 Hz, 3H).

**$^{13}$C NMR (126 MHz, CDCl$_3$)** δ 171.68, 157.69, 130.00, 98.91, 66.46, 37.96, 18.74, 13.76, 11.40.

**FTIR (Neat, cm$^{-1}$):** $v$ 1744.77 cm$^{-1}$, 3351.32 cm$^{-1}$.

**HRMS (+ESI):** calcd for $C_9H_{14}O_4Na$ $[M+Na]^+$ 209.0790, found 209.0784 (-2.9 ppm)

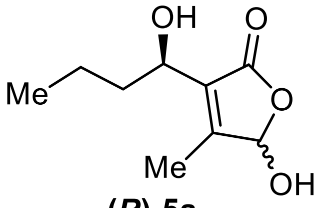

**(*R*)-5a**

**5-hydroxy-3-((*R*)-1-hydroxybutyl)-4-methylfuran-2(5H)-one (**(*R*)-SAB1), 61% as a 6:1 mix of regioisomers

**TLC:** $R_f$ = 0.30 (1:1 Hexanes: EtOAc, CAN).

**$^1$H NMR (500 MHz, CDCl$_3$)** δ 5.86 (q, *J* = 4.7 Hz, 1H), 4.57 – 4.43 (m, 1H), 4.34 (d, *J* = 9.0 Hz, 0.5H), 4.10 (d, *J* = 8.3 Hz, 0.5H), 2.94 (br s, 0.5H), 2.83 (br s, 0.5H) 2.06 (d, *J* = 4.7 Hz, 3H), 1.87 – 1.74 (m, 1H), 1.74 – 1.54 (m, 3H), 1.44 (s, 1H), 1.32 (d, *J* = 3.1 Hz, 1H), 0.95 (dt, *J* = 7.5, 2.2 Hz, 3H).

**$^{13}$C NMR (126 MHz, CDCl$_3$)** δ 156.81, 130.80, 130.43, 98.58, 98.36, 66.66, 66.62, 38.65, 38.34, 18.76, 13.80, 13.77, 11.52, 11.39.

**FTIR (Neat, cm$^{-1}$):** *v* 1742.68 cm$^{-1}$, 3361.40 cm$^{-1}$.

**HRMS (+ESI):** calcd for $C_9H_{14}O_4Na$ $[M+Na]^+$ 209.0790, found 209.0784 (-2.9 ppm)

$[\alpha]_D^{21}$ **:** +13.54$^O$ (*c* 0.93, CHCl$_3$)

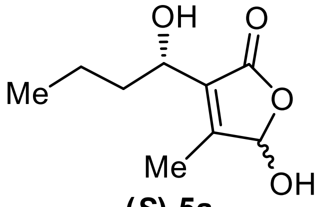

**(*S*)-5a**

**5-hydroxy-3-((*S*)-1-hydroxybutyl)-4-methylfuran-2(5H)-one** ((*S*)-SAB1), 52% as a 9:1 mix of regioisomers

**TLC:** $R_f$ = 0.30 (1:1 Hexanes: EtOAc, CAN).

**$^1$H NMR (500 MHz, CDCl$_3$)** δ 5.85 (s, 1H), 4.48 (t, *J* = 7.1 Hz, 1H), 2.06 (s, 3H), 1.86 – 1.73 (m, 1H), 1.70 – 1.56 (m, 1H), 1.48 – 1.36 (m, 1H), 1.36 – 1.23 (m, 1H), 0.94 (t, *J* = 7.1 Hz, 3H).

**$^{13}$C NMR (126 MHz, CDCl$_3$)** δ 171.80, 157.67, 130.29, 98.85, 66.49, 38.20, 18.76, 13.77, 11.50.

**FTIR (Neat, cm$^{-1}$):** *v* 1742.77 cm$^{-1}$, 3359.40 cm$^{-1}$.

**HRMS (+ESI):** calcd for $C_9H_{14}O_4Na$ $[M+Na]^+$ 209.0790, found 209.0785 (-2.4 ppm)

$[\alpha]_D^{21}$ **:** -16.66$^O$ (*c* = 0.15, CHCl$_3$)

### NMR Spectra

$^{1}H$ for compound **10**

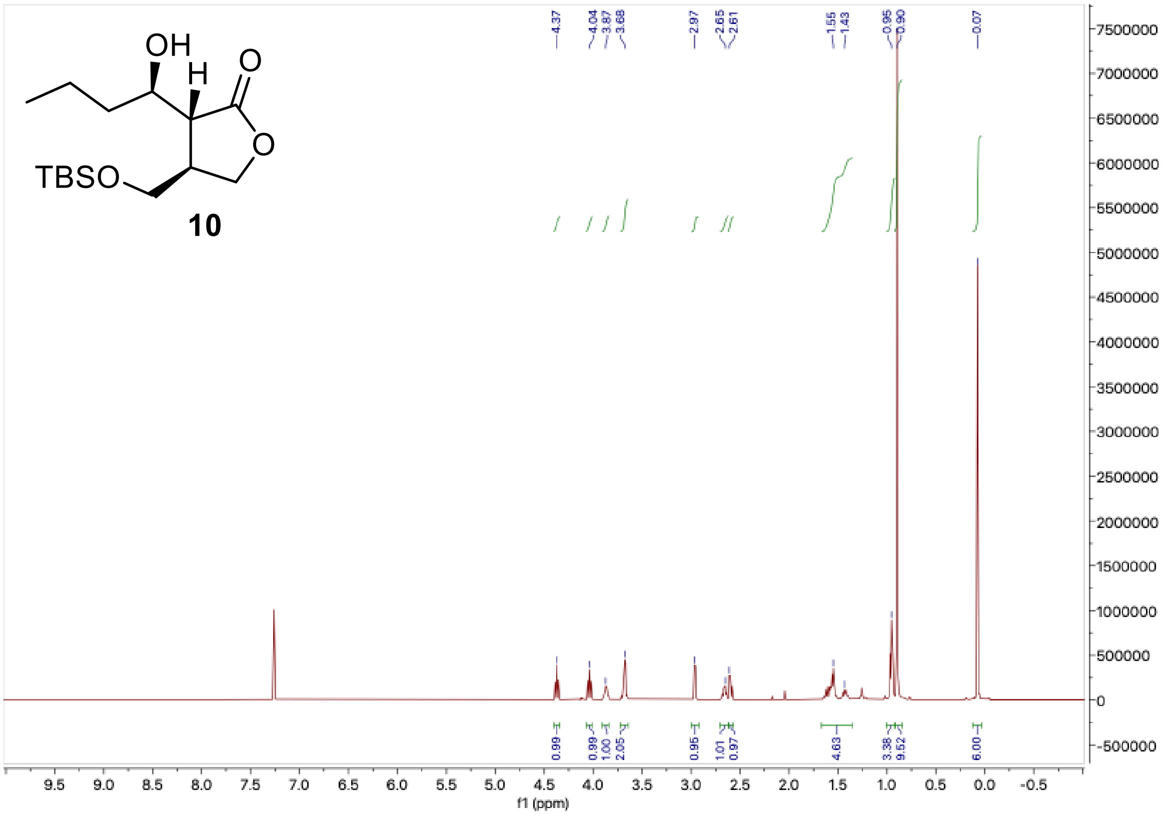

$^{13}C$ for compound **10**

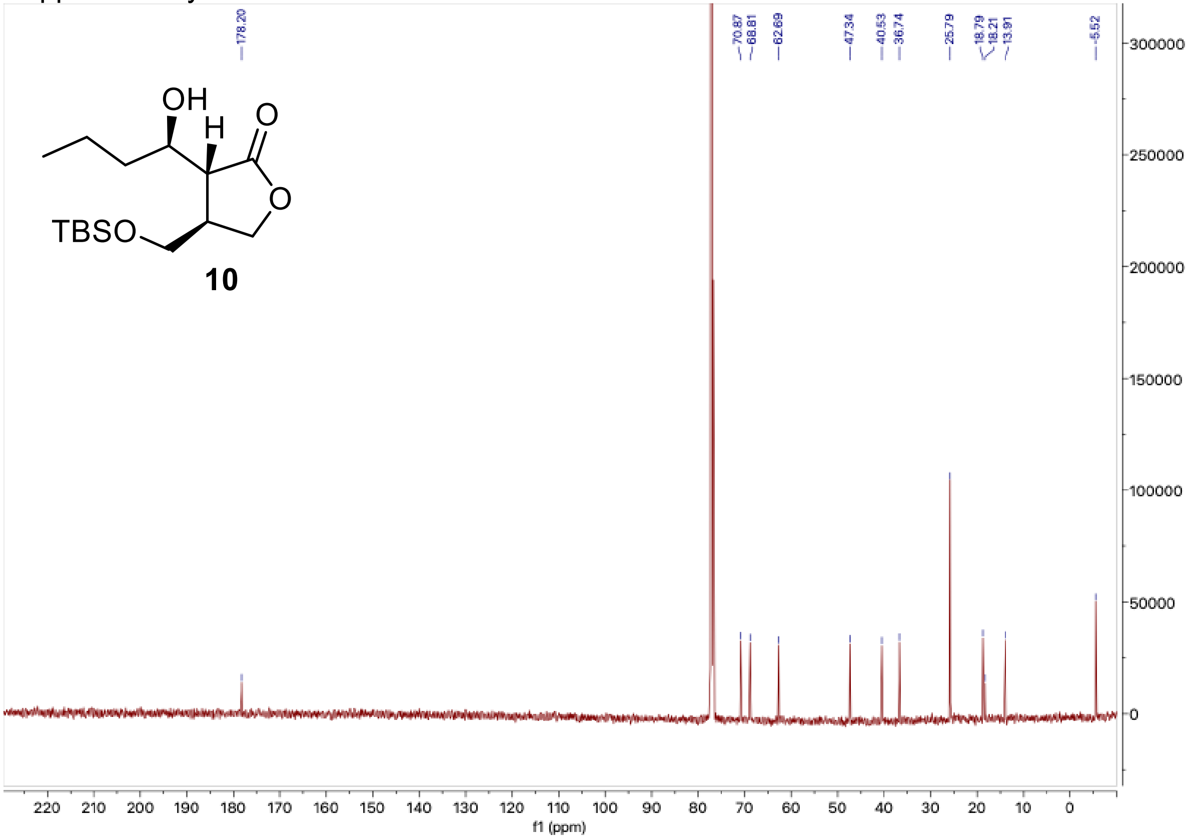

[1]H for IM-2 (**4**)

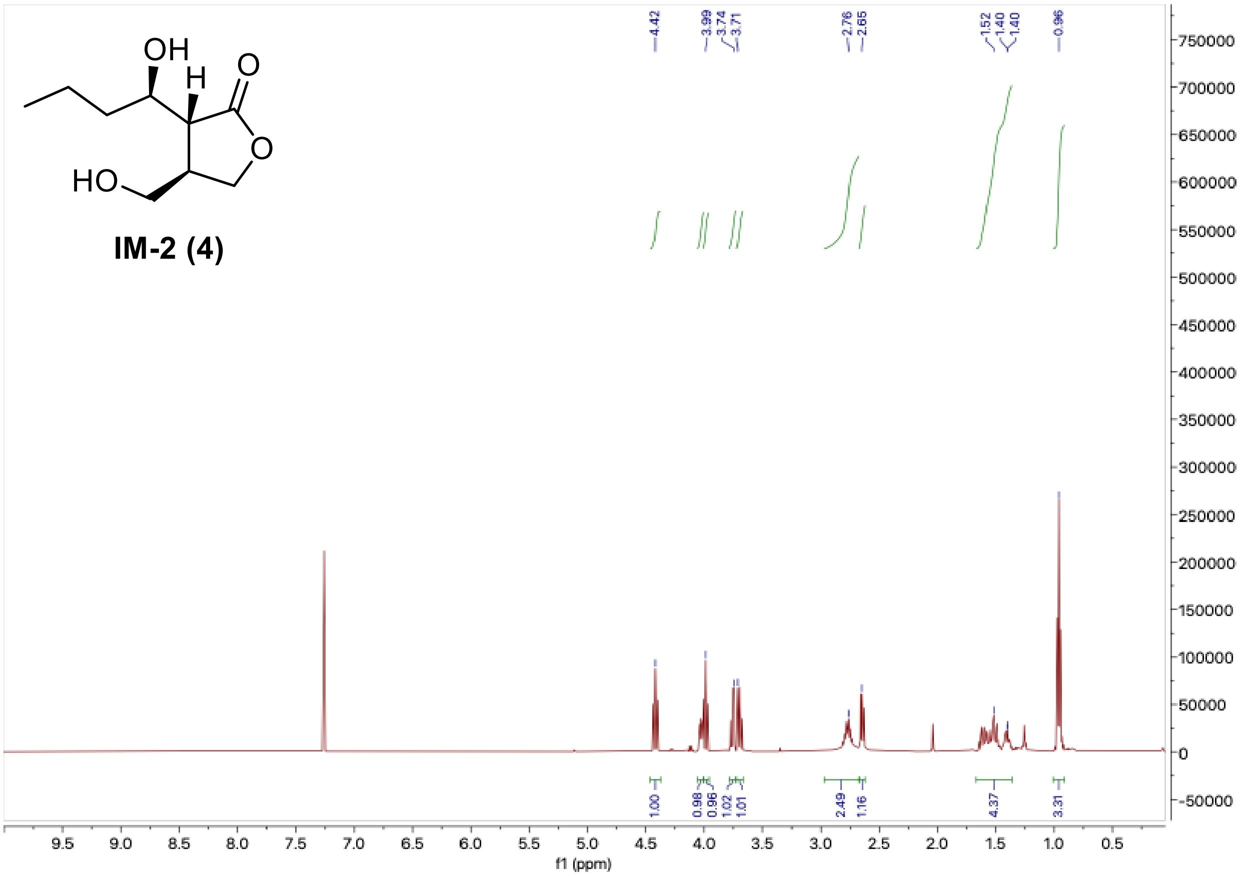

[13]C for IM-2 (**4**)

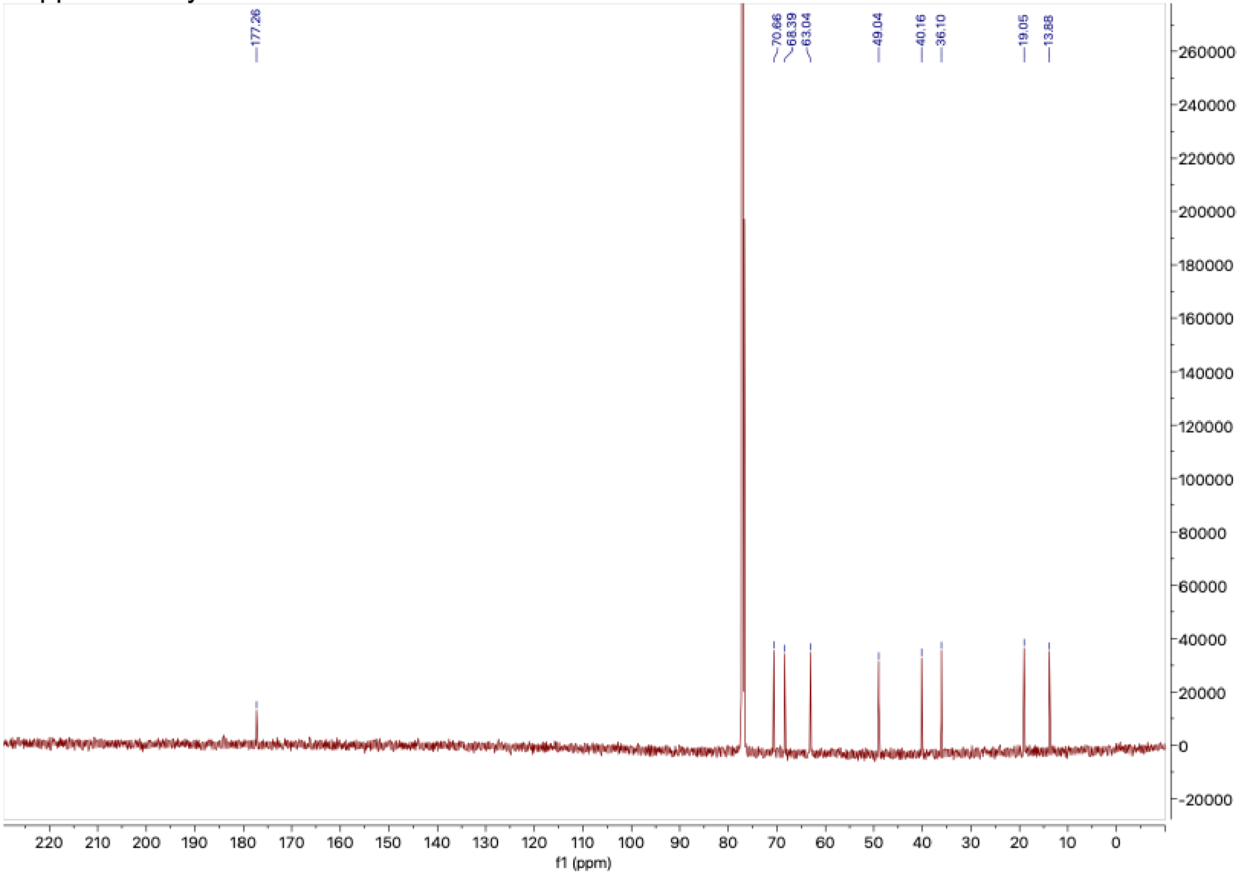

[1]H for Compound **12**

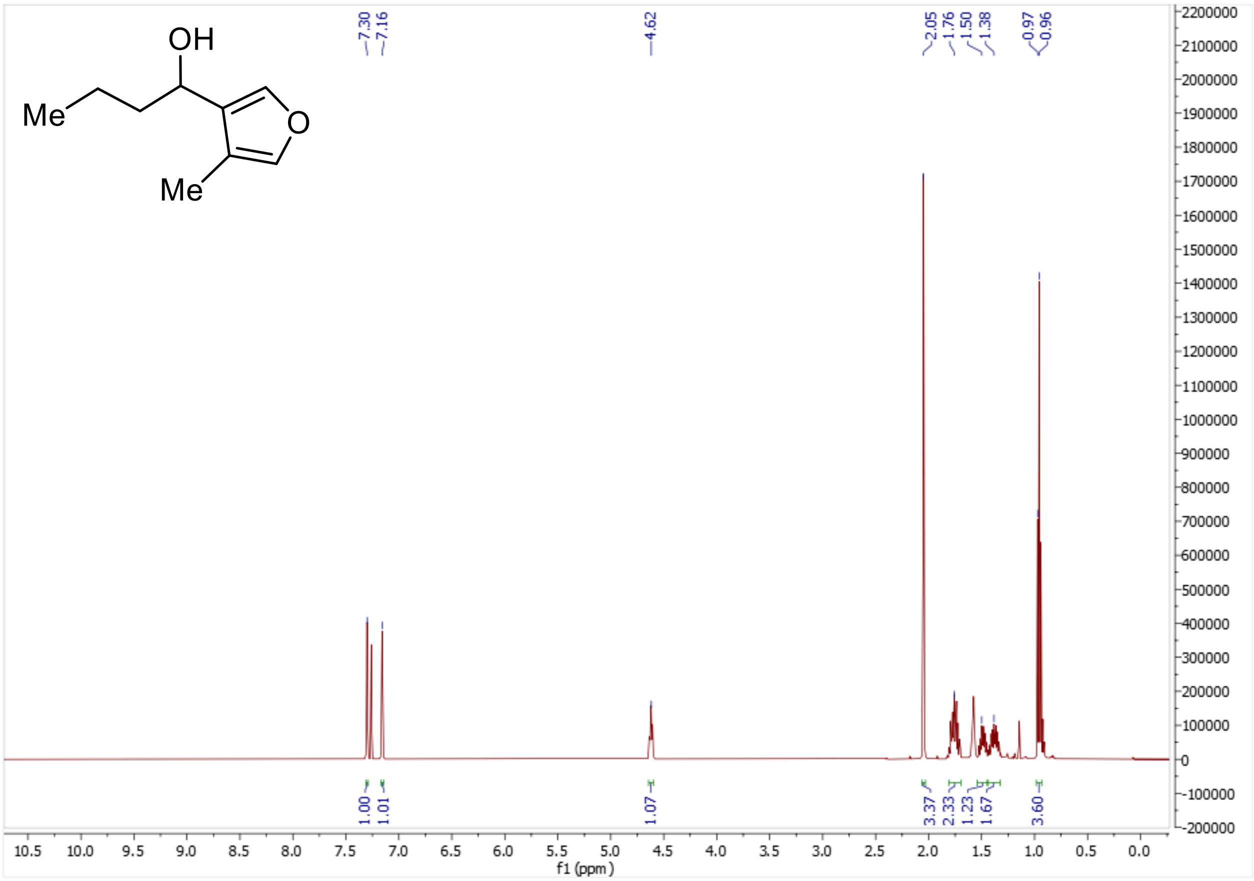

[13]C for Compound **12**

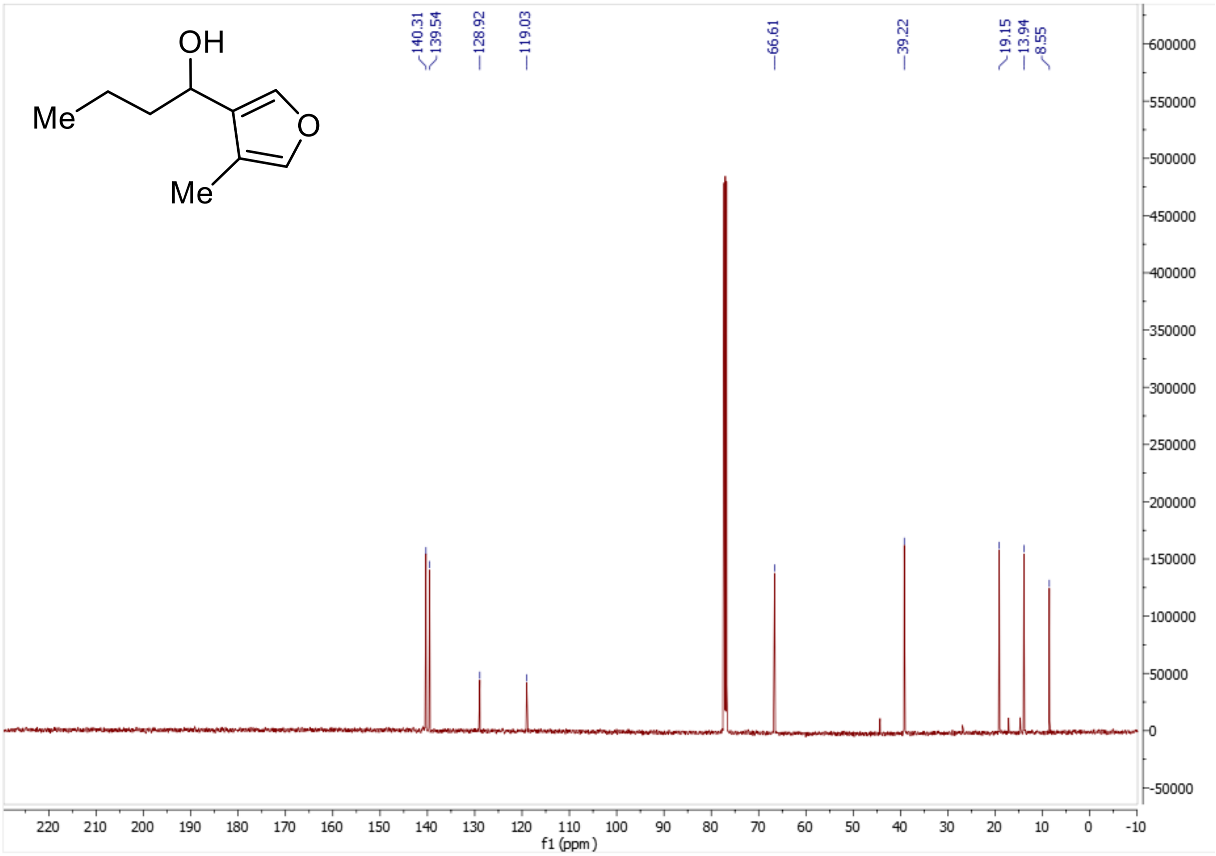

$^{1}$H for Compound **13**

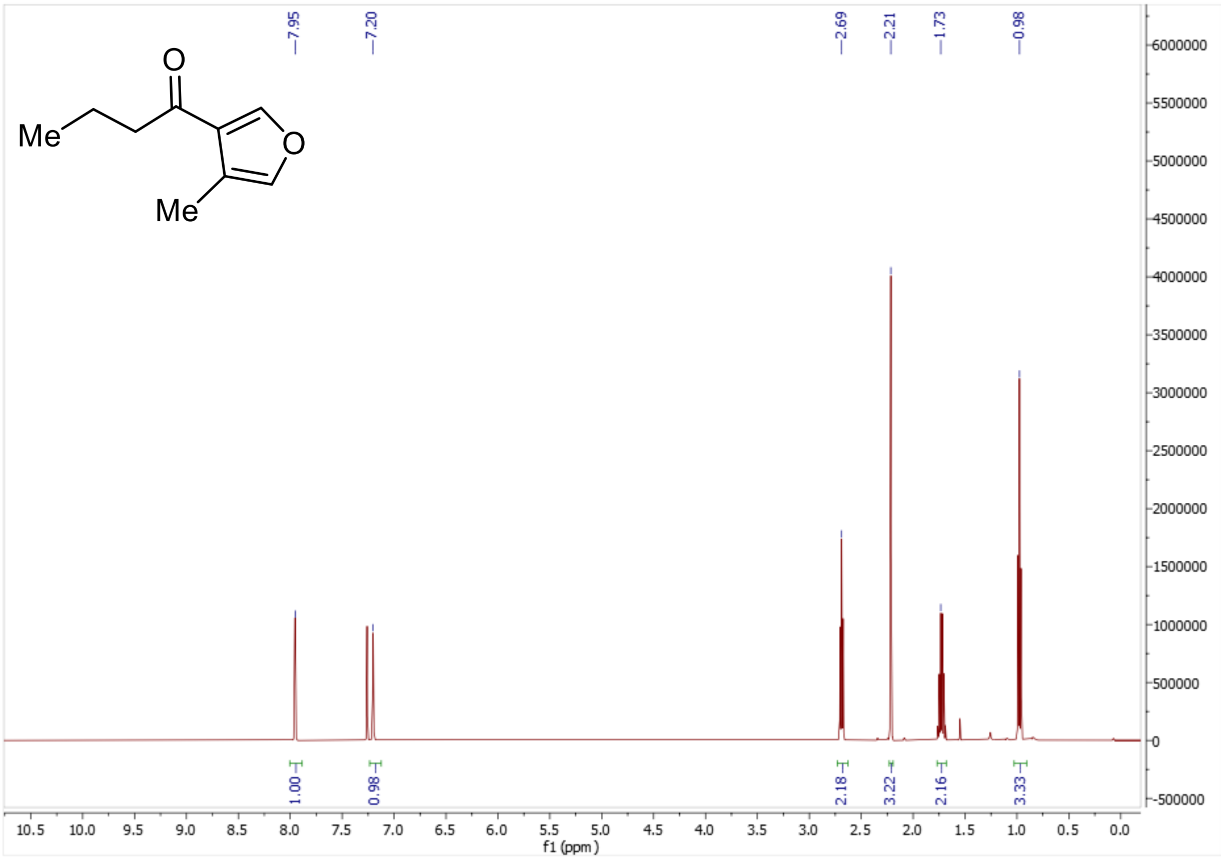

$^{13}$C for Compound **13**

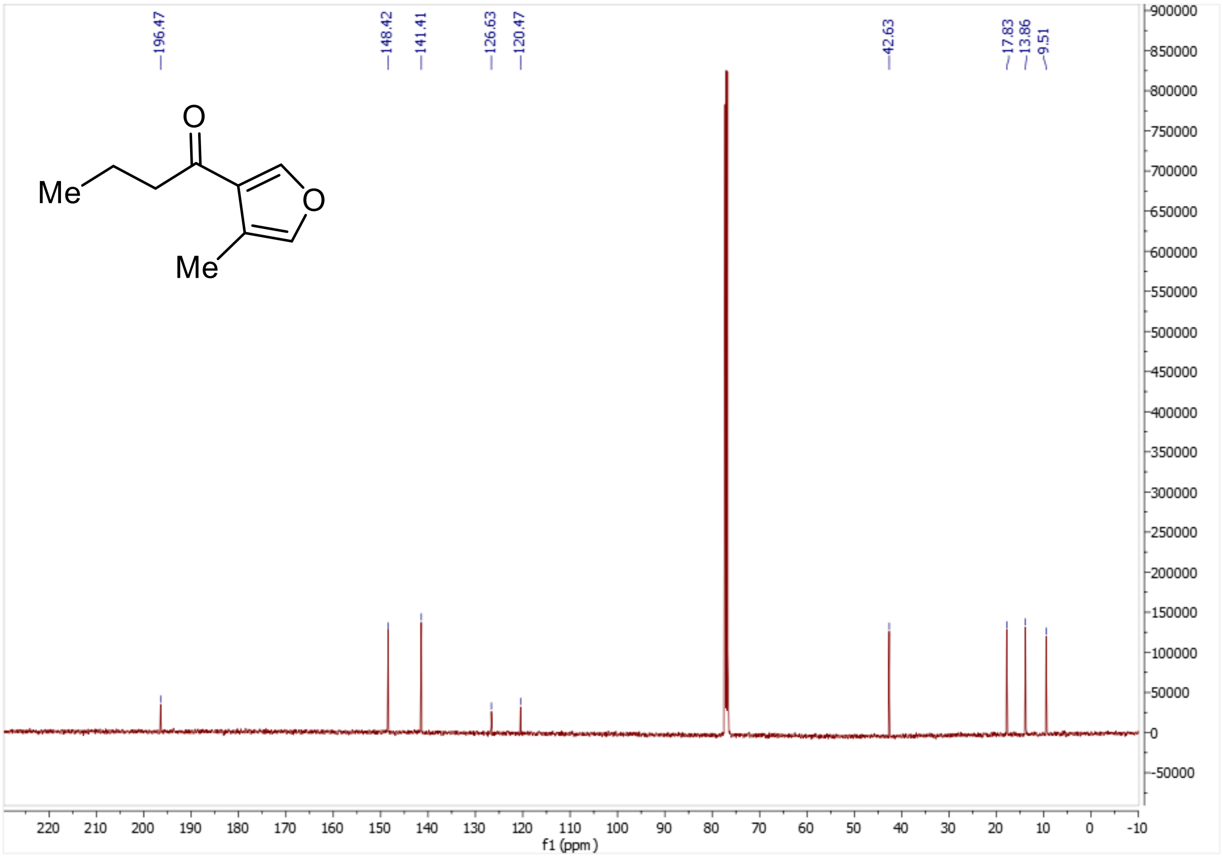

$^{1}$H for Compound ***R*-12**

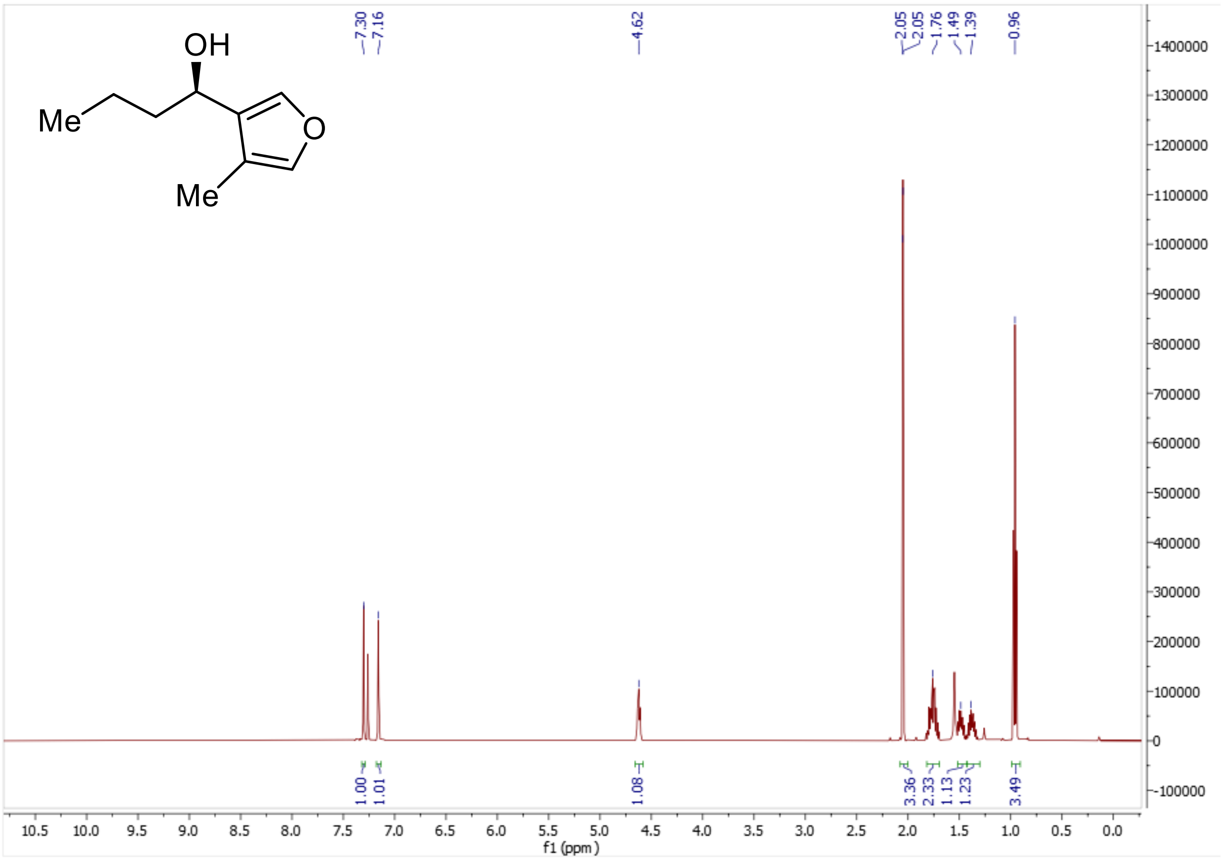

$^{13}$C for Compound ***R*-12**

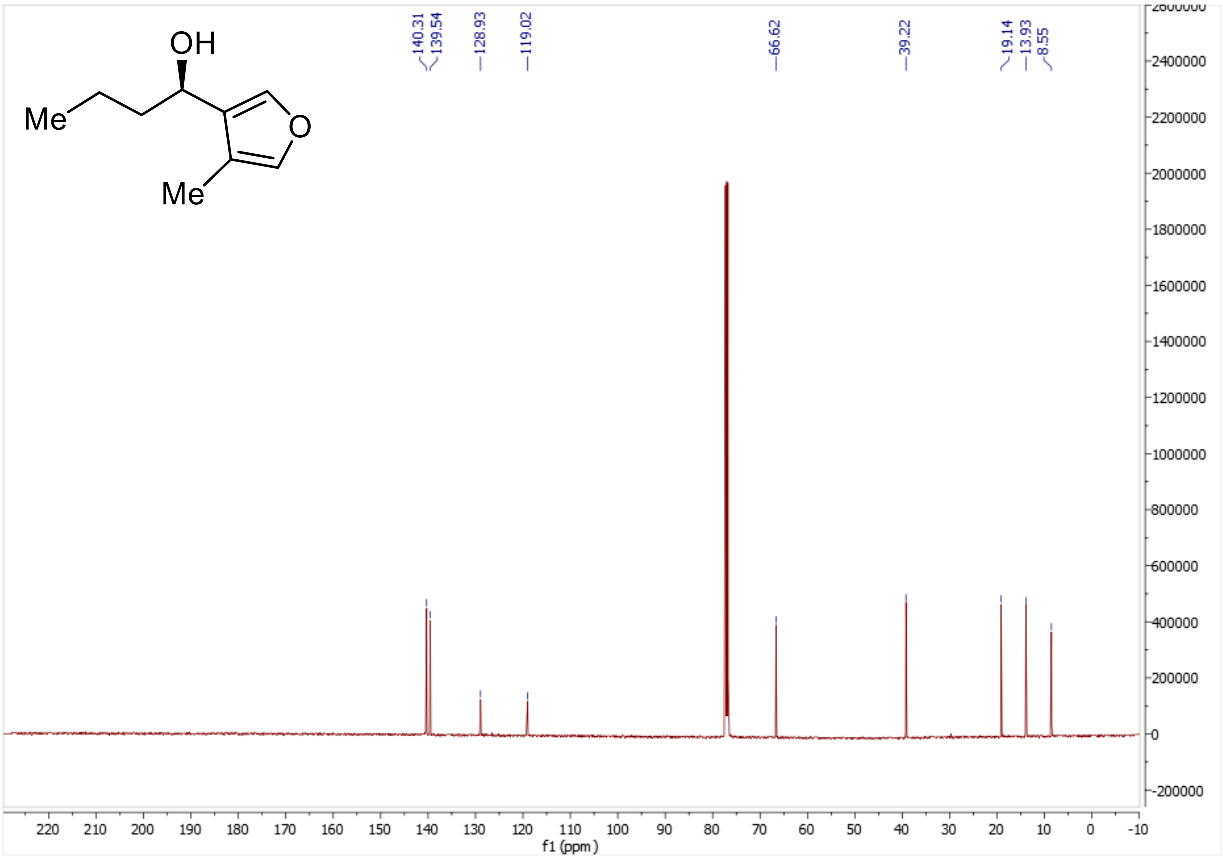

$^{1}$H for Compound ***S*-12**

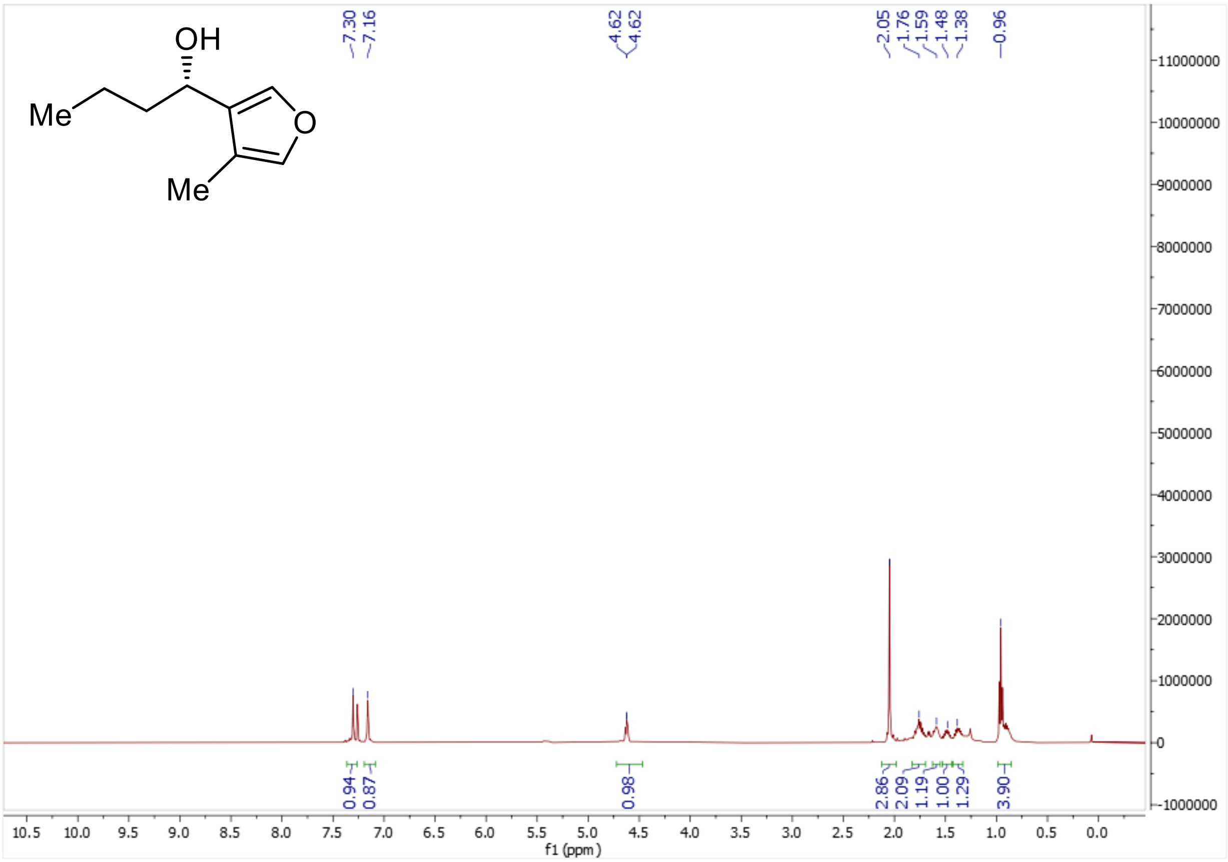

$^{13}$C for Compound ***S*-12**

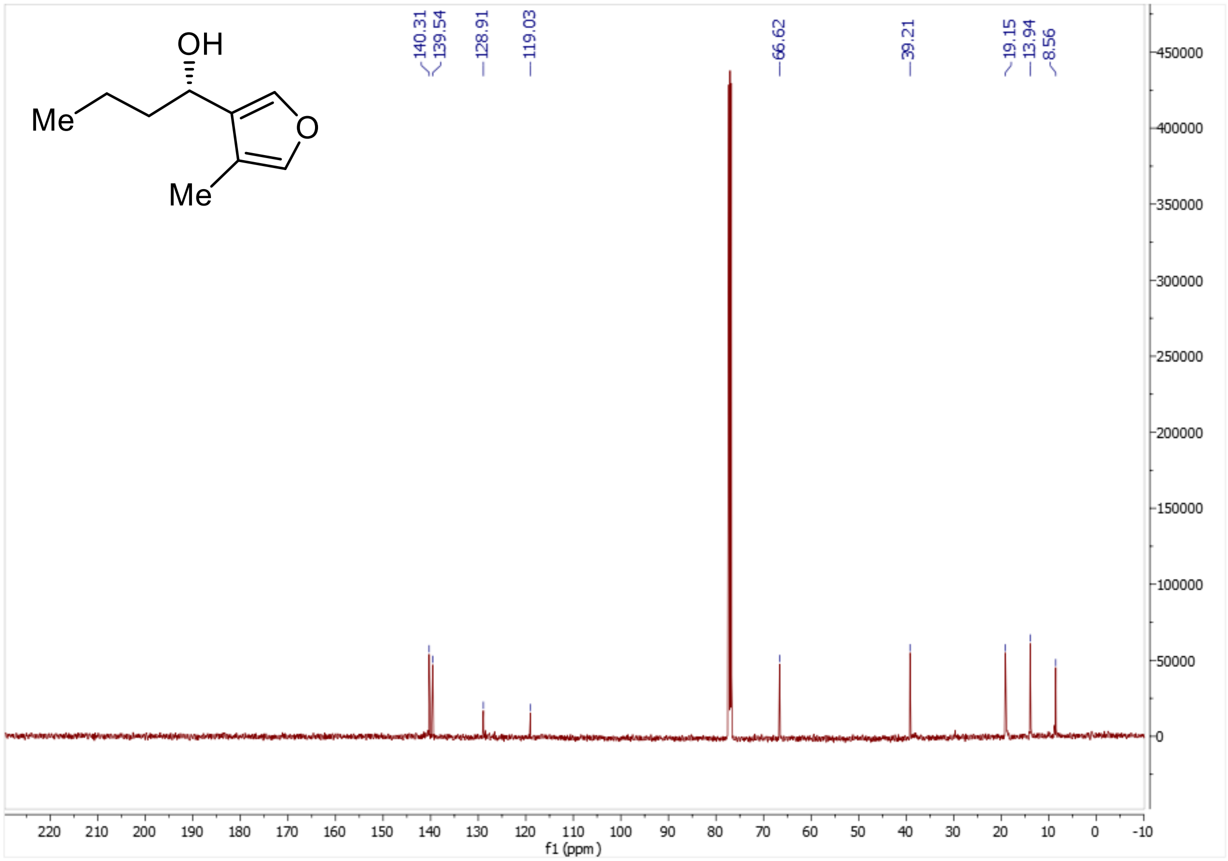

$^{1}$H for racemic **SAB1 (5a)**

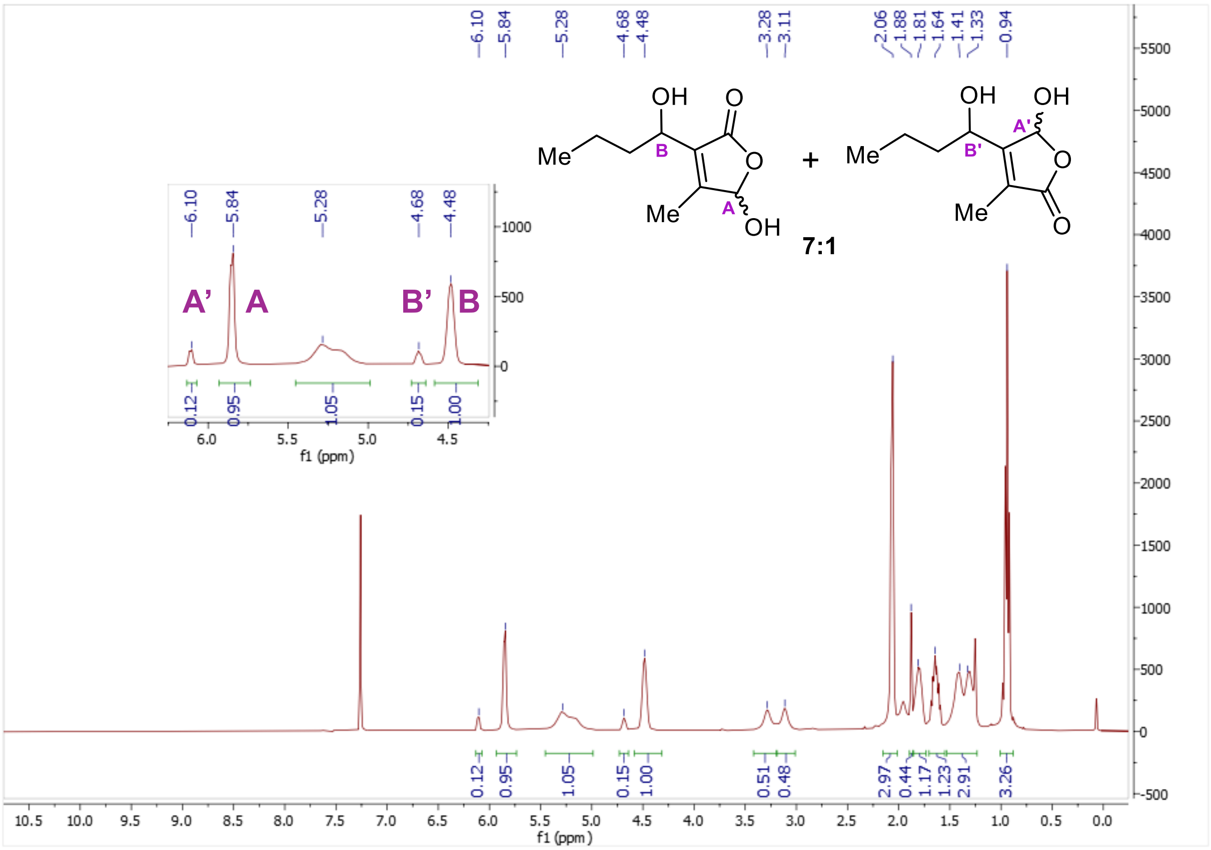

$^{13}$C for racemic **SAB1 (5a)**

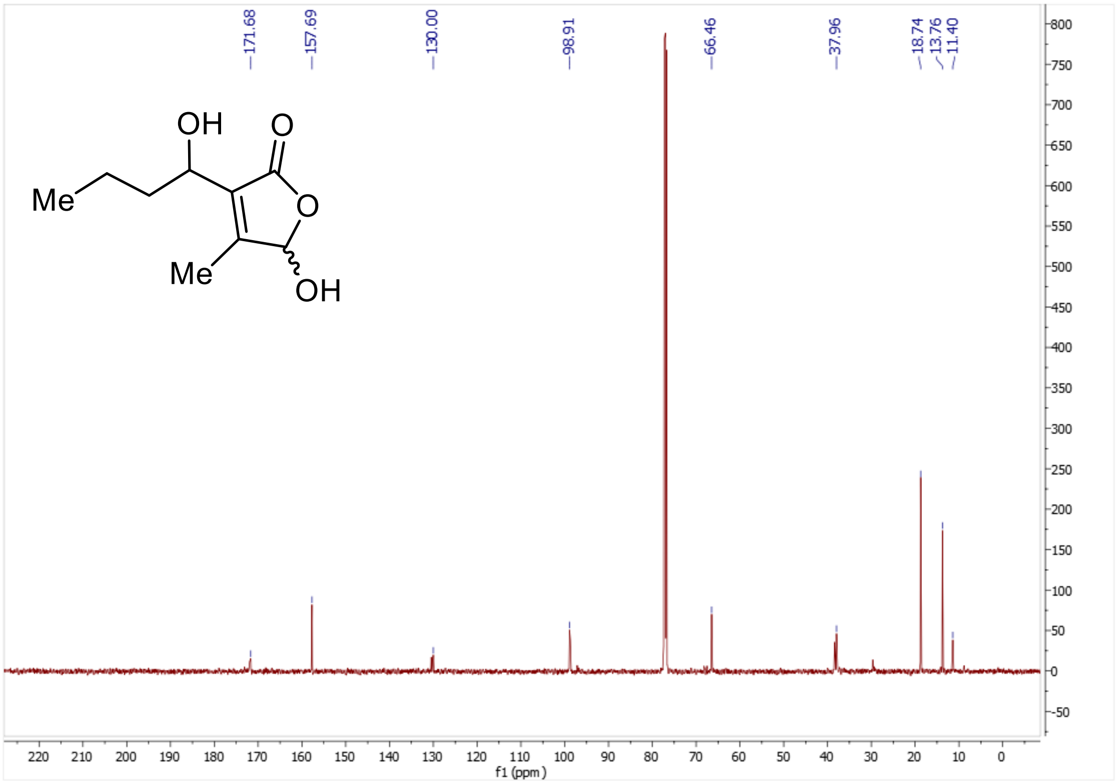

$^{1}$H for ***R*-SAB1 (*R*-5a)**

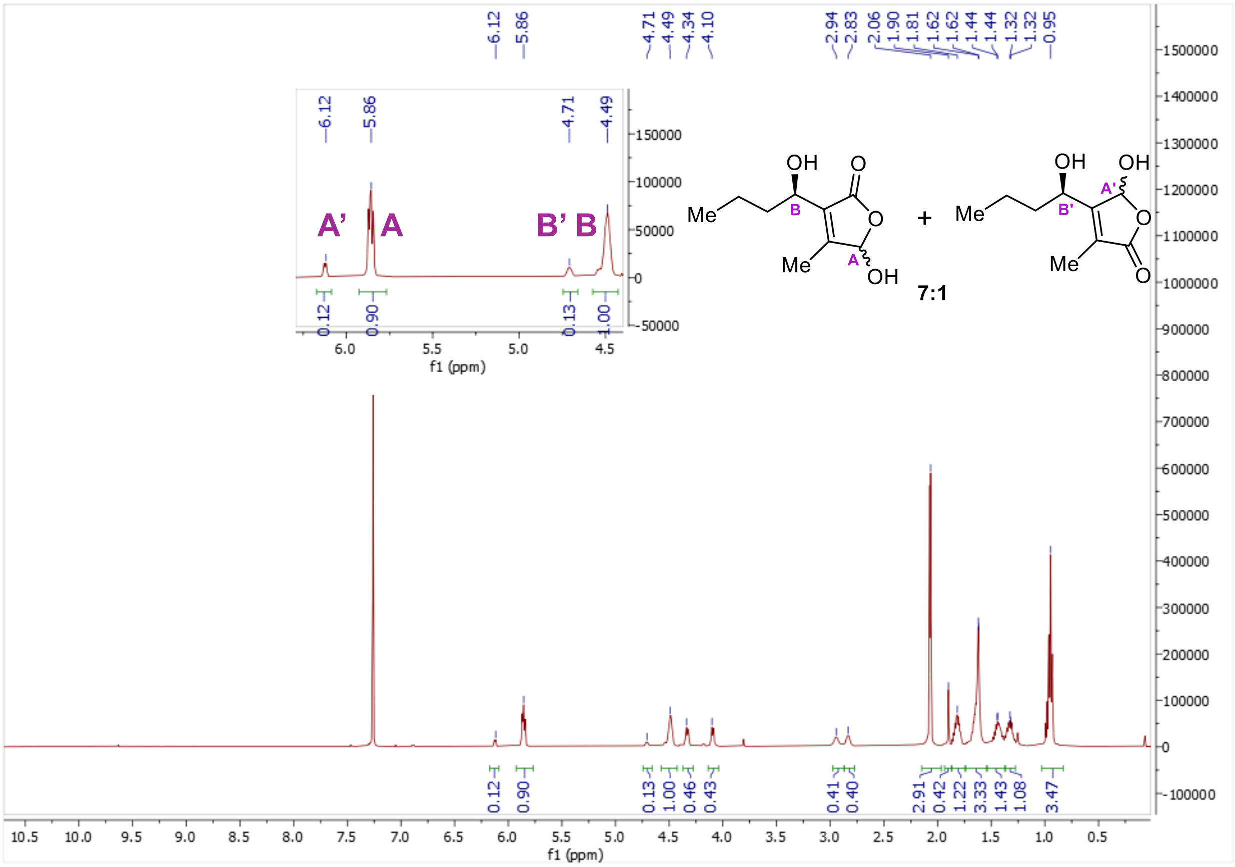

$^{13}$C for ***R*-SAB1 (*R*-5a)**

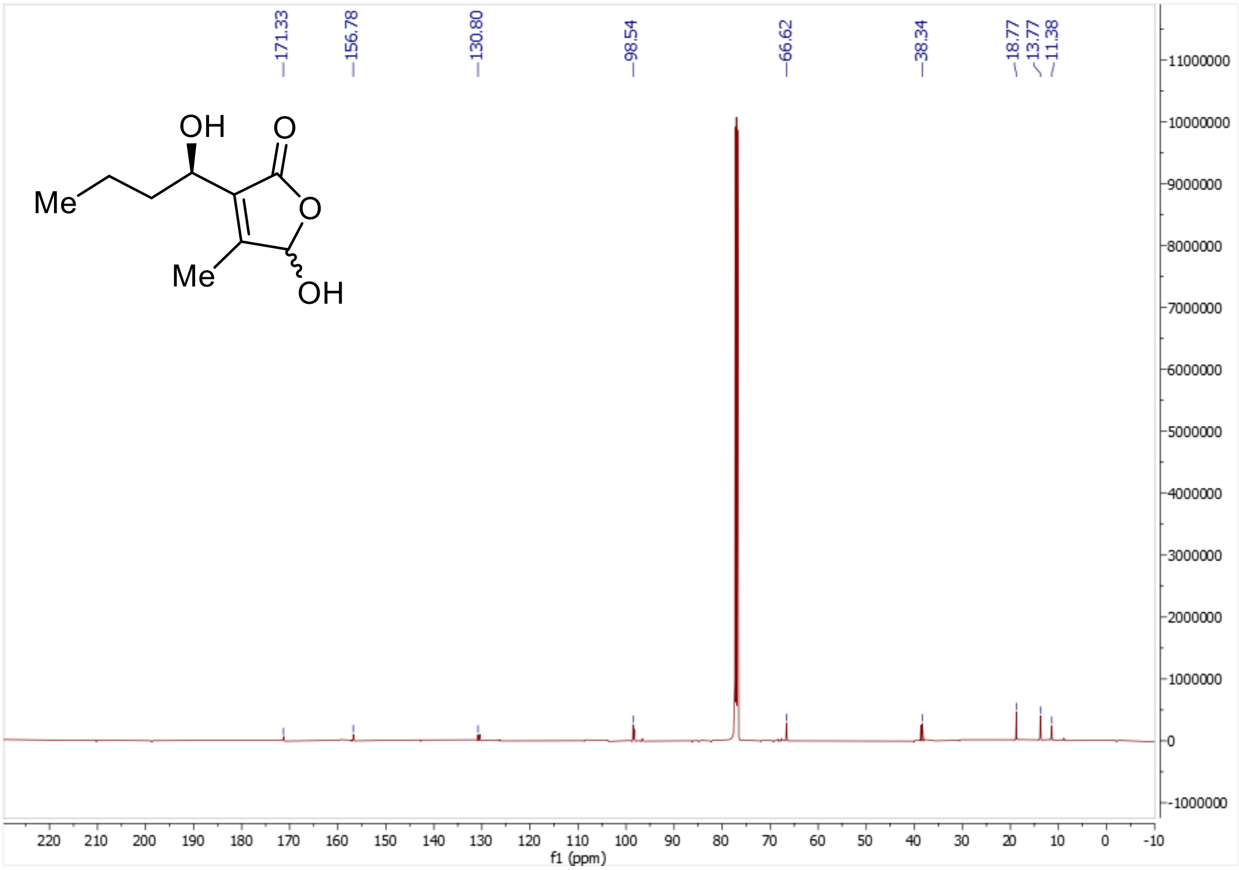

$^{1}$H for ***S*-SAB1 (*S*-5a)**

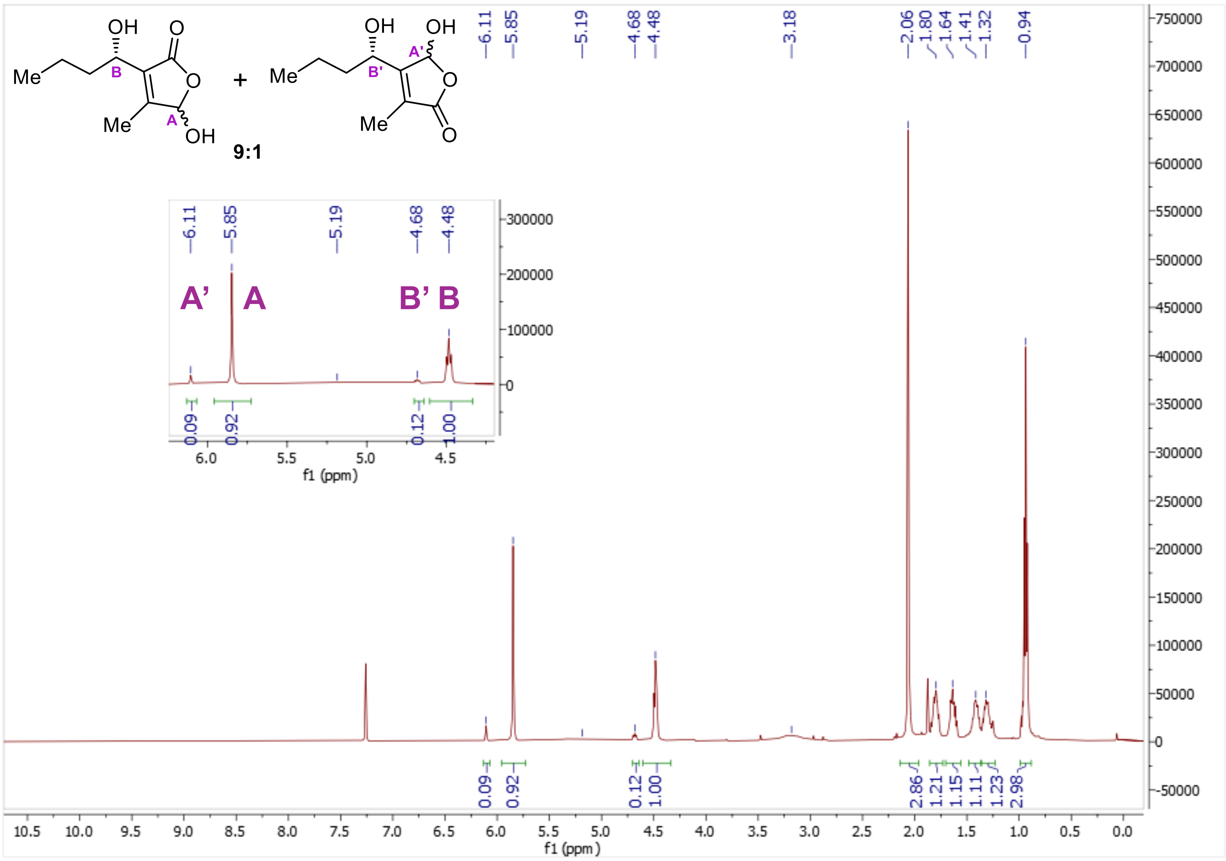

$^{13}$C for ***S*-SAB1 (*S*-5a)**

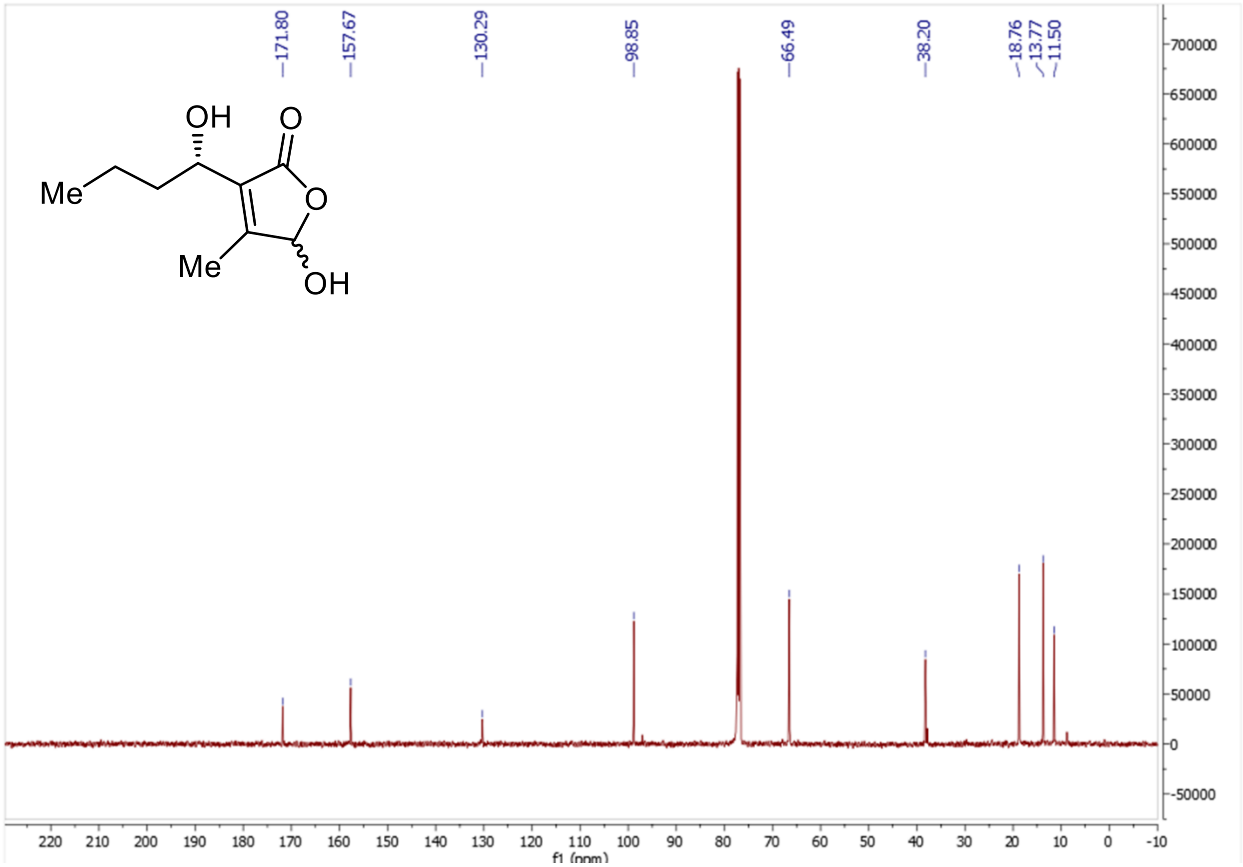

**Chiral HPLC Data**

**Racemic 12**

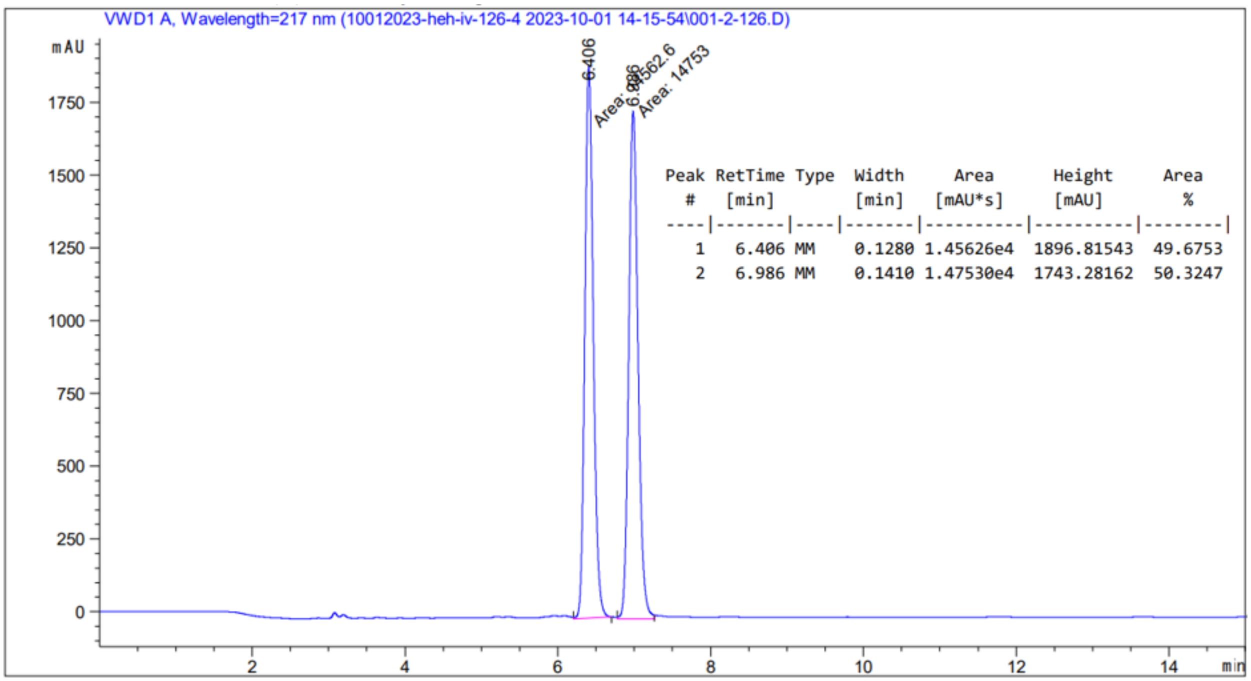

| Peak # | RetTime [min] | Type | Width [min] | Area [mAU*s] | Height [mAU] | Area % |
|---|---|---|---|---|---|---|
| 1 | 6.406 | MM | 0.1280 | 1.45626e4 | 1896.81543 | 49.6753 |
| 2 | 6.986 | MM | 0.1410 | 1.47530e4 | 1743.28162 | 50.3247 |

***(R)*-12**: $t_R$ (major) = 6.6 min and $t_R$ (minor) = 7.1 min.

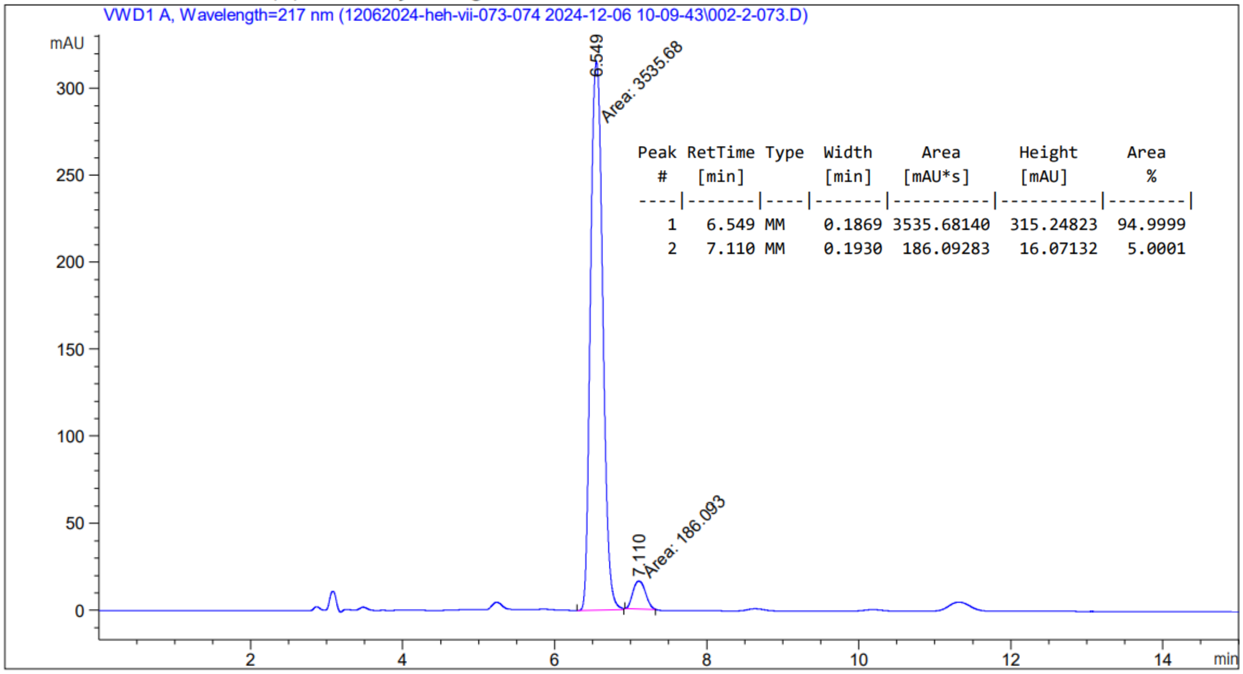

| Peak # | RetTime [min] | Type | Width [min] | Area [mAU*s] | Height [mAU] | Area % |
|---|---|---|---|---|---|---|
| 1 | 6.549 | MM | 0.1869 | 3535.68140 | 315.24823 | 94.9999 |
| 2 | 7.110 | MM | 0.1930 | 186.09283 | 16.07132 | 5.0001 |

(***S***)-**12** $t_R$ (major) = 7.1 min and $t_R$ (minor) = 6.6 min.

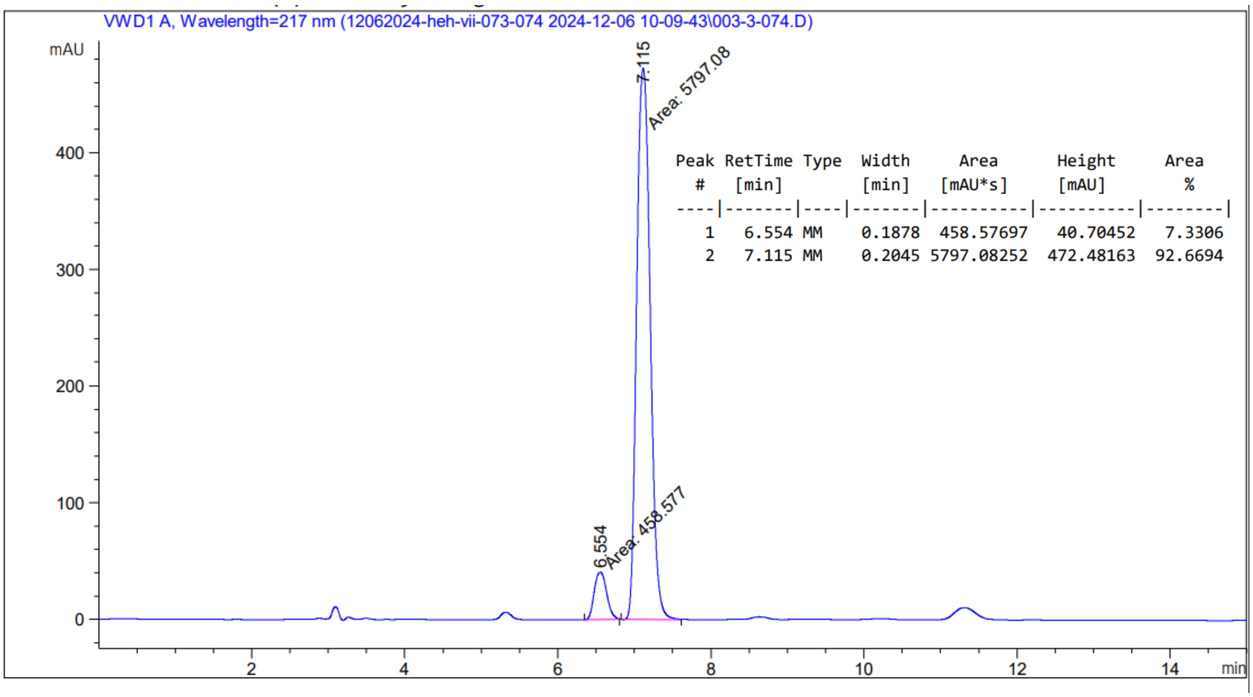

| Peak # | RetTime [min] | Type | Width [min] | Area [mAU*s] | Height [mAU] | Area % |
|---|---|---|---|---|---|---|
| 1 | 6.554 | MM | 0.1878 | 458.57697 | 40.70452 | 7.3306 |
| 2 | 7.115 | MM | 0.2045 | 5797.08252 | 472.48163 | 92.6694 |

17. Kapoor, I., Olivares, P. & Nair, S. K. Biochemical basis for the regulation of biosynthesis of antiparasitics by bacterial hormones. eLife 9, e57824 (2020).

26. Zhu, J., Chen, Z., Li, J. & Wen, Y. AvaR1, a butenolide-type autoregulator receptor in Streptomyces avermitilis, directly represses avenolide and avermectin biosynthesis and multiple physiological responses. Front. Microbiol. 8, (2017).
